# *RNAlysis*: analyze your RNA sequencing data without writing a single line of code

**DOI:** 10.1101/2022.11.25.517851

**Authors:** Guy Teichman, Dror Cohen, Or Ganon, Netta Dunsky, Shachar Shani, Hila Gingold, Oded Rechavi

**Author notes:** These authors contributed equally (Or Ganon, Netta Dunsky, Shachar Shani).

## Abstract

**Background:** Amongst the major challenges in next-generation sequencing experiments are exploratory data analysis, interpreting trends, identifying potential targets/candidates, and visualizing the results clearly and intuitively. These hurdles are further heightened for researchers who are not experienced in writing computer code, since the majority of available analysis tools require programming skills. Even for proficient computational biologists, an efficient and replicable system is warranted to generate standardized results.

**Results:** We have developed *RNAlysis*, a modular Python-based analysis software for RNA sequencing data. *RNAlysis* allows users to build customized analysis pipelines suiting their specific research questions, going all the way from raw FASTQ files, through exploratory data analysis and data visualization, clustering analysis, and gene-set enrichment analysis. *RNAlysis* provides a friendly graphical user interface, allowing researchers to analyze data without writing code. We demonstrate the use of *RNAlysis* by analyzing RNA data from different studies using *C. elegans* nematodes. We note that the software is equally applicable to data obtained from any organism.

**Conclusions:** *RNAlysis* is suitable for investigating a variety of biological questions, and allows researchers to more accurately and reproducibly run comprehensive bioinformatic analyses. It functions as a gateway into RNA sequencing analysis for less computer-savvy researchers, but can also help experienced bioinformaticians make their analyses more robust and efficient, as it offers diverse tools, scalability, automation, and standardization between analyses.

## Background

RNA sequencing continues to grow in popularity as an investigative tool for biologists. A huge variety of RNA-sequencing analysis methods allow researchers to compare gene expression levels between different biological specimens or experimental conditions, cluster genes based on their expression patterns, and characterize expression changes in genes involved in specific biological functions and pathways.

Specific tools exist to perform the different tasks described above (see **Discussion** and **Supplementary Table 1** for a detailed comparison of available tools). However, most analysis tools can only perform a subset of these tasks, and any out-of-the-ordinary research questions require researchers to write customized analysis scripts, which may not be easy to share or replicate. Moreover, many of the existing tools require users to be familiar with reading and writing code, making them usable only by researchers experienced in computer programming.

*RNAlysis* offers a solution to these problems by (1) using a modular approach, allowing users to either analyze their data step-by-step, or construct reproducible analysis pipelines from individual functions; and (2) providing an intuitive and flexible graphical user interface (GUI), allowing users to answer a wide variety of biological questions, whether they are general or highly specific, and explore their data interactively without writing a single line of code. *RNAlysis* includes thorough documentation and step-by-step guided analyses, to help new users to learn the software quickly and acquire good data analysis practices (available online, and also available as **Supplementary File 1**).

### Implementation

*RNAlysis* was designed to perform three major tasks: (1) pre-processing and exploratory data analysis; (2) finding gene sets of interest through filtering, clustering, and set operations; (3) visualizing intersections between gene sets and performing enrichment analysis on those sets (**Figure 1**).

**Figure 1:**
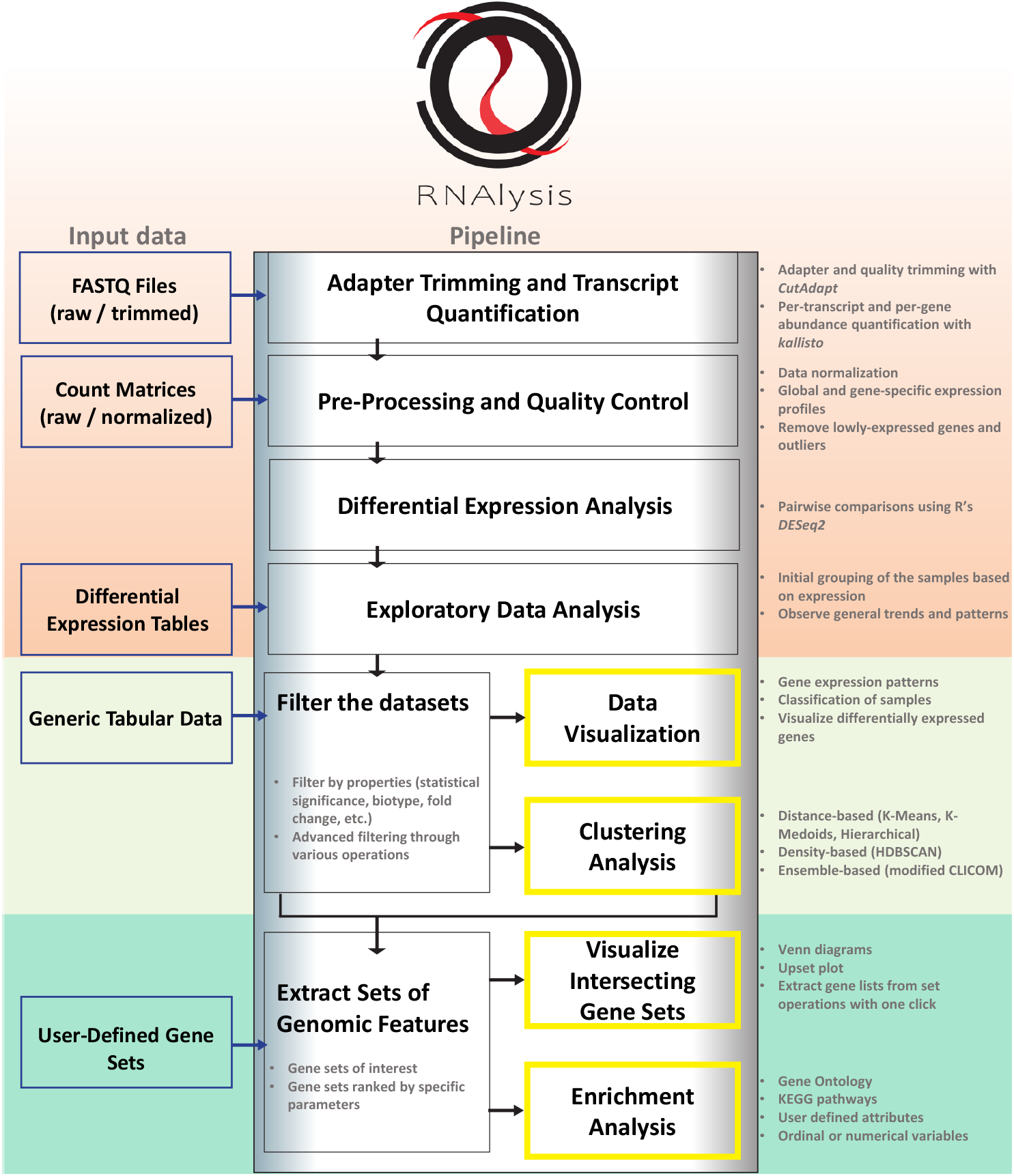
The workflow of *RNAlysis*. **Top section:** a typical analysis with *RNAlysis* can start at any stage from raw/trimmed FASTQ files, through more processed data tables such as count matrices, differential expression tables, or any form of tabular data. **Middle section:** data tables can be filtered, normalized, and transformed with a wide variety of filtering functions, allowing users to clean up their data, fine-tune their analysis to their biological questions, or prepare the data for downstream analysis. *RNAlysis* also provides users with a broad assortment of customizable clustering methods, to help recognize genes with similar expression patterns, and visualization methods to aid in data exploration. All of these functions can be arranged into customized Pipelines that can be applied to multiple tables in one click, or exported and shared with others. **Bottom section:** Once users have focused their data tables into gene sets of interest, or imported such gene sets from another source, they can use *RNAlysis* to visualize the intersection between different gene sets, extract lists of genes from any set operations applied to their gene sets and data tables, and perform enrichment analysis for their gene sets, using either public datasets such as GO and KEGG or customized, user-defined enrichment attributes.

### Input

*RNAlysis* can interface with existing tools, such as CutAdapt, *kallisto*, and DESeq2, (Bray et al., 2016; Love et al., 2014; Martin, 2011; Soneson et al., 2015) to enable users to run basic adapter-trimming, RNA sequencing quantification, and differential expression analysis through a graphical user interface. That is to say, users can begin their analysis with *RNAlysis* with sequencing data at any stage. Alternatively, users can load into *RNAlysis* data tables that were generated elsewhere. *RNAlysis* has a tabbed interface, which allows users to examine and analyze multiple data tables in parallel, seamlessly switching between them.

*RNAlysis* can accept data from any organism. *RNAlysis* can analyze gene expression matrices (raw or normalized), differential expression tables, or user-defined gene sets of interest. Moreover, *RNAlysis* accepts annotations for user-defined attributes of genes. Since *RNAlysis* works with tabular data, *RNAlysis* is applicable to any type of data table.

### Data validation and pre-processing

First, *RNAlysis* allows users to validate their data by summarizing and visualizing the data’s patterns and distribution. For instance, users may compare the distribution of gene expression between samples through scatter plots and pair plots, examining general trends in the data, as well as potential batch effects via clustergram plots and PCA projections.

Moreover, *RNAlysis* allows users to pre-process their data by normalizing it through one of various methods (such as Median of Ratios, Relative Log Ratio, Trimmed Mean of M-values, and more) (Anders and Huber, 2010; Bullard et al., 2010; Love et al., 2014; Maza et al., 2013; Robinson and Oshlack, 2010), filtering out lowly-expressed genes, and eliminating rows with missing data from their tables.

### Data filtering and clustering

After data pre-processing, users can further filter their data tables according to a broad array of parameters, depending on the nature of their data and biological questions. These filtering functions can be applied in particular orders and combinations to suit the user’s specific needs. These functions include, among many others, filtering by statistical significance or the direction and magnitude of fold change, filtering genomic features by their type, performing set operations between different data tables and gene sets (for instance - intersections, differences, majority vote intersections, etc.) between tables, etc.

One of the powerful features of *RNAlysis* is the ability to easily extract gene lists ifrom set operations applied to the user’s tables and gene sets, and use these lists in downstream analyses. This can be done either by applying a pre-defined set operation (like intersection or difference), or hand-picking subsets of interest through an interactive graphical platform.

Finally, *RNAlysis* allows users to cluster genes based on the similarity of their expression patterns. *RNAlysis* supports an extensive selection of clustering algorithms, including distance-based clustering (K-Means, K-Medoids, Hierarchical clustering), density-based clustering (HDBSCAN) (McInnes et al., 2017), and ensemble-based clustering (a modified version of the CLICOM algorithm) (Mimaroglu and Yagci, 2012).

Moreover, *RNAlysis* provides users with a wide array of distance metrics for clustering analysis. This includes the implementation of distance metrics that were specially developed for biological applications such as time-course gene expression data (Son and Baek, 2008), and distance metrics that were empirically found to best suit transcriptomics analysis (Jaskowiak et al., 2014).

### Modularity and building customized pipelines

Filtered data tables can be saved or loaded at any stage during the analysis. The operations performed on the data, as well as their order, will automatically be reflected in the output files’ names. Additionally, any operation applied to the data can be undone with a single click, and *RNAlysis* displays the history of commands applied to each table in the order they were applied.

As mentioned earlier, users can ‘bundle’ any of the functions *RNAlysis* offers into distinct Pipelines, which can then be applied in the same order and with the same parameters to any number of similar data tables. This helps users to save time and avoid mistakes and inconsistencies when analyzing a large number of datasets. Pipelines can also be exported and shared with other researchers, who can then use these Pipelines on any machine that installed *RNAlysis*. This feature makes analysis pipelines easier to report and share, increasing the reproducibility and transparency of bioinformatic results.

### Enrichment analysis

After applying the aforementioned analyses to summarize data tables down to gene sets of interest, users can carry out enrichment analysis for those gene sets via the Enrichment window. Gene set enrichment analysis is a collection of methods for identifying classes of genes, biological processes, or pathways, that are over- or under-represented in a gene set of interest (Subramanian et al., 2005). Enrichment analysis is very popular in the context of RNA sequencing analysis, since it allows researchers to associate a differentially-expressed gene set with underlying biological functions (Ashburner et al., 2000; Kanehisa and Goto, 2000).

*RNAlysis* supports multiple approaches and statistical methods for enrichment analysis, including classic gene-set enrichment analysis, permutation tests (Phipson and Smyth, 2010), background-free enrichment analysis (Eden et al., 2007; Wagner, 2017), and enrichment for ordinal or continuous variables.

*RNAlysis* can automatically retrieve enrichment analysis annotations of all major model organisms from widely accepted databases such as Gene Ontology categories and KEGG pathways (Carbon et al., 2021; Kanehisa et al., 2022). However, unlike most other analysis pipelines, *RNAlysis* also accepts annotations for user-defined attributes and groups (see **Supplementary Table 1**). This allows users to tailor their analyses to their specific needs and biological questions.

### Documentation and accessibility

While *RNAlysis* can be operated entirely within a graphical interface, all the functions and features *RNAlysis* offers can also be imported and used in standard Python scripts, allowing users with coding experience to further automate and customize their bioinformatic analyses.

*RNAlysis* includes extensive documentation to guide new and returning users. A User Guide offers a bird’s eye view of the modules and features of *RNAlysis*, along with video demonstrations, usage examples, and recommended practices. The remainder of the documentation provides a complete reference of the functions and features available in *RNAlysis*, where users can look up specific entries to get a more thorough review of their theoretical background, use cases, and optional parameters.

The project is available as an open-source, public GitHub repository. A multitude of test cases are also provided within the package, which are executed automatically every time the source code is updated, ensuring that data analysis with *RNAlysis* remains consistent and reliable.

*RNAlysis* is powered by various open-source projects (Harris et al., 2020; Heyer et al., 1999; Hunter, 2007; Lam et al., 2015; Lex et al., 2014; Mckinney, 2010; Pedregosa et al., 2011; Seabold and Perktold, 2010; Son and Baek, 2008; Virtanen et al., 2020; Waskom, 2021) which are installed automatically and used when needed.

## Results

We examined the ability of *RNAlysis* to facilitate analyses of multiple different publicly available datasets (Davis et al., 2022; Dodd et al., 2018; Finger et al., 2019; Schreiner et al., 2019). First, we analyzed time-series gene expression data, using clustering analysis to group genes based on their expression pattern. Then, we demonstrated the analysis of multiple RNA sequencing datasets from raw FASTQ files and showing the applications of analysis Pipelines and set operations between datasets. A step-by-step tutorial of these analyses is available in the online *RNAlysis* documentation (also available as **Supplementary File 1**).

### Analysis #1: Exploring gene expression patterns across the development of *Caenorhabditis elegans*

In the first analysis, we examined a dataset describing average gene expression under different developmental stages of *Caenorhabditis elegans* nematodes, derived from the control samples of many publicly available RNA sequencing experiments (Davis et al., 2022).

Exploratory data analysis revealed that the different developmental stages show the highest correlations with contiguous developmental stages (**Figure 2A**), and PCA uncovered a semi-circular pattern, with over 75% of the data’s variance explained by the first two principal components (**Figure 2B**). Interestingly, the first Principal Component appears to arrange the samples by their relative germline content, with embryos and adult nematodes on one end, L1-L3 larvae on the other end, and L4 larvae in between. The second Principal Component appears to arrange the samples by their developmental stage, with embryos appearing at the top of the graph and adults at the bottom.

**Figure 2:**
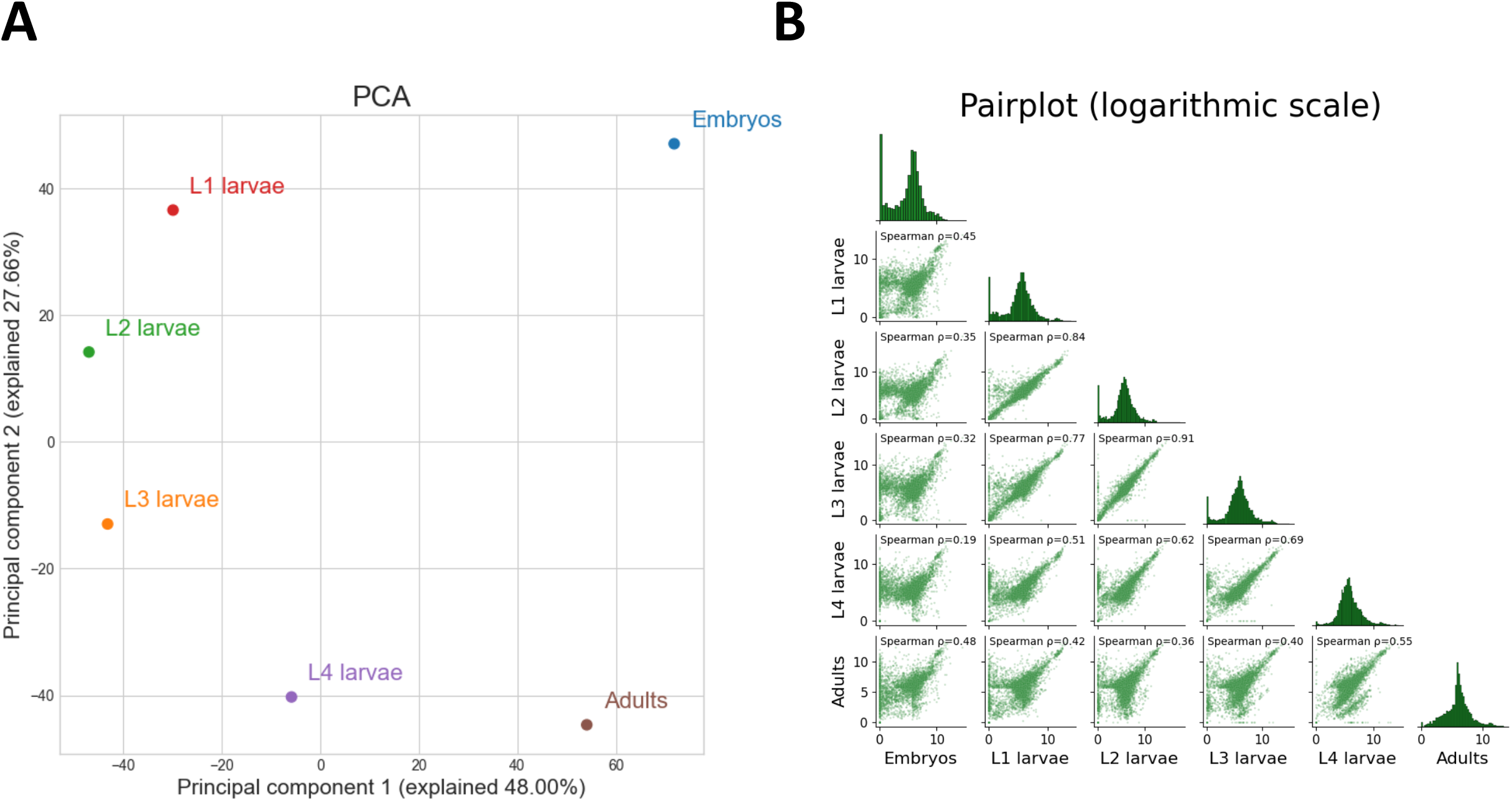
Exploratory data analysis reveals patterns in time-series gene expression data. (A) Principal Component Analysis projection of the time series data. Depicted are the first two principal components, explaining >75% of the variance in the data. Data was power-transformed and regularized before the analysis. (B) Pair-plot, depicting the pairwise Spearman correlation between each pair of samples, and a histogram of normalized gene expression in each sample. Each dot represents the log of normalized expression of a single gene.

Next, we extracted clustering results at three different resolutions by using exemplars from three different classes of clustering algorithms: a distance-based algorithm (K-Medoids) (**Supplementary Figure 1**), a density-based algorithm (HDBSCAN) (**Supplementary Figure 2**), and an ensemble-based algorithm (CLICOM) (**Figure 3**). While one of the most challenging aspects of RNA sequencing clustering analysis is the requirement to specify in advance the number of clusters, *RNAlysis* provides unbiased clustering methods that can either estimate a good number of clusters to detect, or require no such input at all – instead specifying the smallest cluster size that would be of interest to the user.

**Figure 3:**
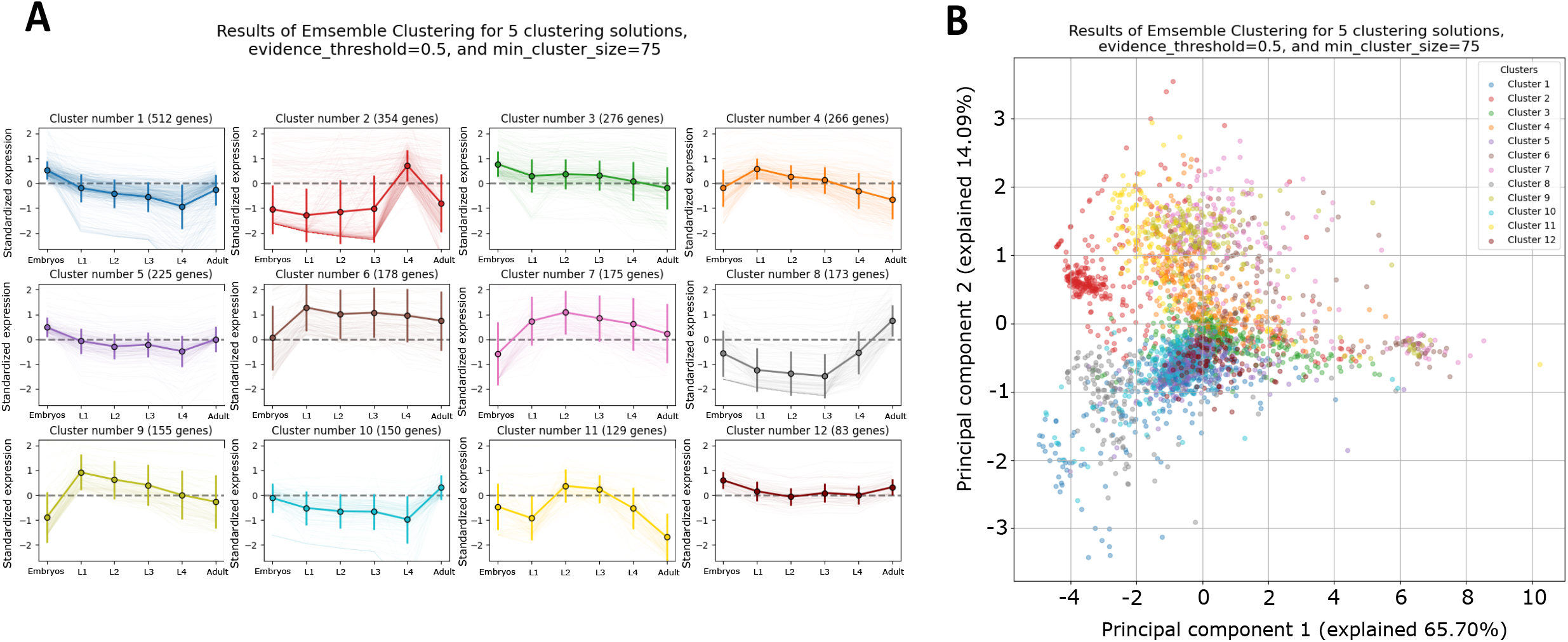
Clustering analysis of time-series gene expression data. (A) Clustering analysis of the data using modified CLICOM clustering, using five underlying clustering setups, evidence threshold of 50%, and a minimal cluster size of 75 (Mimaroglu and Yagci, 2012). Clusters are sorted by their size. Each graph depicts the power-transformed and regularized expression of all genes in the cluster, with the center lines denoting the clusters’ means and standard deviations across developmental stages of *C. elegans* nematodes. (B) PCA projection of the power-transformed and regularized gene expression data. Each dot represents a gene. The points are colored according to the cluster they belong in the CLICOM clustering result depicted in (C) above.

While the data examined here only contains a single sample for each experimental condition, *RNAlysis* is well suited for clustering analysis of replicate data, since it’s able to cluster each batch of replicates separately and combine the results of those batches, resulting in more accurate and robust clustering results (Sloutsky et al.).

Finally, we plotted the expression level of specific genes of interest under the different developmental stages (**Figure 4A**) and performed GO enrichment on one of the clusters we previously detected, revealing a strong enrichment for neuropeptide signaling pathways (**Figure 4B**).

**Figure 4:**
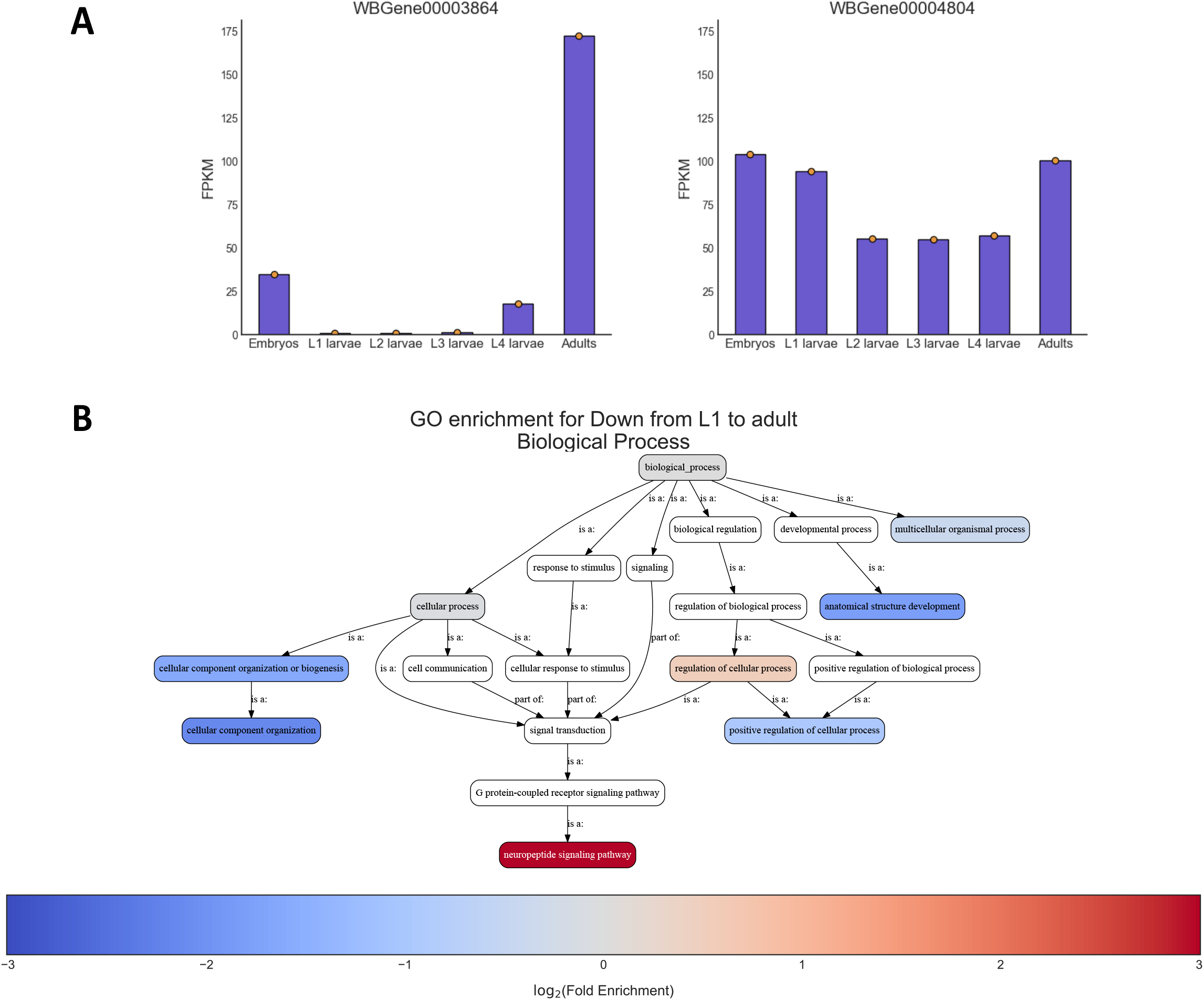
Gene expression plots and enrichment analysis of time-series gene expression data. (A) Normalized gene expression values of the normalized time series data for two sample genes *oma-1* (WBGene00003664) and *skn-1* (WBGene00004804). (B) GO enrichment ontology graph, depicting enrichment results for cluster #9 (**Figure 3**). The graph depicts the hierarchical relationship between the GO terms. Each GO term was colored according to its log2 Fold Enrichment score if it was statistically significant (q-value ≤ 0.05).

### Analysis #2: Measuring the effect of stress on the expression of small RNA factors

In the second analysis we demonstrate here, we analyzed three datasets that examined the effects of three different stress conditions (osmotic stress, heat shock, and starvation) on gene expression (Dodd et al., 2018; Finger et al., 2019; Schreiner et al., 2019). This is a replication of a previously published analysis (Houri-Zeevi et al., 2021) done with an earlier version of *RNAlysis* (version 1.3.5, 2019), where the purpose was to examine the effects of stress exposure on the expression of small RNA factors. This analysis shows how *RNAlysis* facilitates intuitively answering highly specific biological questions.

We started the analysis with raw FASTQ files, applying adapter trimming, transcript expression quantification, and differential expression analysis to the three datasets, all executed through the *RNAlysis* graphic interface.

Next, we examined the distribution of differentially expressed genes under each condition with a Volcano Plot (**Figure 5A**) and extracted from each differential expression table the lists of significantly up-regulated and down-regulated genes. This step was automated by building and applying a Pipeline, allowing us to analyze all three tables in the exact same manner, with the click of a button.

**Figure 5:**
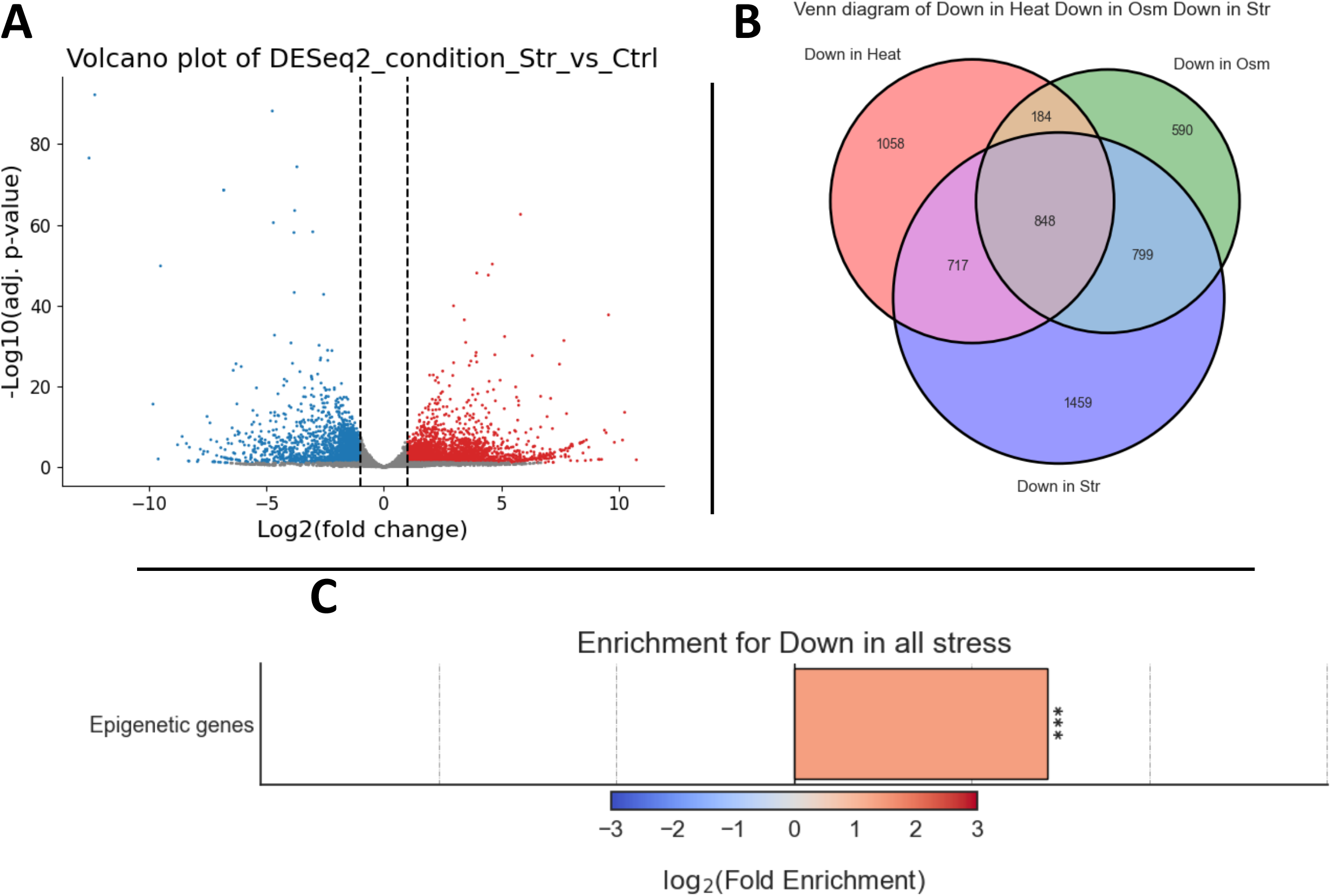
Analysis of stress-induced gene expression changes. (A) Volcano plot depicting differential expression results, comparing worms that experienced starvation to worms that grew under normal conditions. Each dot represents a gene. Differentially expressed genes with log2 fold change ≥ 1 were painted in red, and differentially expressed genes with log2 fold change ≤ −1 were painted in blue. (B) A proportional Venn Diagram depicting the intersection between genes that are significantly downregulated under heat shock, osmotic stress, or starvation, compared to their matching control samples. (C) Log2 X-fold enrichment score for a curated list of epigenetic genes, in the set of genes significantly downregulated under all stress conditions. The p-value for enrichment was calculated using 10,000 random gene sets identical in size to the tested group. *** indicates p-value ≤ 0.001.

Following these filtering steps, we examined the intersection of the up- and down-regulated genes between the different stress conditions (**Figure 5B**) and extracted the list of genes that are significantly up/down-regulated under all stress conditions. We then created an appropriate background set for enrichment analysis by calculating the union gene lists of all genes which are sufficiently expressed under at least one stress condition.

Finally, we ran enrichment analysis on the stress-downregulated genes, measuring whether they are significantly enriched for a user-defined list of epigenetic-related genes (**Figure 5C**). We found the stress-downregulated genes to be significantly enriched for epigenetic-related genes, as previously shown (Houri-Zeevi et al., 2021). Enrichment for user-defined attributes is a feature unique to *RNAlysis*, allowing users to answer highly specific biological questions. This means that the users are not limited to widely available datasets, but can directly analyze any gene sets and attributes of interest without the need to write any code.

## Discussion

*RNAlysis* offers researchers a robust, scalable, and easy-to-use tool to analyze RNA sequencing data. *RNAlysis* was designed not only to be intuitive and approachable for new users, but also to provide a high degree of efficiency, control, and robustness to experienced bioinformaticians.

Other useful software tools for the analysis of RNA sequencing data exist (see **Supplementary Table 1**). For example, *Galaxy* (Afgan et al., 2018) is a web-based scientific analysis platform for the analysis of biological data. *Galaxy* offers many shared features with *RNAlysis*, including integration of existing analysis tools, extensive documentation, and the ability to filter, sort, and intersect data tables. However, contrary to *Galaxy, RNAlysis* aims to simplify commonly used actions, such as filtering and set operations, by providing users with dozens of ready-made filtering functions relevant to RNA sequencing data, and supporting set operations on an arbitrary number of datasets with an intuitive, point-and-click interface. Moreover, *RNAlysis* offers analysis methods that are particularly useful to RNA sequencing data, such as advanced clustering methods, and enrichment analysis for user-defined attributes.

Tools such as *ARPIR* (Spinozzi et al., 2020) and *NetSeekR* (Srivastava et al., 2022), can take users all the way from the alignment of reads and differential expression analysis through GO enrichment and other tertiary analyses such as gene network analysis. Other tools like *ideal* (Marini et al., 2020), *PIVOT* (Zhu et al., 2018), and *DEBrowser* (Kucukural et al., 2019) provide users with a graphical interface to perform differential expression analysis and enrichment analysis.

While these tools allow less experienced bioinformaticians to perform basic transcriptomic analysis, they are limited in their capability to filter datasets, perform set operations between datasets, use more sophisticated clustering algorithms, or automate and streamline data analysis with pipelines. In contrast to these tools, *RNAlysis* is highly modular and customizable, allowing users to tailor their analyses to their biological questions through advanced data filtering, intersecting multiple datasets, and a high degree of control over analysis parameters at every stage of the process. Moreover, many of these tools cannot analyze RNA sequencing experiments from start to finish, since they do not support pre-processing, alignment, or quantification utilities of FASTQ files.

## Conclusion

*RNAlysis* offers a modular toolbox for RNA sequencing data analysis, with the unique combination of an intuitive graphical interface and highly customizable analysis workflows, setting it apart from most other RNA sequencing analysis tools.

We believe that the ability to build customized and reproducible analysis pipelines, combined with the user-friendly interface, will allow researchers to easily gain novel biological insights from RNA sequencing data.

## Acknowledgments

We thank all members of the Rechavi lab for fruitful discussions, feedback, and support. O.R. is grateful to funding from the Eric and Wendy Schmidt Fund for Strategic Innovation (Polymath Award #0140001000) and the generous support from the Morris Kahn foundation. GT is grateful to the Milner Foundation. The Rechavi lab is funded by ERC grant #335624 and the Israel Science Foundation (grant#1339/17).

## Abbreviations

GUI: Graphical user interface
GO: Gene Ontology
PCA: Principal component analysis
RNA: Seq: RNA sequencing
MRN: Median ratio normalization
TMM: Trimmed Mean of M-values
RLE: Relative Log Expression
WT: Wild type
KEGG: Kyoto Encyclopedia of Genes and Genomes

## Availability and requirements

**Project name**: *RNAlysis*

**Project home page**: https://github.com/GuyTeichman/RNAlysis

**Operating system(s)**: Platform independent

**Programming language**: Python 3

**Other requirements:** Python 3.7.9 or higher, GraphViz 3.0 or higher (optional), *kallisto* 0.44.0 or higher (optional), R 4.1.0 or higher (optional), Microsoft C++ Build Tools 14.0 or higher (optional, on Windows computers only)

**License**: MIT

**Any restrictions to use by non-academics**: none

---

**Supplementary Figure 1:**

**K-Medoids Clustering analysis of time-series gene expression data**

Clustering analysis of the data using K-Medoids clustering, after selecting an appropriate number of clusters (K=11) using the Gap Statistic method (Tibshirani et al.). Clusters are sorted by their size. Each graph depicts the power-transformed and regularized expression of all genes in the cluster, with the center lines denoting the clusters’ Medoids and standard deviations across developmental stages of *C. elegans* nematodes.

**Supplementary Figure 2:**

**HDBSCAN Clustering analysis of time-series gene expression data**

Clustering analysis of the data using HDBSCAN clustering, with a minimal cluster size of 75 (McInnes et al., 2017). Clusters are sorted by their size. Each graph depicts the power-transformed and regularized expression of all genes in the cluster, with the center lines denoting the clusters’ means and standard deviations across developmental stages of *C. elegans* nematodes.

## Notes

### Competing Interest Statement

The authors have declared no competing interest.

