## Supplementary File 1 for "*RNAlysis*: analyze your RNA sequencing data without writing a single line of code"

### A-to-Z tutorial - example analysis

#### Intro and recommendations

In this A-to-Z tutorial, we are going to analyze two datasets step-by-step, with each analysis covering different aspects of using *RNAlysis*.

These analyses both focus on RNA sequencing data from nematodes, but the same principles can be applied to data from any organism.

#### Analysis #1 - time series RNA sequencing

##### The dataset

For the first analysis, we will use a publicly available dataset from WormBase. This dataset describes the median expression level of each gene over the developmental stages of *C. elegans* nematodes, averaged from RNA-sequencing short-read experiments that have been identified in the literature as being 'controls'.

The original table linked above contains the mean expression, median expression, and number of samples for each developmental stage. Since we are only interested in the medians, before starting the analysis, I copied the Median columns to a new table, and gave the columns more readable names. You can find the table I created here.

Let's start by loading the dataset - in the main window, click on the "Load" button and choose the table's file from your computer. We will then use the drop-down menu to change our table type from "Other" to "Count matrix". This will allow us to later on use analysis methods that are dedicated to count matrix-style datasets.

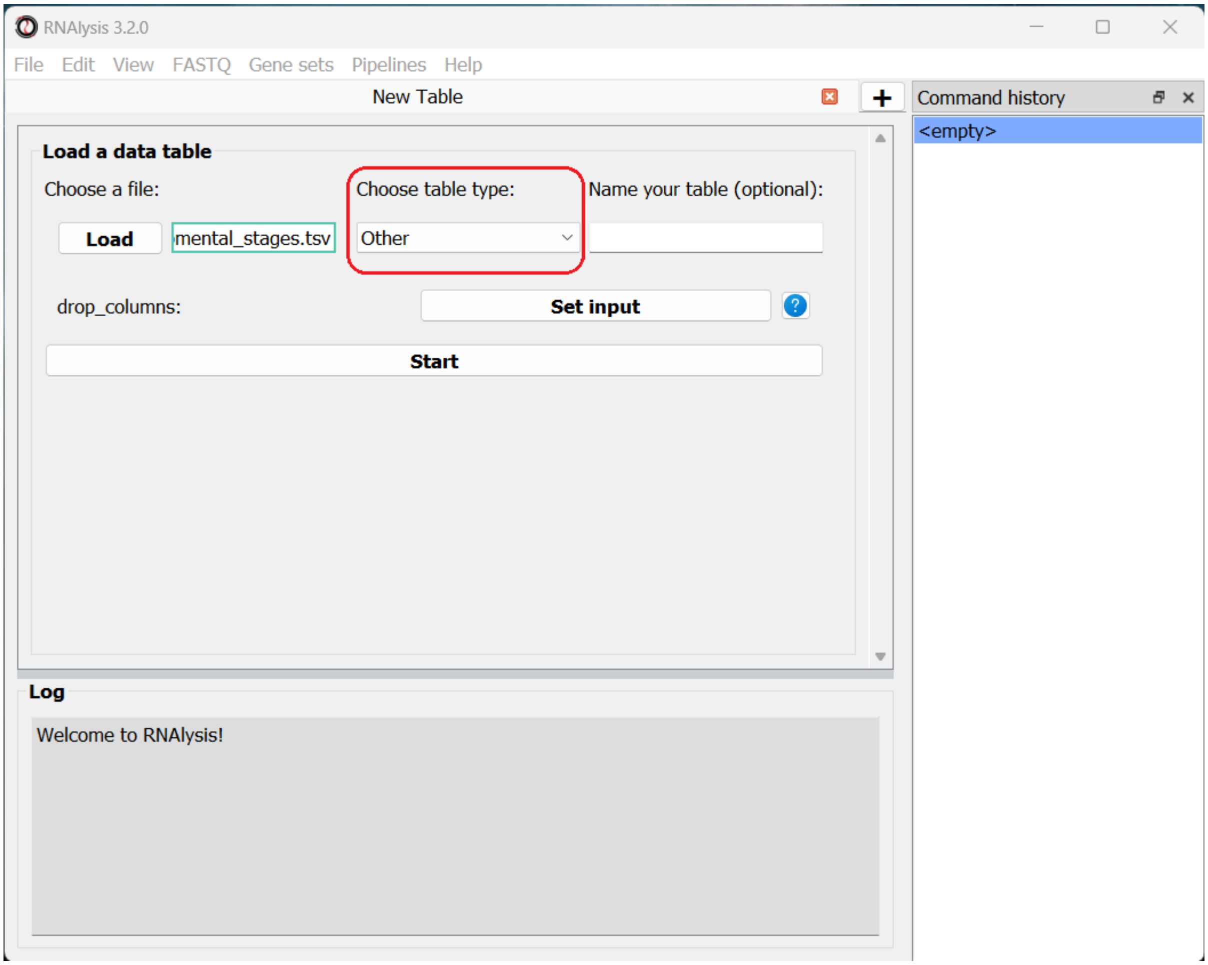


Once we picked a table type, a new option will appear, allowing us to specify whether our table was pre-normalized. We will set this option to “True”, since our dataset was already normalized to Fragments Per Kilobase Million (FPKM). This is not a mandatory step, but if we don’t do it, *RNAlysis* will warn us whenever we run analyses that expect normalized values.

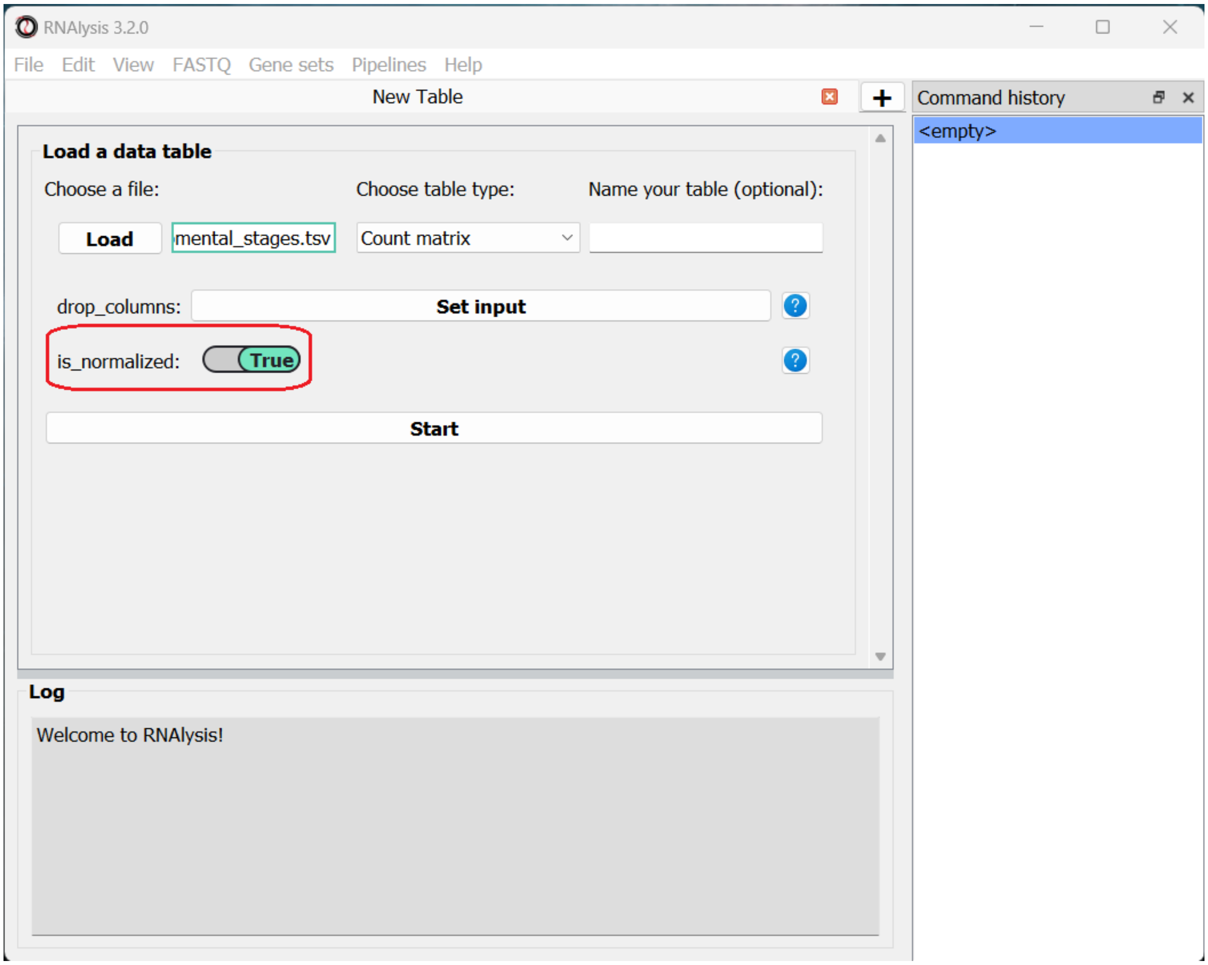


Finally, we can click the “start” button to actually open our table on *RNAlysis*. The window will now display a preview of our table, as well as a a short summary of our table’s content (table name, table type, number of rows and columns).

At the bottom of the window we can also see a log box - this is where *RNAlysis* will display notes and warnings regarding our analysis.

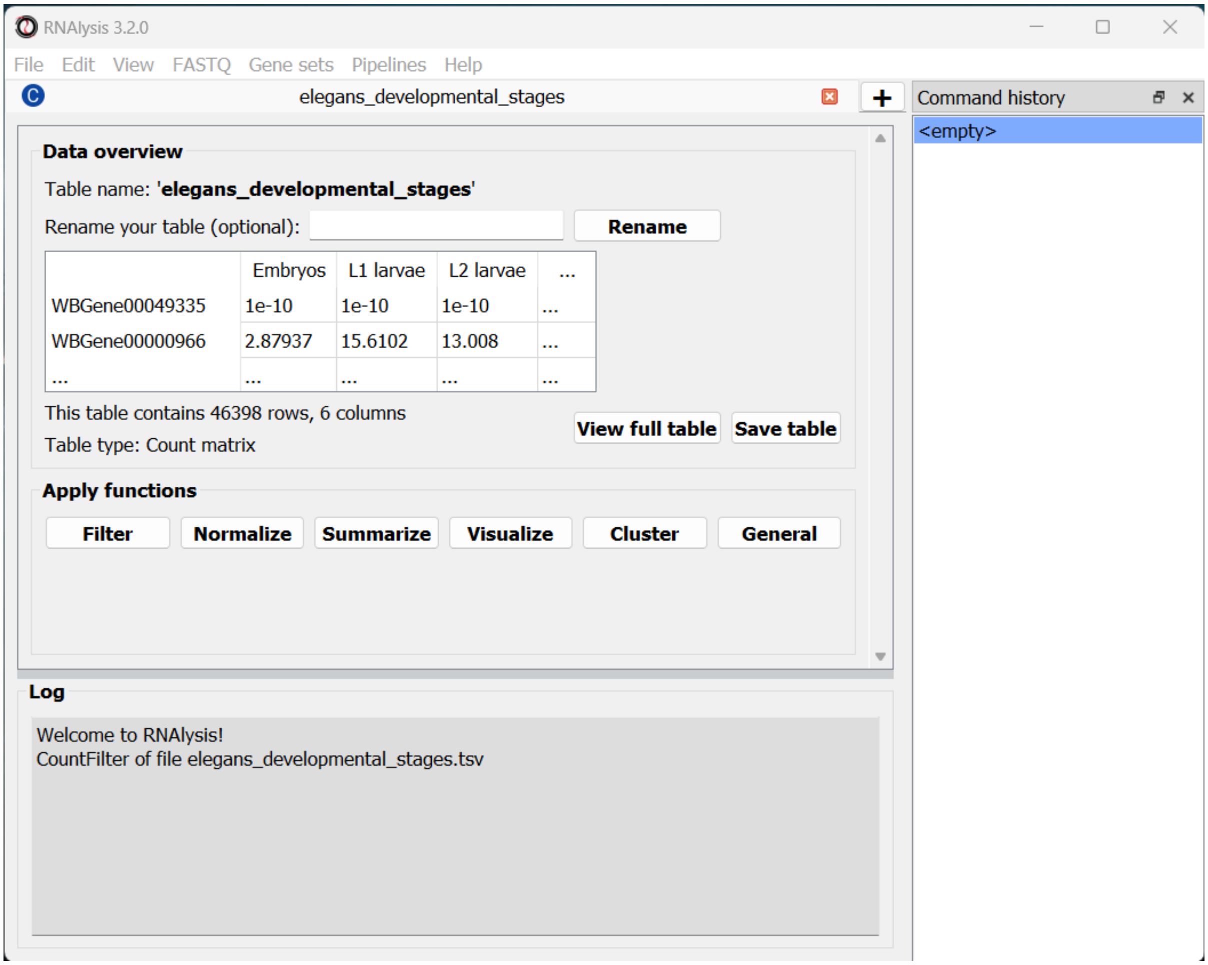


If we want to see the entire table, we can click on the “View full table” button, and the entire table will appear in a new window:

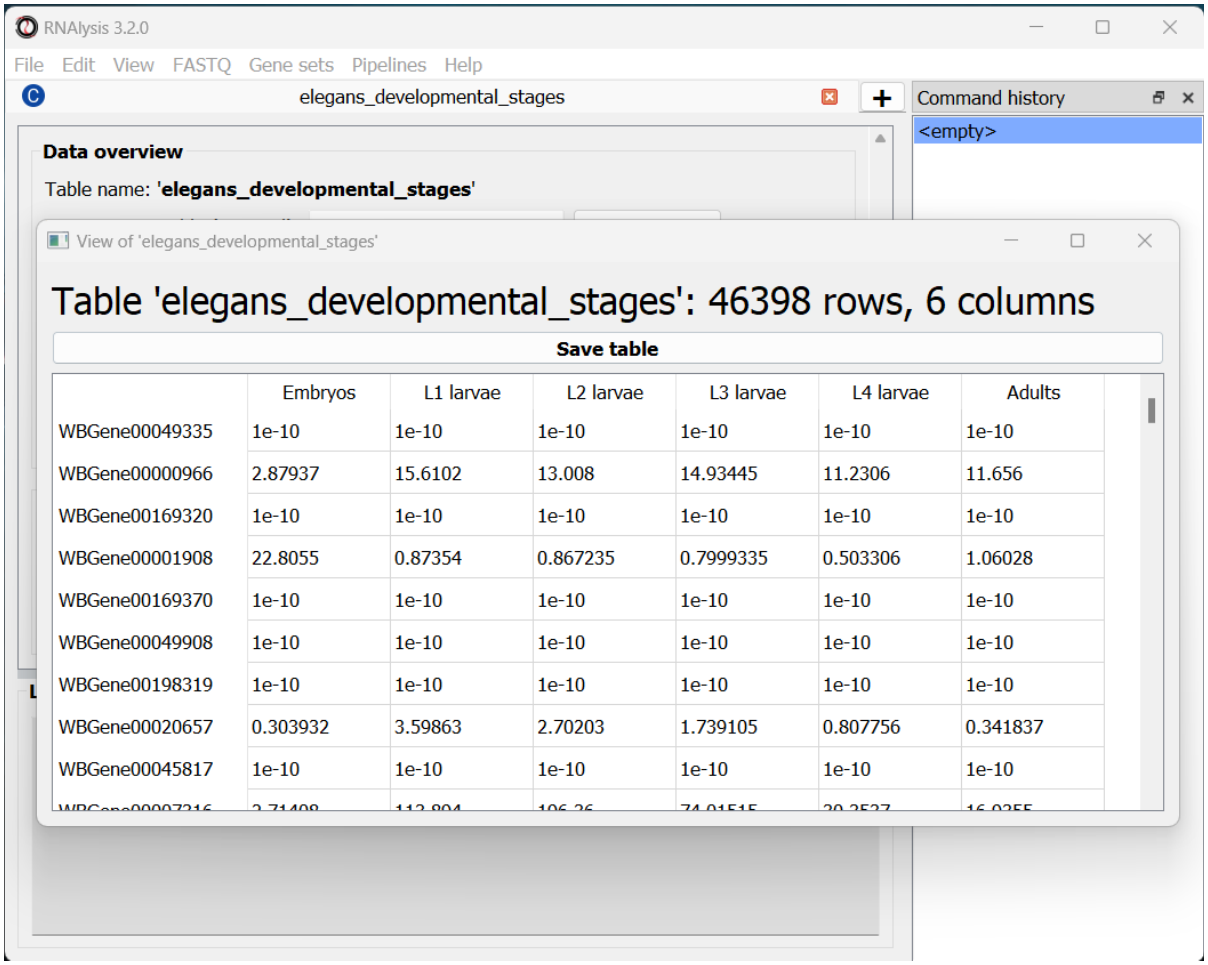


At any point during the analysis, we can click the "Save table" button to save our table into a new file.

#### Data preprocessing and exploratory data analysis

We can now begin exploring our data! Let's start with a pre-processing step - removing lowly-expressed genes.

##### Filter out lowly-expressed genes

We want to filter out the genes that have not been expressed or that have low expression across all samples. Lowly-expressed genes can negatively affect our analysis downstream, since the % error in the expression of these genes is relatively high, and these genes are likely to add noise rather than useful signal to our analysis.

In this particular example we also want to analyze a fairly small gruop of genes. This is because later down the line we will take our data through clustering analysis, and trying to cluster such a large group of genes could take your computer an extremely long time to finish. Therefore, we are going to filter our table, so that we keep only genes with 50 or more normalized reads in at least 1 experimental condition.

To apply a filtering function to our table, click on the “Filter” tab, and select a function from the drop-down menu that opens:

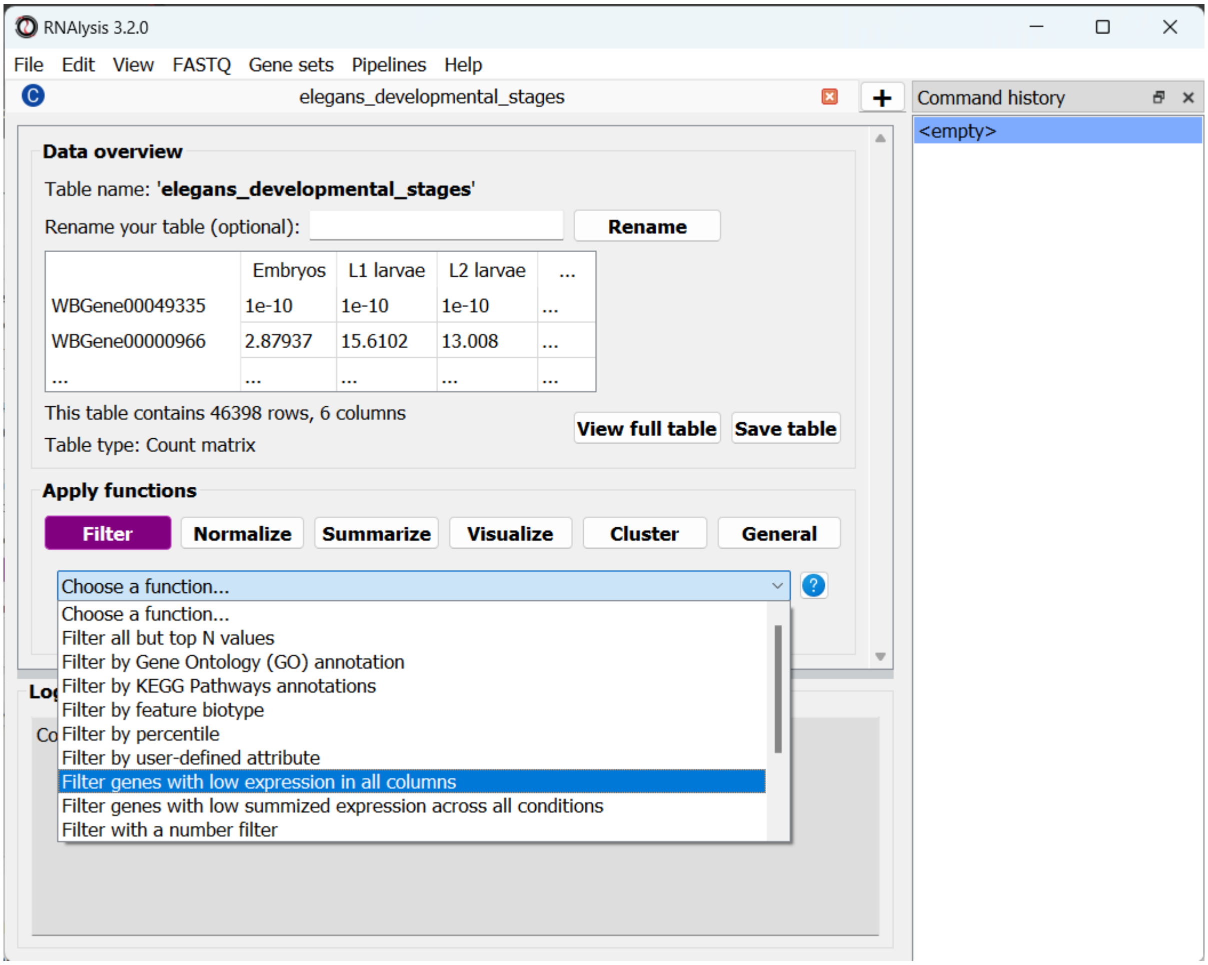


*RNAlysis* contains a wide diversity of filtering functions - filtering genes by expression level, fold change direction and magnitude, statistical significance, removing rows with missing data, and many more. We are going to select “Filter genes with low expression in all columns”. This function will filter our any gene whose expression level is lower than X in every single column. This means we only keep genes that are desirably expressed in at least one experimental condition. If you are not sure what a function does, you can click on the blue question mark button next to the function’s name to read a short description, or go to the function’s help page by clicking on the blue link at the bottom of the main window.

Once we choose a function, the window will expand, and the filtering function’s parameters will be displayed below. We can modify these parameters to choose exactly how to filter our table.

In this example, we are going to set the parameter *threshold* to 50 FPKM. This means that any genes which have less than 50 FPKM in **all** of the table’s columns will be filtered out. If you are not sure what a parameter does, you can hover your cursor over its name, or click on the blue question mark button next to the parameter’s name.

Every filtering function in *RNAlysis* supports two additional parameters: *opposite* and *inplace*.

*opposite*, as the name indicates, allows you to apply the **inverse** of your filtering operation to your table. For example, in our case, instead of filtering out lowly-expressed genes, we will filter out everything **but** the lowly-expressed genes.

*inplace* allows us to choose whether we want to apply the filtering operation in the same window (default), or to keep an unfiltered version of the table and apply the filtering operation in a new tab. This can be conveniet if we want to try multiple filtering approaches in parallel, or if we need to use the unfiltered table later down the line. Regardless of what we choose, the original file we loaded will not be changed by our filtering unless we expelicitly save our filtering results, and any filtering operation we apply can be undone with the click of a button.

Once we are happy with the parameters we chose, we can scroll down and click on the "Apply" button to apply the filtering function to our table.

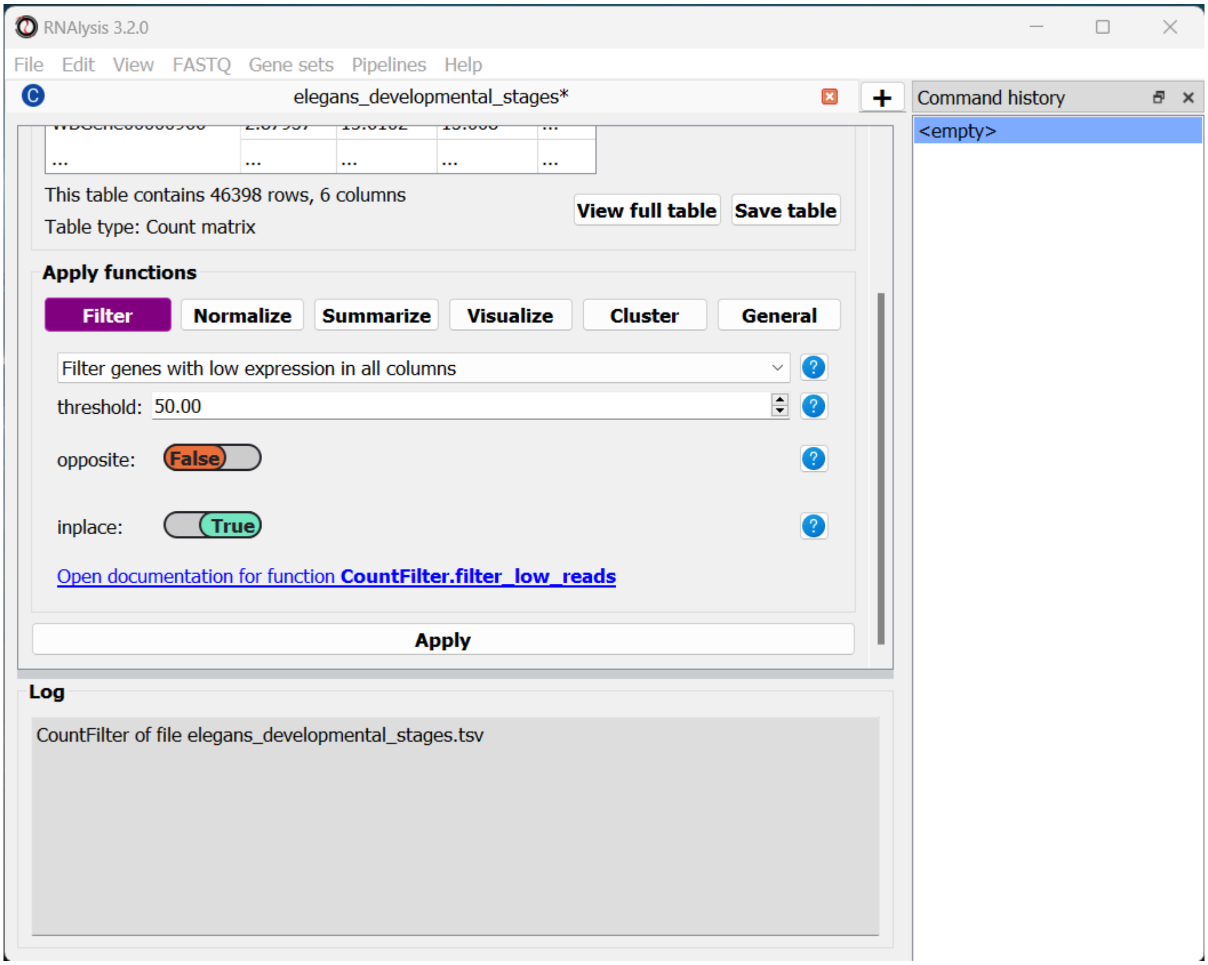


After clicking "Apply", we can see that our action has been added to the *command history* panel on the right of the window. We can undo and redo any operation we apply inplace by clicking on a specific row in this panel. Moreover, we can see that a short summary of our filtering action has printed to the log box at the bottom, and that the name and size of the table have been updated.

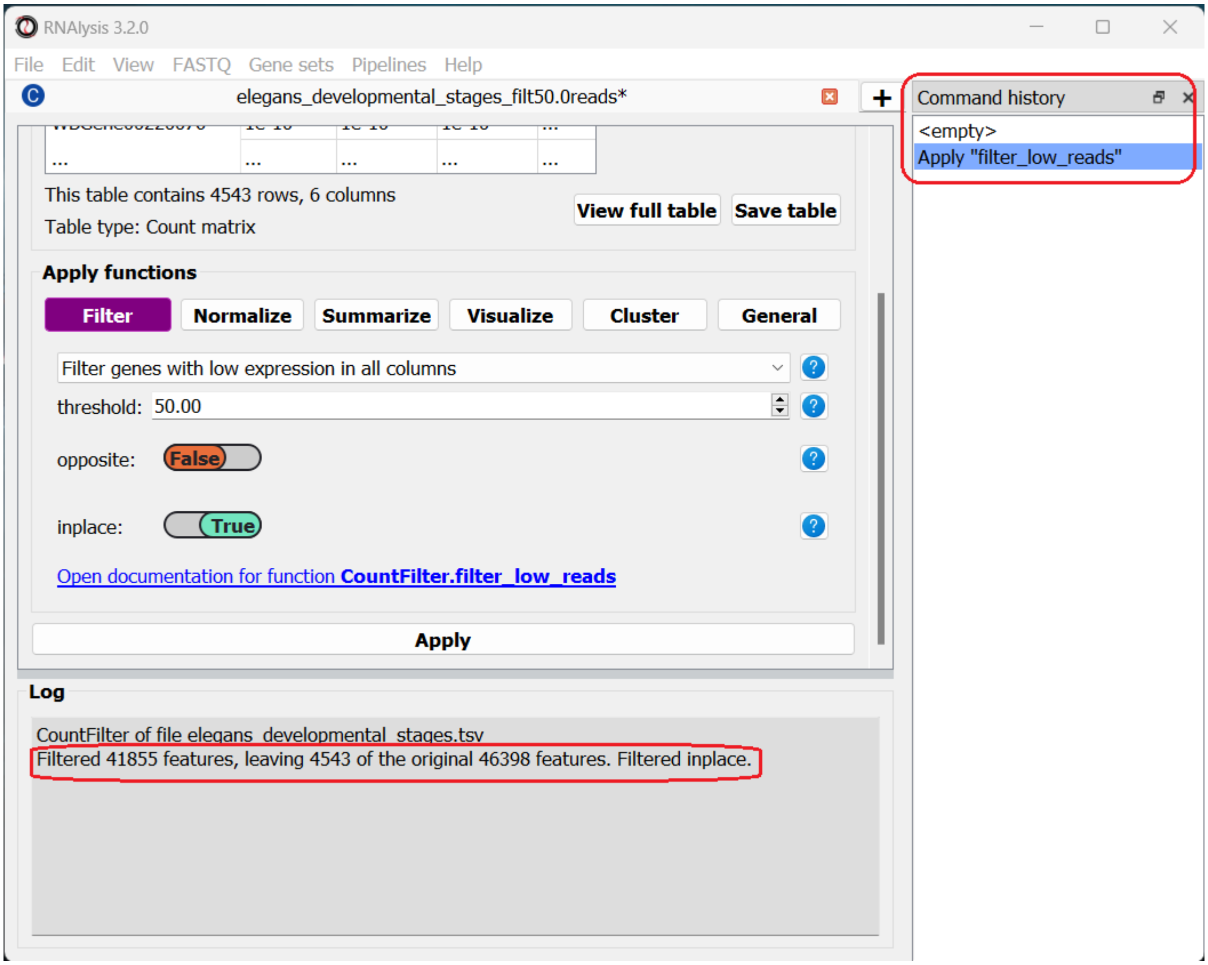


#### Examine variance in our data with Principal Component Analysis

The first analysis we will perform is Principal Component Analysis, or PCA. Principal component analysis is a dimensionality-reduction method. Meaning, it can reduce the dimensionality of large data sets (e.g. the expression of thousands of genes), by transforming the expression data of these genes into a smaller dataset that still contains most of the information from the original dataset.

Since our dataset contains thousands of genes, it can be difficult to see how the different conditions vary in the expression of those genes. To remedy that, PCA analysis can "rearrange" the axes of our data such that we can capture most of the variance of our dataset in very few dimensions (~2-10 dimensions).

You can read more about PCA here.

To run a PCA, click on the "Visualize" tab, and select "Principal Component Analysis" from the drop-down menu.

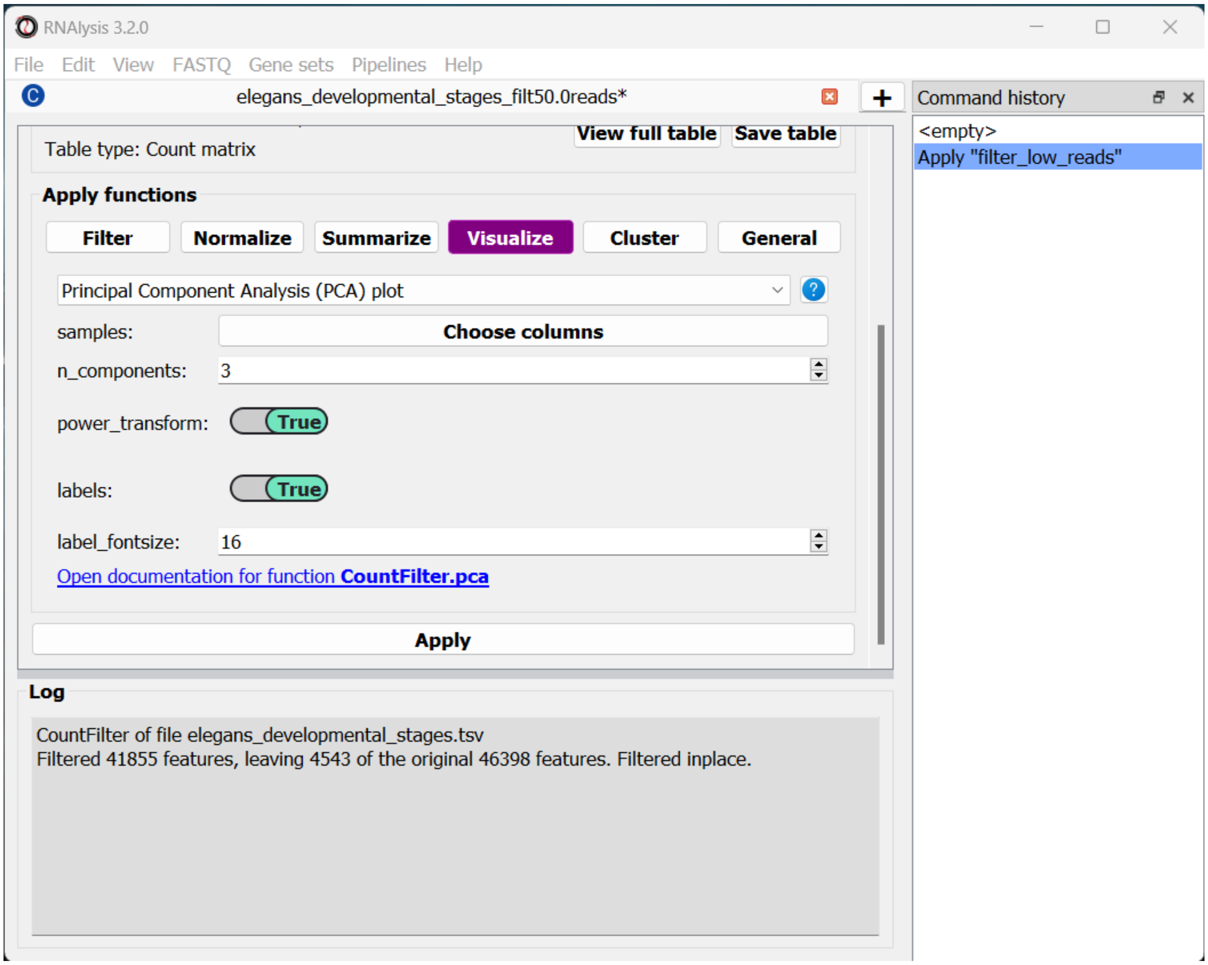


The 'samples' parameter allows you to choose which samples will be analyzed with PCA, and also lets you group these samples into sub-groups for the purpose of visualization (for example, group replicate data by experimental condition). That way, each sub-group will be drawn with a different color on the final graph. In our case we only have one column per condition, and we want to examine them all, so we don't need to change this parameter.

By default, *RNAlysis* will apply a power-transform (Box-Cox) to the data before standardazing it and running PCA. This is the case for many functions in *RNAlysis*, since applying power transform minimizes undesirable characteristics of counts data, such as skeweness, mean-variance dependence, and extreme values. However, this feature can always be disabled with the *power_transform* parameter.

Whether we apply a power transform to our data or not, *RNAlysis* will then standardize our data to neutralize any differences in absolute gene expression level, and then applies the Principal Component Analysis

Let's apply the analysis and look at the output:

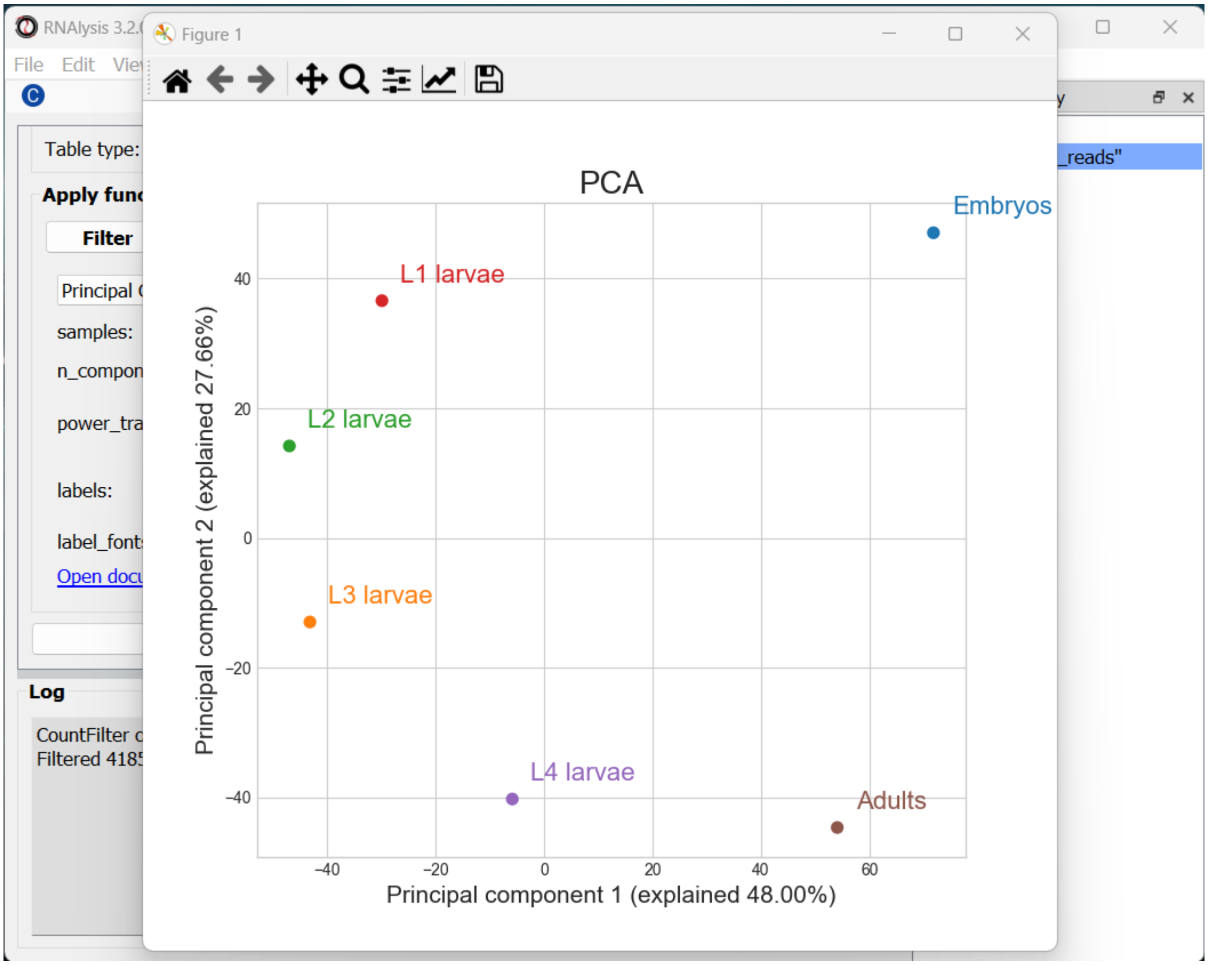


*RNAlysis* generated three graphs, depicting all of the pair-wise combinations between Principal Components #1-3. We can visualize less or more principal components by changing the value of the *n_components* parameter.

Usually the most relevant graph is the one depicting the first and second PCs, since they explain the largest amount of variance in the data. However, when the % of variance explained by preceeding PCs is similar (perhaps PCs 2 and 3 explain about the same % of variance), some of the PCs may be more informative than others, so it's worth keeping an eye out for those.

In our case, we can see that PCs 1-2 together explain ~75% of the variance in our dataset. Interestingly, the PCA shows a semi-circular pattern, ordered by the developmental stage of the worms. My hypothesis would be that PC1 arranges the samples by their relative "germline" content - embryos are mostly gonads, adult nematodes contain a rather large quantity of germ cells, L4 larvae is the developmental stage where germline proliferation begins, and during the L1-L3 stages the relative germline content of the worms is relatively minimal. PC2 appears to arrange the samples by their developmental stage, with embryos appearing at the top of the graph and adults at the bottom.

#### Examine similarity between developmental stages

Let's now examine the distribution of gene expression across developmental stages with a Pair-plot. Pair-plots displays the pairwise relationships between samples, or experimental conditions, in our dataset, and also display a histogram of gene expression values within each sample/condition.

To generate a Pair-plot, select "Pair-plot" from the function drop-down menu.

The Pair-plot function allows you to group samples by experimental condition in order to average them together.

We also have the option, like with PCA, to choose whether or not to power transform our data.

Let's click "Apply" and check out the result:

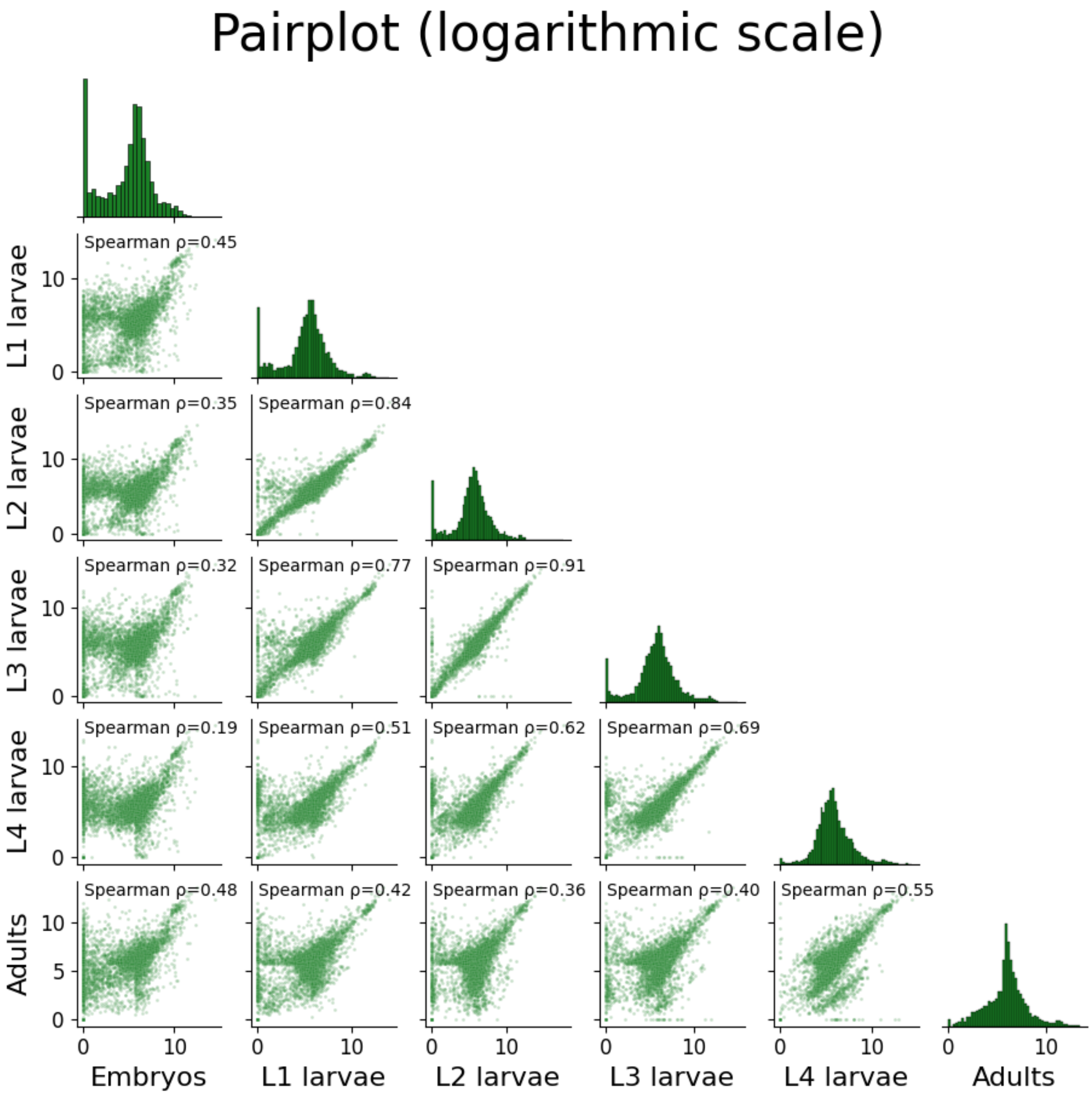

We can spot some interesting trends from this plot: the embryo condition seems to be the most dissimilar from the rest of the conditions; sequential developmental stages seem to be fairly correlated with one another; and the earlier developmental stages appear to have a larger fraction of unexpressed genes compared to the later developmental stages.

#### Compare the expression of specific genes over the developmental stages

We already have a hypothesis about the expression of some genes over the developmental stages, which we would like to test. For example, the gene *oma-1* (gene ID **WBGene00003864**) should be expressed almost exclusively in the two-cell stage of embryonic development, and we expect the gene *skn-1* (gene ID **WBGene00004804**) to show a fairly consistent expression level across development. Let's go to the "Visualize" tab to plot the expression of these genes over the developmental stages.

The *features* parameter will let us choose which genomic features we want to plot expression for. Since we can decide to add as many genes as we would like to this graph, *RNAlysis* will give us the option to choose how many genes to input. To start, click on the "Set input" button next to the *features* parameter:

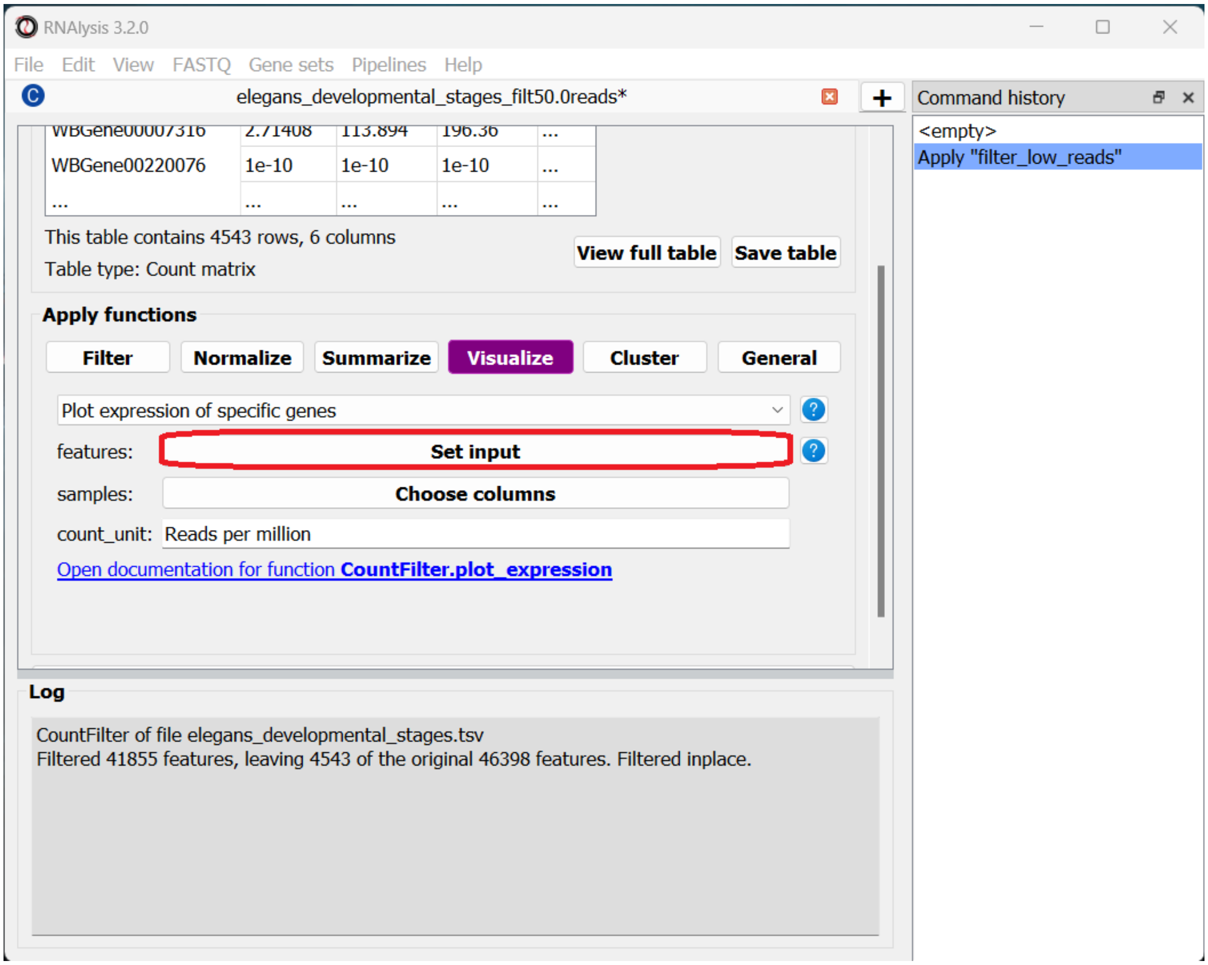

In the pop-up window that opens, you can use the “Add field” and “Remove field” buttons to choose how many genes to plot the expression of. Then, in each field, enter the gene ID of a gene you want to visualize. When you’re done, click on the “Done” button.

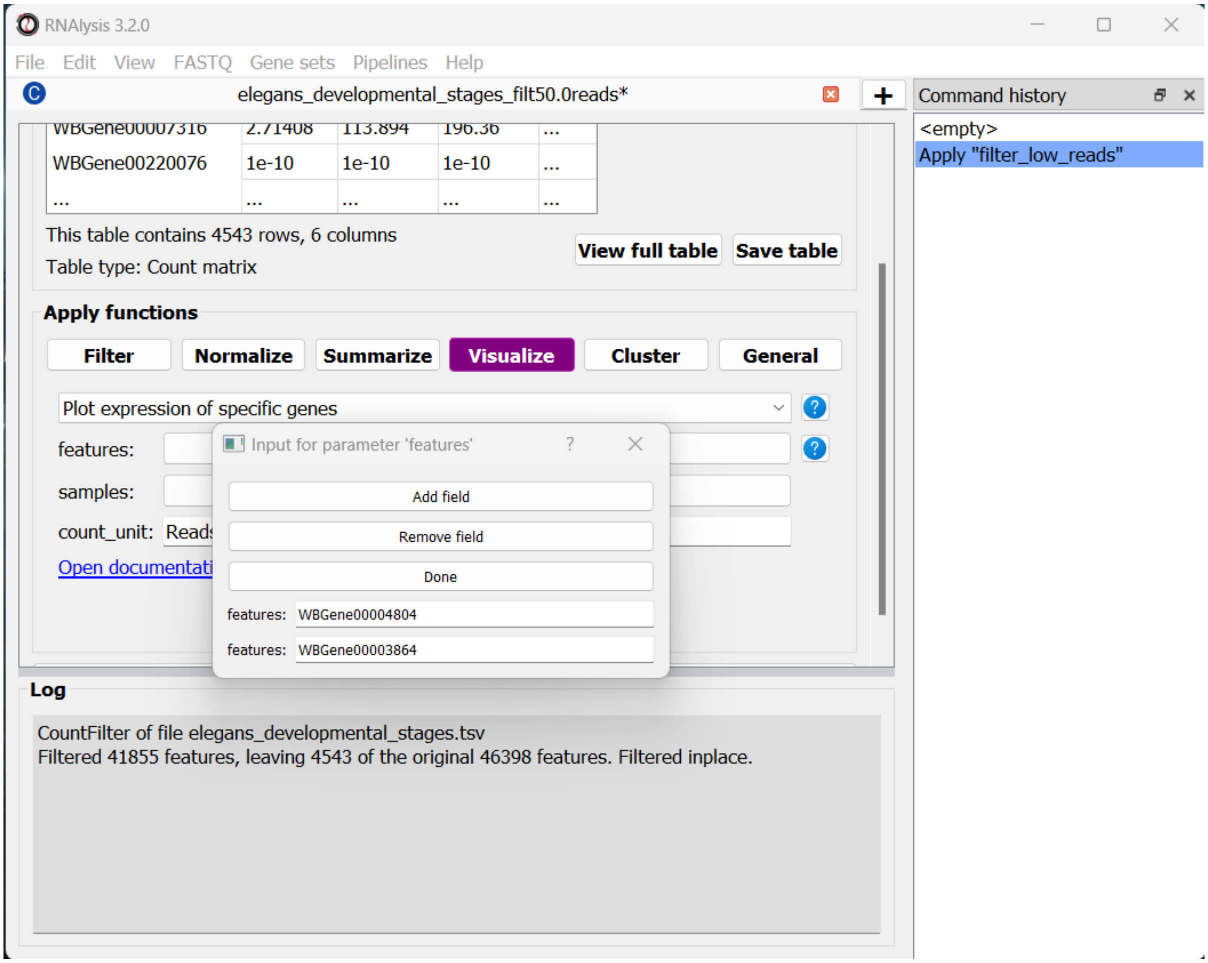


Let’s also set the count unit to FPKM, and click “Apply” to create the plot:

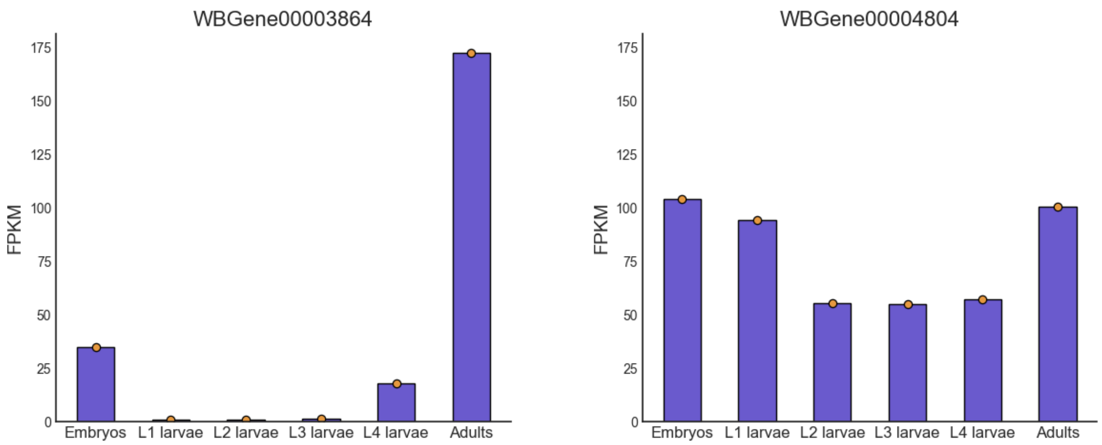


Interestingly, we can see that the expression of *oma-1* is actually highest in the adult worms. This is possibly because the adult worm contains a large number of unlaid embryos, some of which are the two-cell stage.

Our data only has one sample per condition, so our bar plots do not convey any information about the variance in gene expression within each condition. However, if we had multiple sample per condition, we could have grouped the samples into subgroup (like with the PCA and Pair-plot functions), and then each bar plot will also display a scatter of the expression values in each sub-group, and the standard error of the expression value.

#### Clustering analysis

We are interested in how different groups of genes change in expression level over the life cycle of the worm. We can use clustering analysis to group the genes in this dataset by their expression pattern over the developmental stages. There is an abundance of approaches when it comes to clustering analysis of gene expression. To illustrate this point, we will cluster our data using three different types of clustering algorithms, arranged from the simplest to the most complex. These algorithms are only a few representatives of the many clustering methods available in *RNAlysis*.

##### The simple approach - distance-based clustering with K-Medoids

K-Medoids clustering is a distance-based clustering method, where the algorithm attempts to divide all of our genes into K clusters (K is the number of clusters we are looking for), with each cluster being centered around one gene (Medoid). K-Medoids clustering takes in two major parameters - the number of clusters we expect (K), and the distance metric by which we want to measure the similarity of expression between genes.

Specifying the exact number of clusters we expect is a bit challenging for us, since we aren't actually sure how many biologically-meaningful clusters are there in our data. Moreover, this number could depend on how fine-grained we want our analysis to be - we could reasonably divide our genes into a small number of more generalized clusters (such as "genes expressed more in the start of development" vs "genes expressed more near the end of development"), or we could further divide our genes into smaller groups based on their exact temporal expression pattern.

Fortunately, some methods were developed to suggest a good number of clusters for our dataset (a "good number of clusters" meaning that the genes in each clusters are most similar to each other and most different than genes in other clusters). Two of these methods, known as the Silhouette Method and the Gap Statistic, are available in *RNAlysis*. We will use the Gap Statistic method to determine some good options for the number of clusters in our analysis.

To start, let's click on the Clustering tab and choose K-Medoids clustering from the drop-down menu. We can then set the value of the parameter *n_clusters* to 'gap', to indicate that we want to use the Gap Statistic to determine the number of clusters in this analysis:

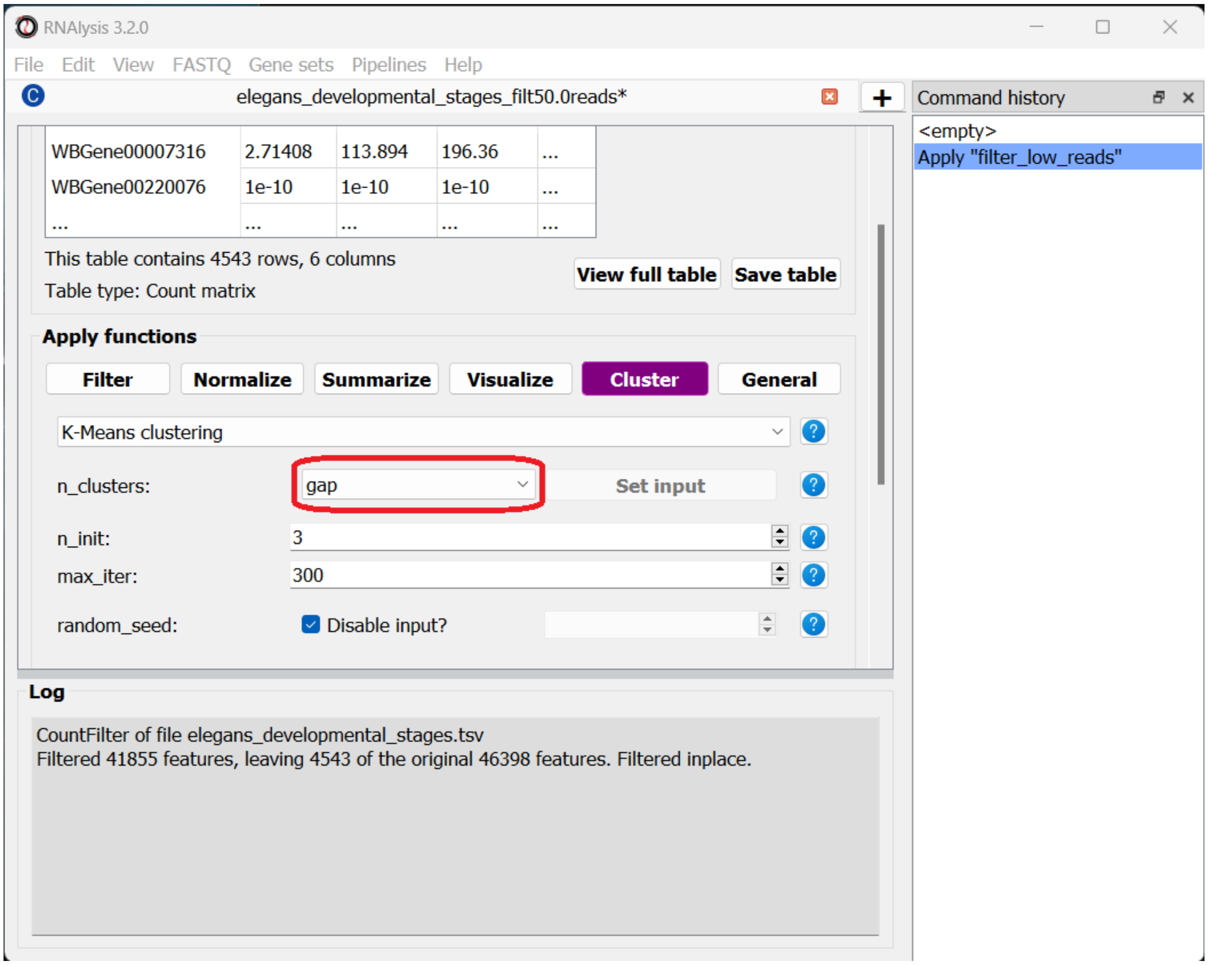


Next, we can set the distance metric. Different distance metrics can be more or less effective on specific types of data. *RNAlysis* offers a large array of distance metrics, about which you can read in the *RNAlysis* user guide. We will use a lesser-known distance metric called YR1, that was developed especially for time-series gene expression data and implemented in *RNAlysis*. You can read more about it in Son and Baek 2007:

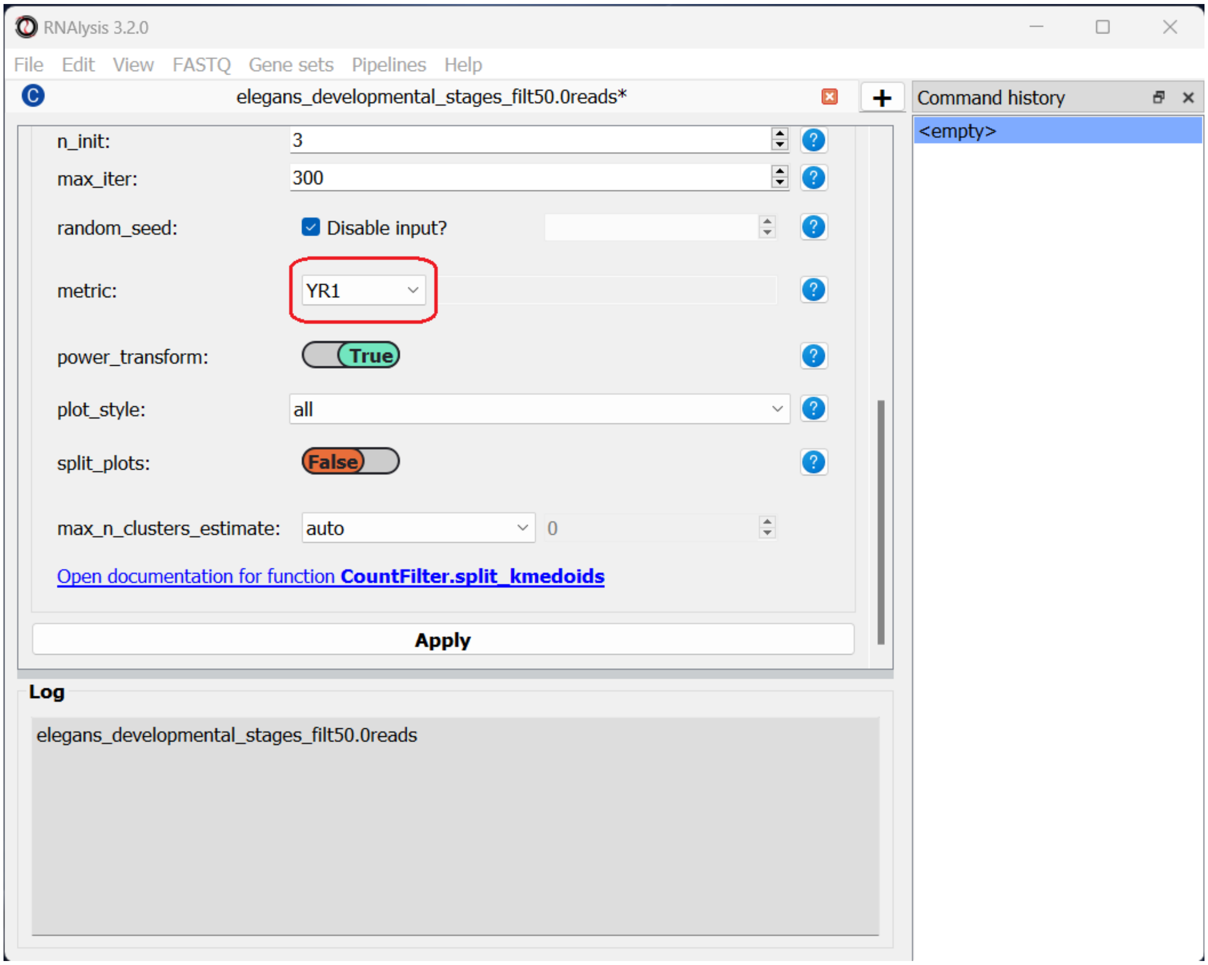


We can now scroll all the way down, click the “Apply” button, and wait for the analysis to finish:

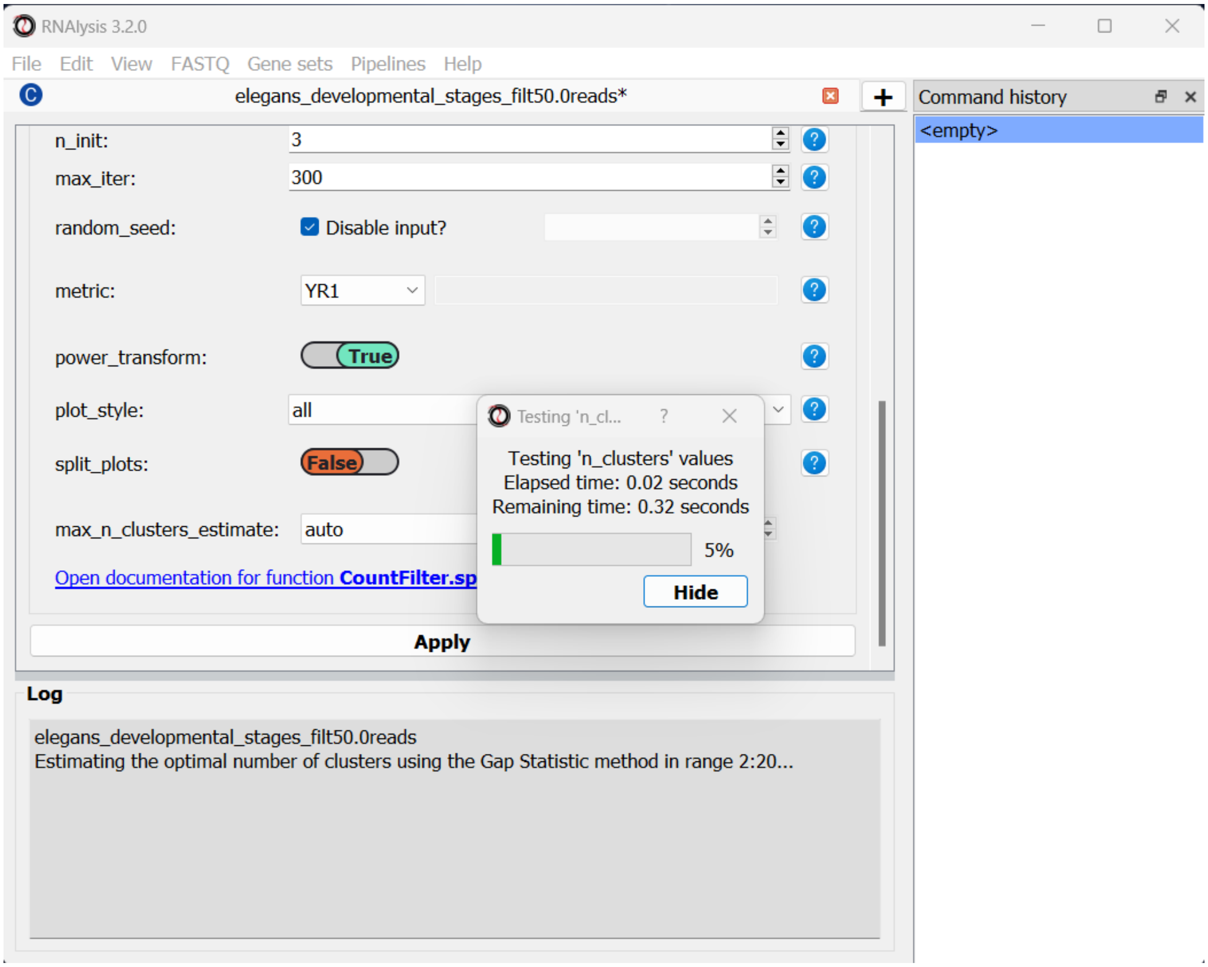


Since K-Medoids is not a deterministic algorithm (it has randomized starting conditions every time you use it), the results you get will probably not be identical to those appearing in this tutorial, but they should be similar to what we observe here. If you want to make sure your clustering results are 100% reproducible, you can set the *random_seed* parameter when setting up your clustering setup to a specific number. Re-running the algorithm with an identical random seed wiil ensure that you get the exact same results every time.

One the clustering anslysis is finished, a few figures will open up. Let's examine them one by one. The first figure will show us the results of the Gap Statistic algorithm. The graph on the left will show us, for each value of *n_clusters* tested, the natural logarithm (ln) of within-cluster dispersion. We expect this value to go down the more clusters there are (the more clusters there are, the fewer genes will be in each cluster - therefore the genes within each cluster will be more similar to each other), and therefore we show both the actual dispersion values for the clustering solutions we calculated (the blue line), and also dispersion values for clustering solutions on random data, drawn from the same distribution (the orange line). On the graph to the right we can see the natural logarithm of the ratio between the observed and expected dispersion - this is the Gap Statistic. We are looking for local 'peaks' in the graph - values of *n_clusters* that have a larger Gap Statistic than their neighbors. *RNAlysis* automatically picks the lowest value of *n_clusters* that fits this criterion, but also suggests other good values based on the results.

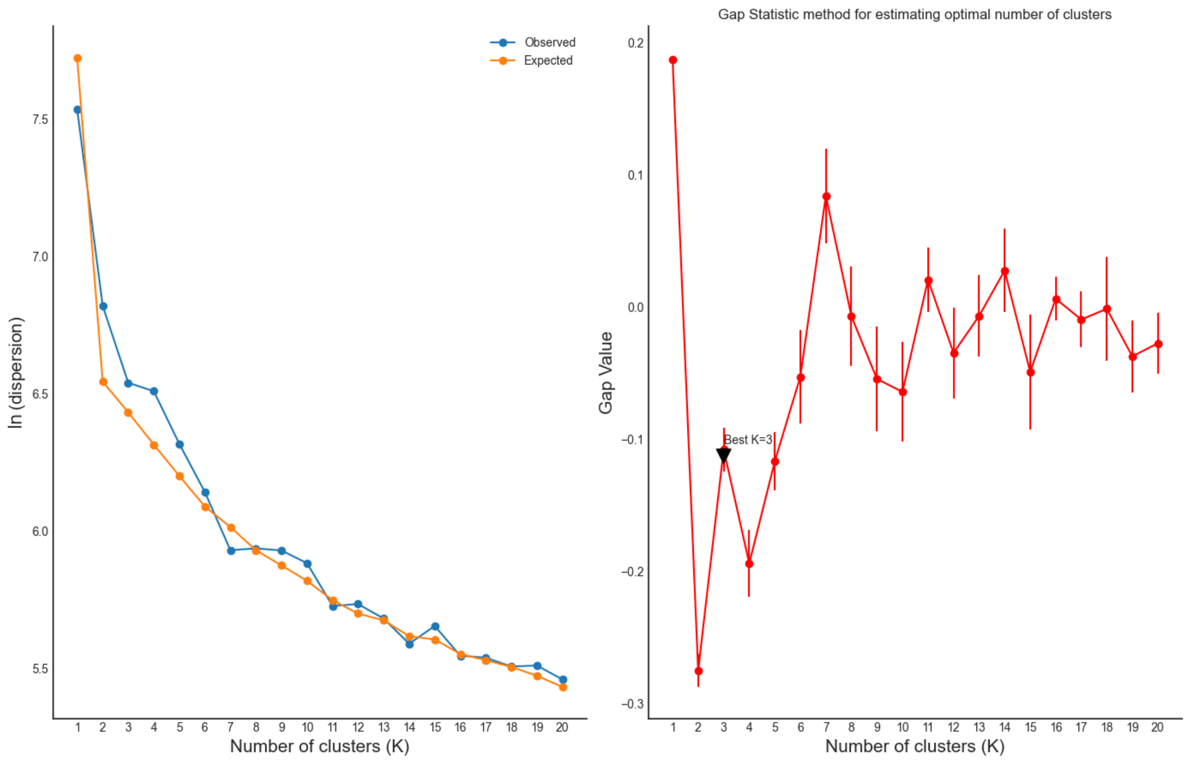


In our case, *RNAlysis* recommended 3 clusters as the optimal number of clusters. This value might not be granular enough for the kind of analysis we want to run. Therefore, we will re-run the K-Medoids algorithm with the same parameters, but set the value of *n_clusters* to one of the next good values discovered in the Gap Statistic analysis - 11 clusters.

Next, we can look at the rest of the output of the K-Medoids clustering algorithm for `n_clusters`=11. The first graph will show us the distributions of gene expression in each discovered cluster. Note that the expression values are regularized and power-transformed, since we are interested in grouping the different genes by their relative pattern of expression, and not by their absolute expression levels (highly/lowly-expressed genes). The clusters are sorted by their size, from the biggest to the smallest cluster. This type of graph can help us see the general expression pattern that characterizes each cluster. Moreover, it can help point out how internally similar our clusters are - indicating the quality of our clustering result.

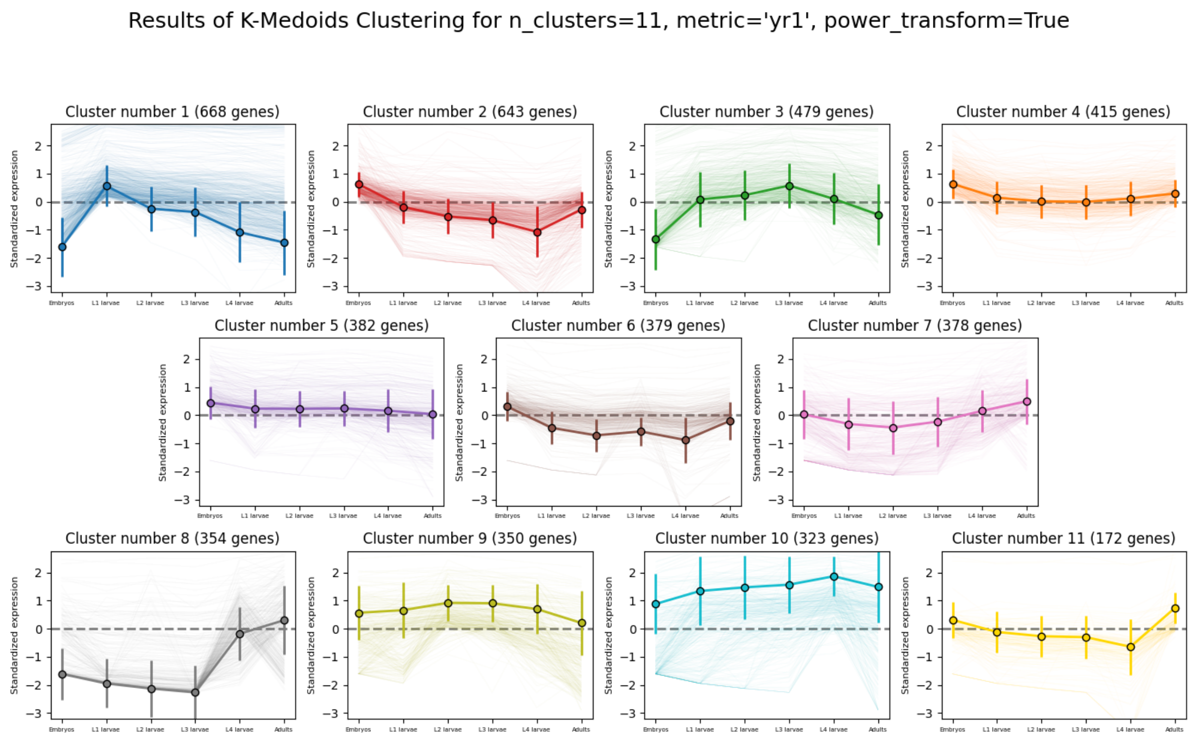


In this case, we can see that while some clusters seem very internally consistent, quite a few clusters seem to contain a significant number of 'outlier' genes.

*RNAlysis* also generates a Principal Component Analysis projection of our gene expression data, marking the genes in each cluster with a different color. This is another useful way to look at our clustering results - we would hope to see that the first two principal components explain a large degree of the variance in gene expression, and the genes in the same clusters will be grouped together in the graph.

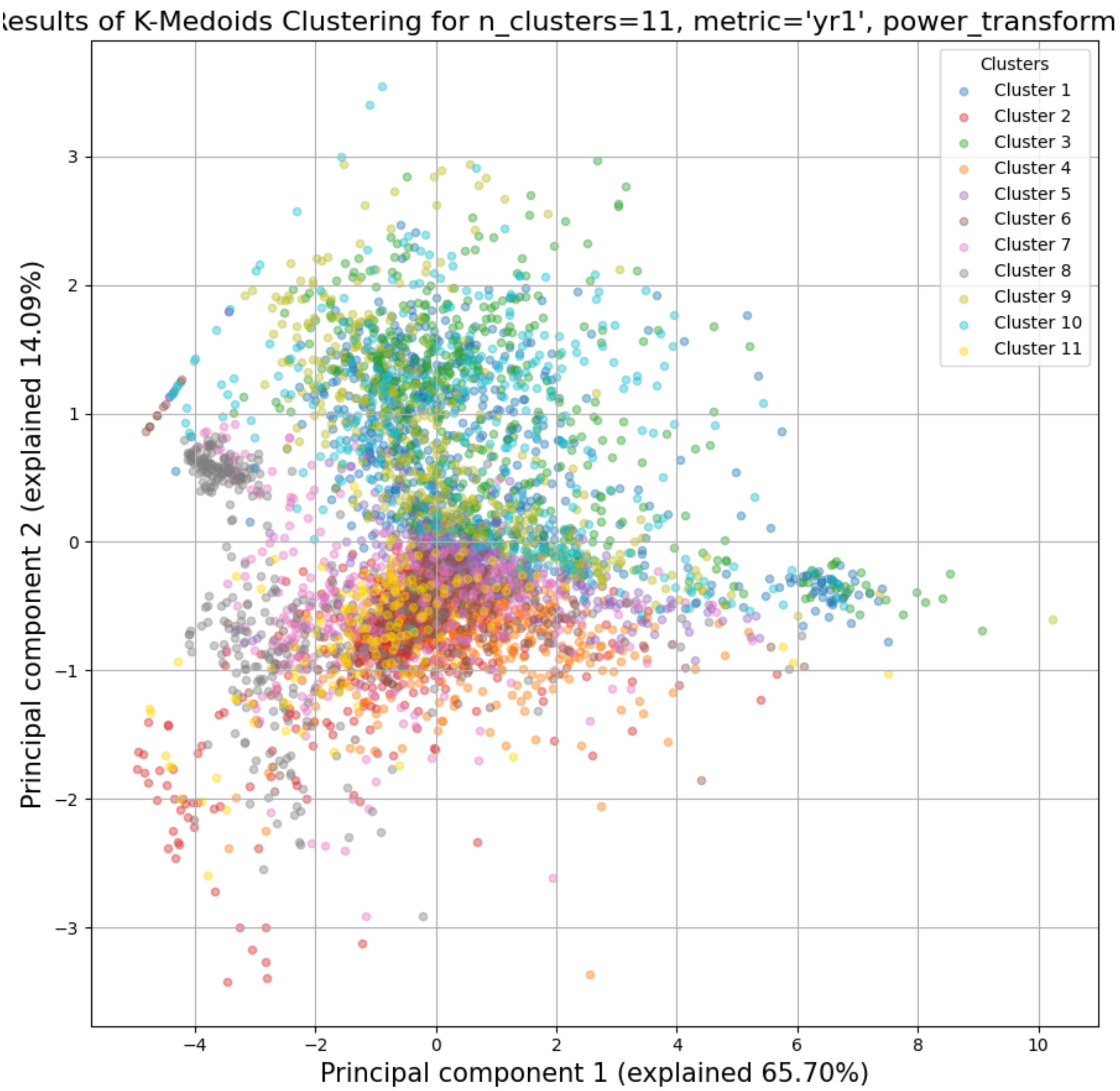


Finally, the following window will open, prompting us to choose which output clusters we want to keep, and giving us the option to give these clusters a new name:

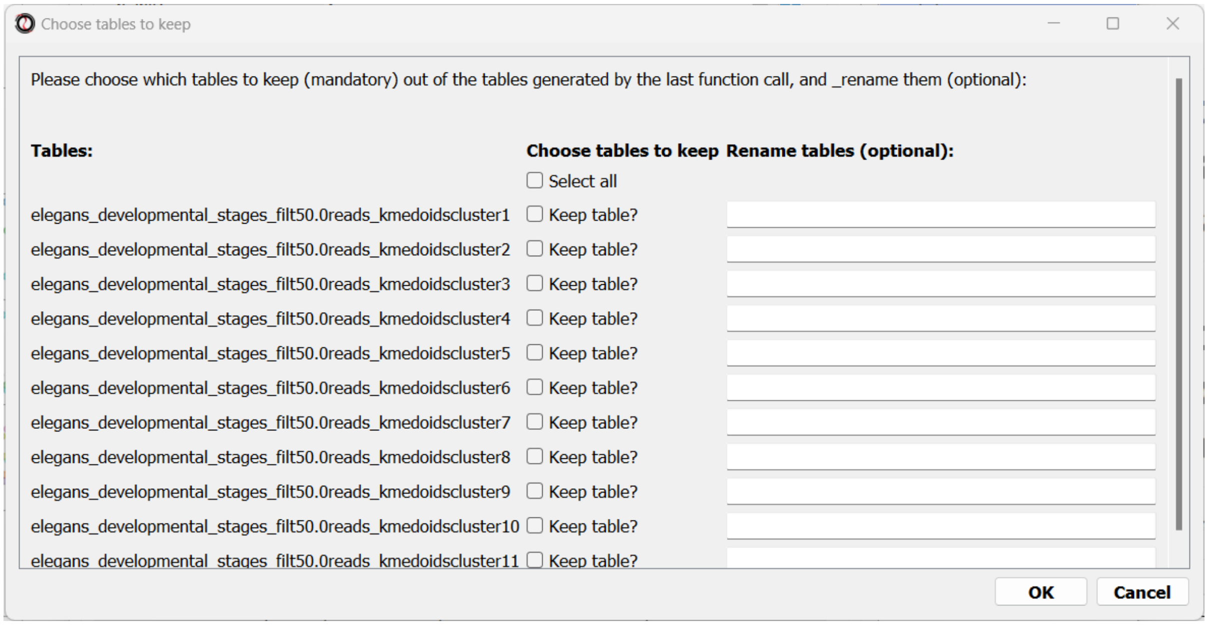


For every cluster we choose to keep, *RNAlysis* will create a new copy of the original count matrix, which includes only the genes that belong to this specific cluster. For now, we will choose to keep none of the clusters, so that we can try out other clustering approaches. Therefore, we click either OK or Cancel without selecting any clusters.

#### Fine-tuning our approach - density-based clustering with HDBSCAN

The next clustering approach we will use, HDBSCAN, belongs to a different category of clustering algorithms - density-based clustering. HDBSCAN stands for Hierarchical Density-Based Spatial Clustering of Applications with Noise (see the publication for more details). HDBSCAN offers multiple advantages over more traditional clustering methods:

1. HSBSCAN makes relatively few assumptions about the data - it assumes that the data contains noise, as well as some real clusters which we hope to discover.
2. Unlike most other clustering methods, HDBSCAN does not “force” every gene to belong to a cluster. Instead, it can classify genes as outliers, excluding them from the final clustering solution.
3. When using HDBSCAN, you don’t have to guess the number of clusters in the data. The main tuning parameter in HDBSCAN is *minimum cluster size* (*min_cluster_size*), which determines the smallest “interesting” cluster size we expect to find in the data.

To run HDBSCAN, we need to pick a value for *min_cluster_size*. Lower values of *min_cluster_size* will return a larger number of small clusters, revealing more fine-grained patterns in our gene expression data. Higher values of *min_cluster_size* will return a smaller number of large clusters, revealing the most obvious and significant patterns in the data. For our example, let’s pick a value of 75:

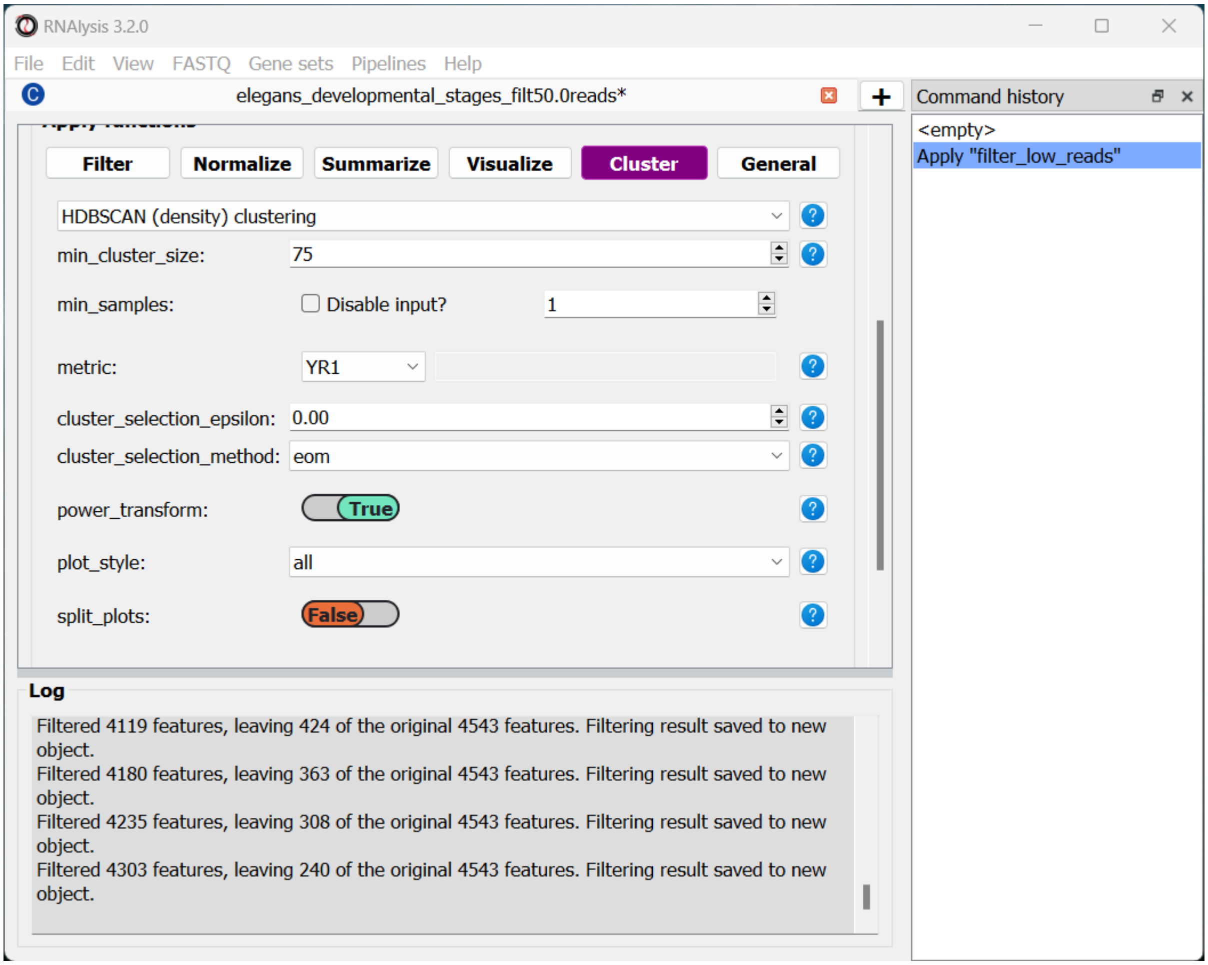


We will, once again, use YR1 as the distance metric.

If we look at the clustering results, we can see that HDBSCAN ended up generating a much larger number of clusters than the previous method, and they look fairly internally consistent.

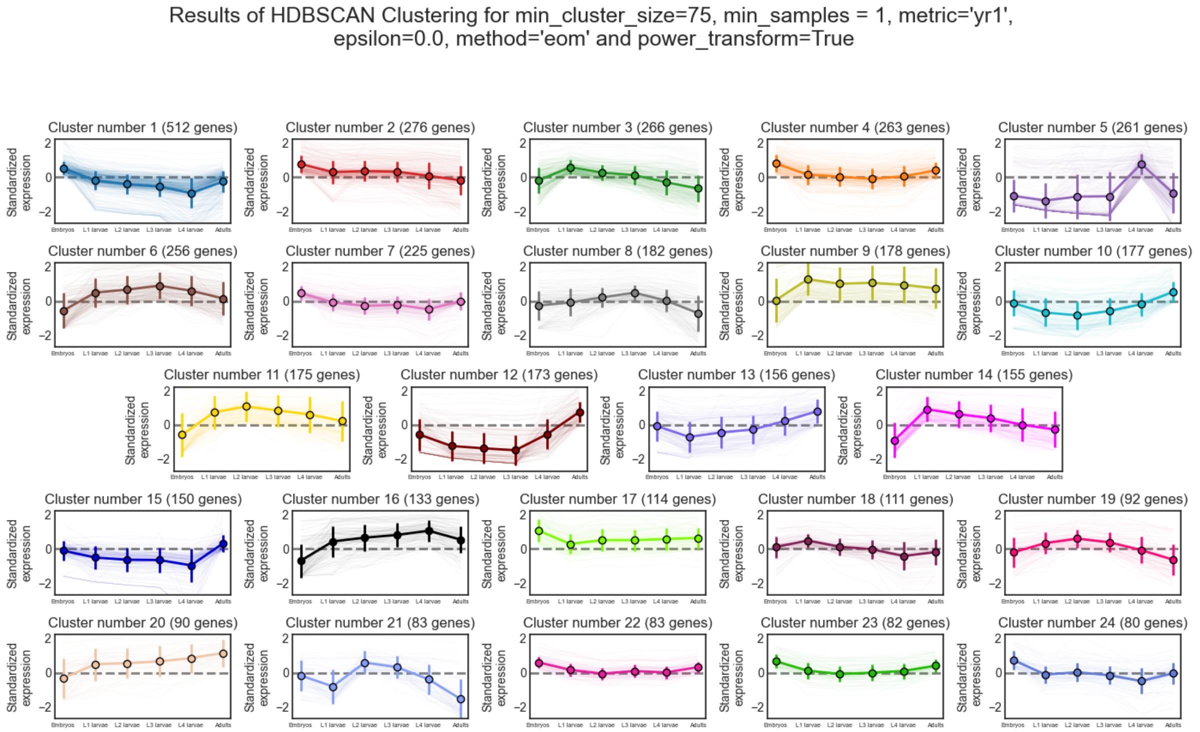
Results of HDBSCAN Clustering for min_cluster_size=75, min_samples = 1, metric='yr1',
epsilon=0.0, method='eom' and power_transform=True
Cluster number 1 (512 genes)
Cluster number 2 (276 genes)
Cluster number 3 (266 genes)
Cluster number 4 (263 genes)
Cluster number 5 (261 genes)
Cluster number 6 (256 genes)
Cluster number 7 (225 genes)
Cluster number 8 (182 genes)
Cluster number 9 (178 genes)
Cluster number 10 (177 genes)
Cluster number 11 (175 genes)
Cluster number 12 (173 genes)
Cluster number 13 (156 genes)
Cluster number 14 (155 genes)
Cluster number 15 (150 genes)
Cluster number 16 (133 genes)
Cluster number 17 (114 genes)
Cluster number 18 (111 genes)
Cluster number 19 (92 genes)
Cluster number 20 (90 genes)
Cluster number 21 (83 genes)
Cluster number 22 (83 genes)
Cluster number 23 (82 genes)
Cluster number 24 (80 genes)
Standardized expression

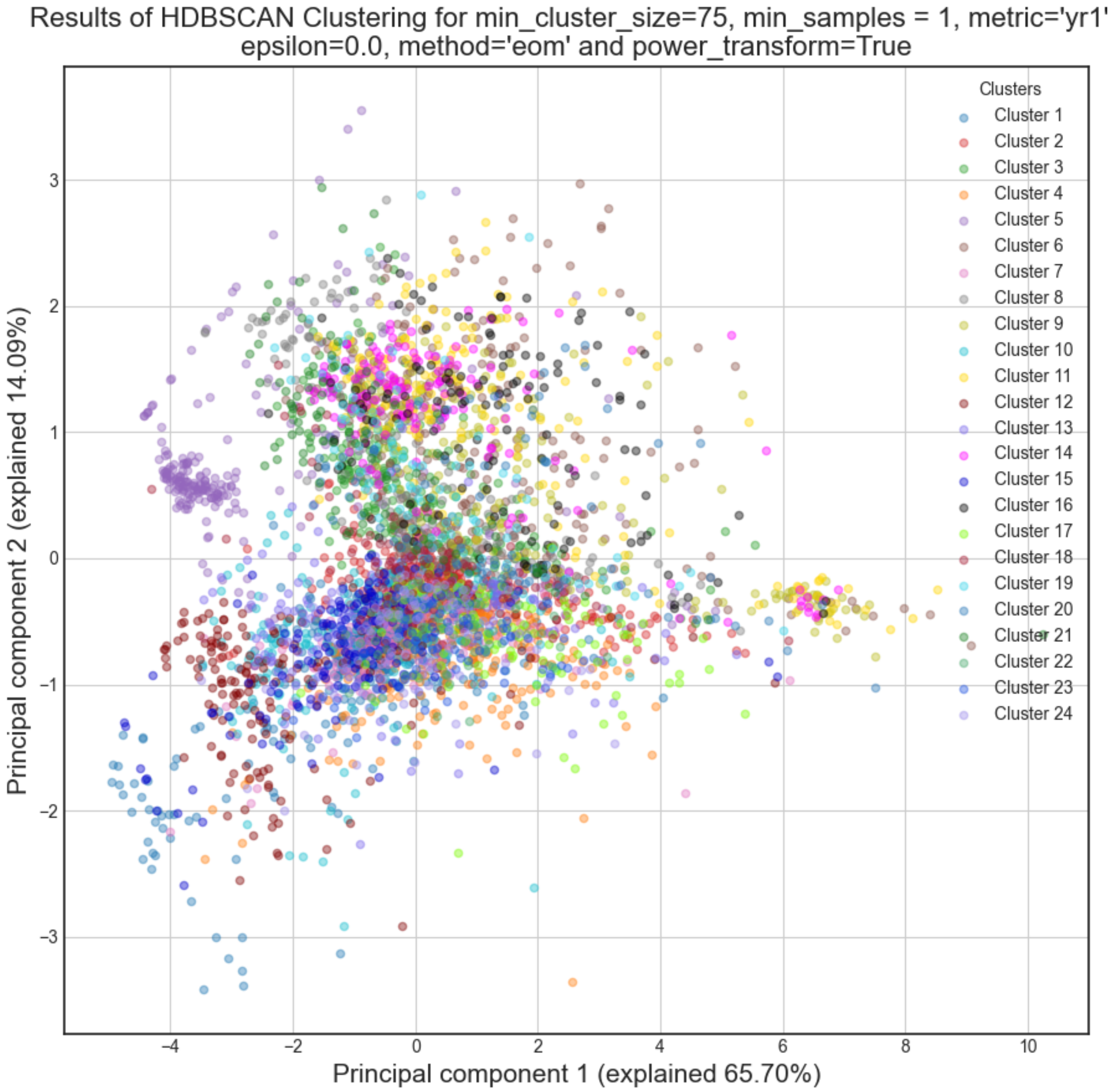
Results of HDBSCAN Clustering for min_cluster_size=75, min_samples = 1, metric='yr1'
epsilon=0.0, method='eom' and power_transform=True
Principal component 1 (explained 65.70%)
Principal component 2 (explained 14.09%)
Clusters
Cluster 1
Cluster 2
Cluster 3
Cluster 4
Cluster 5
Cluster 6
Cluster 7
Cluster 8
Cluster 9
Cluster 10
Cluster 11
Cluster 12
Cluster 13
Cluster 14
Cluster 15
Cluster 16
Cluster 17
Cluster 18
Cluster 19
Cluster 20
Cluster 21
Cluster 22
Cluster 23
Cluster 24

Let's now move on to the final clustering approach - ensemble-based clustering.

##### The complex approach - ensemble-based clustering with CLICOM

The last clustering approach we will use, CLICOM, is an emsemble-based clustering algorithm. CLICOM (see the publication ) incorporates the results of multiple clustering solutions, which can come from different clustering algorithms with differing clustering parameters, and uses these clustering solutions to create a combined "concensus" clustering solution. CLICOM offers multiple advantages over more traditional clustering methods:

1. The ensemble clustering approach allows you to combine the results of multiple clustering algorithms with multiple tuning parameters, potentially making up for the weaknesses of each individual clustering method, and only taking into account patterns that robustly appear in many clustering solutions.
2. When using CLICOM, you don't have to guess the final number of clusters in the data. The main tuning parameter in HDBSCAN is the *evidence threshold* (*evidence_threshold*).

*RNAlysis* offers a modified implementation of CLICOM. The modified version of the algorithm can, like the HDBSCAN algorithm, classify genes as outliers, excluding them from the final clustering solution. Moreover, ths modified version of the algorithm can cluster each batch of biological/technical replicates in your data separately, which can reduce the influence of batch effect on clustering results, and increases the accuracy and robustness of your clustering results.

This modified version of CLICOM supports a few tuning parameters, in addition to the clustering solutions themselves:

- *evidence_threshold*: how many clustering solutions (fraction between 0-1) have to agree about two genes being clustered together in order for them to appear together in the final solution? A lower evidence threshold leads to fewer, large clusters, with fewer features being classified as outliers.
- *cluster_unclustered_features*: if set to True, CLICOM will force every gene to belong to a discovered cluster (like in the original implementation of the algorithm). Otherwise, genes can be classified as noise and remain unclustered (the modified algorithm).
- *min_cluster_size*: determines the minimal size of a cluster you would consider meaningful. Clusters smaller than this would be classified as noise and filtered out of the final result, or merged into other clusters (depending on the value of *cluster_unclustered_features*).
- *replicates_grouping*: allows you to group samples into technical/biological batches. The algorithm will then cluster each batch of samples separately, and use the CLICOM algorithm to find an ensemble clustering result from all of the separate clustering results.

To start the analysis, let's choose the CLICOM algorithm from the Clustering drop-down menu. A new window will open:

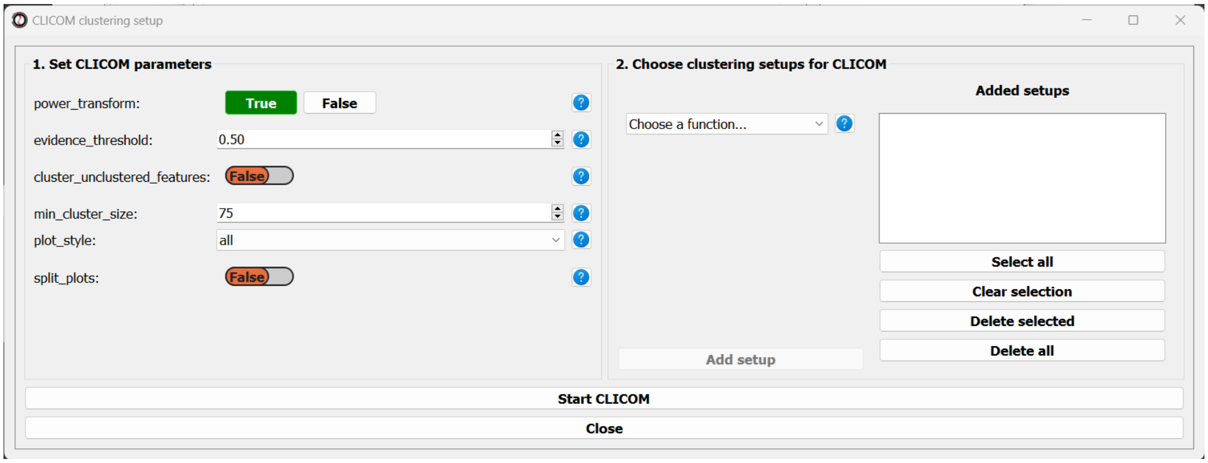


On the left half of the window we can set the tuning parameters of the CLICOM algorithm. For our example, let's set the evidence threshold to 0.5, and the minimum cluster size to 75. Since our data doesn't have replicates, we don't need to tune the *replicates_grouping* parameter. If in the future you try to cluster data with biological replicates, keep this parameter in mind!

On the right half of the window we can add new clustering setups to our run of CLICOM. These clustering setups can be any of the clustering algorithms available in *RNAlysis*, and we can add as many as we want - including multiple clustering setups using the same algorithm. To add a setup, pick an algorithm from the drop-down menu, set it's parameters, and click the "Add Setup" button:

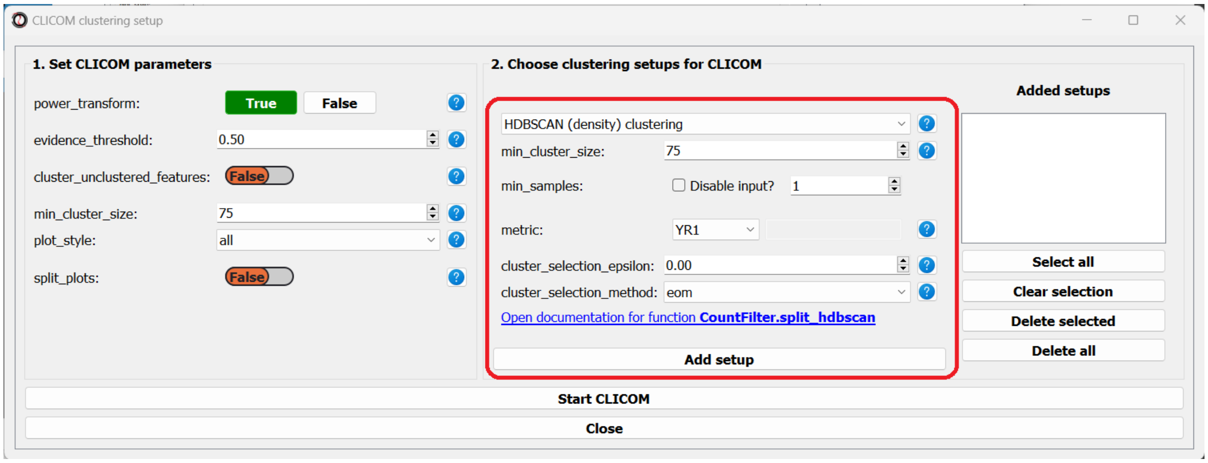


The setups you added will appear under "added setups" on the right, and you can delete a setup from that list if you want:

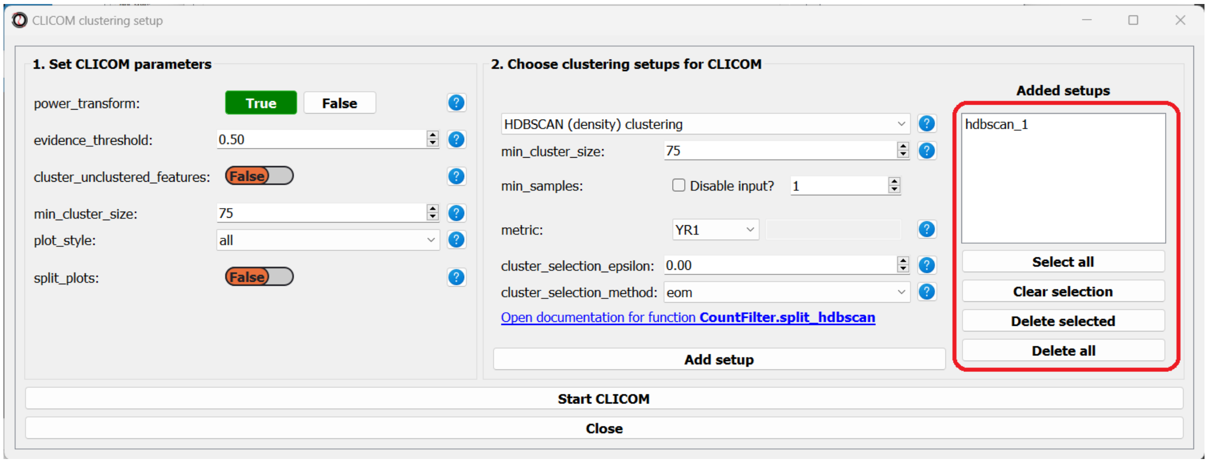


Let's add the two clustering setups we used earlier, plus a few more:

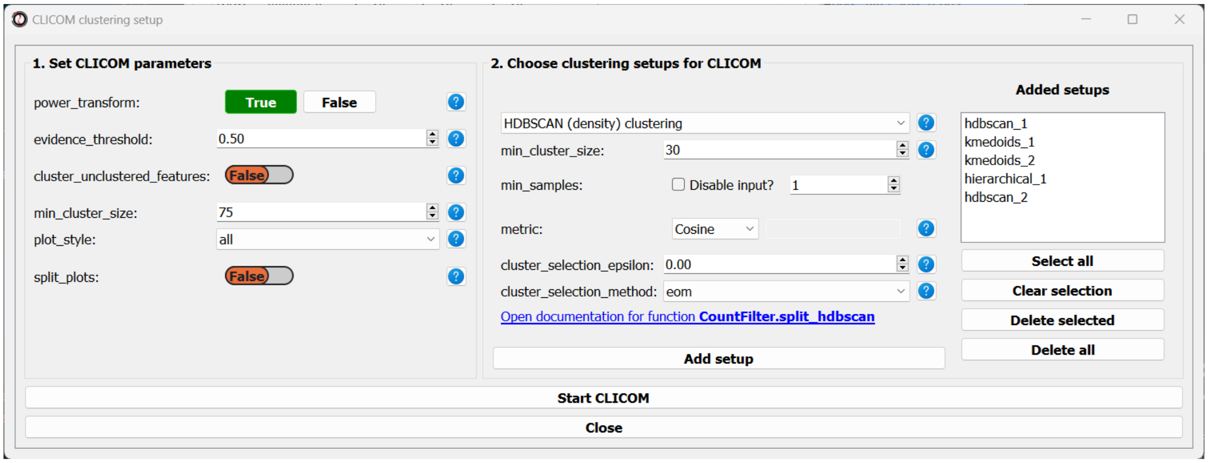


I chose to add, in addition to the two clustering setups from earlier, the following three clustering setups: * K-Medoids clustering with 7 clusters and the Spearman distance metric * Hierarchical clustering 8 clusters, Euclidean distance metric and Ward linkage * HDBSCAN clustering with the Jackknife distance metric and minimal cluster size of 150

Once we are happy with the clustering solutions and tuning parameters, we can click on the "Start CLICOM" button, and see progress reports in the output box on the main window of *RNAlysis*.

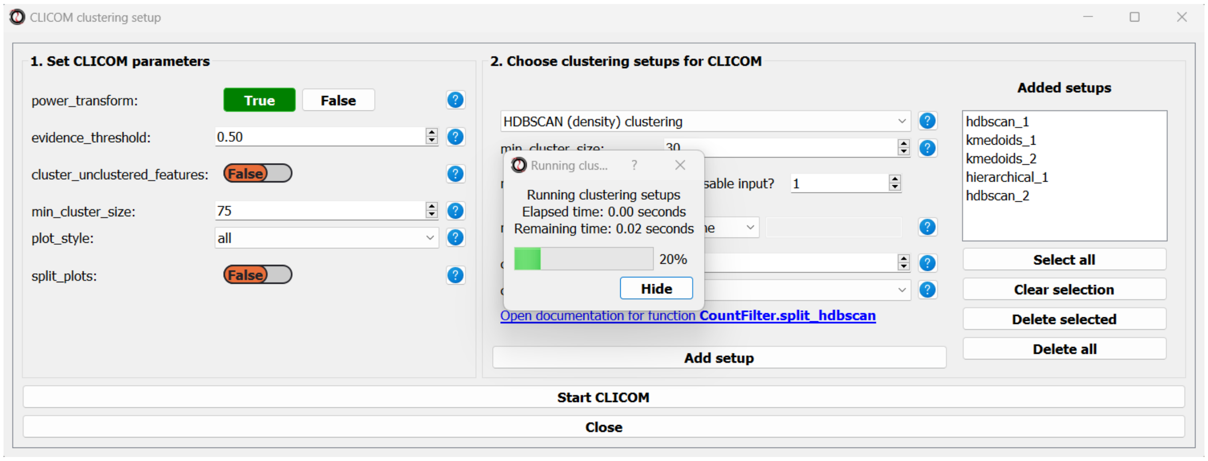

Let’s look at the final result:

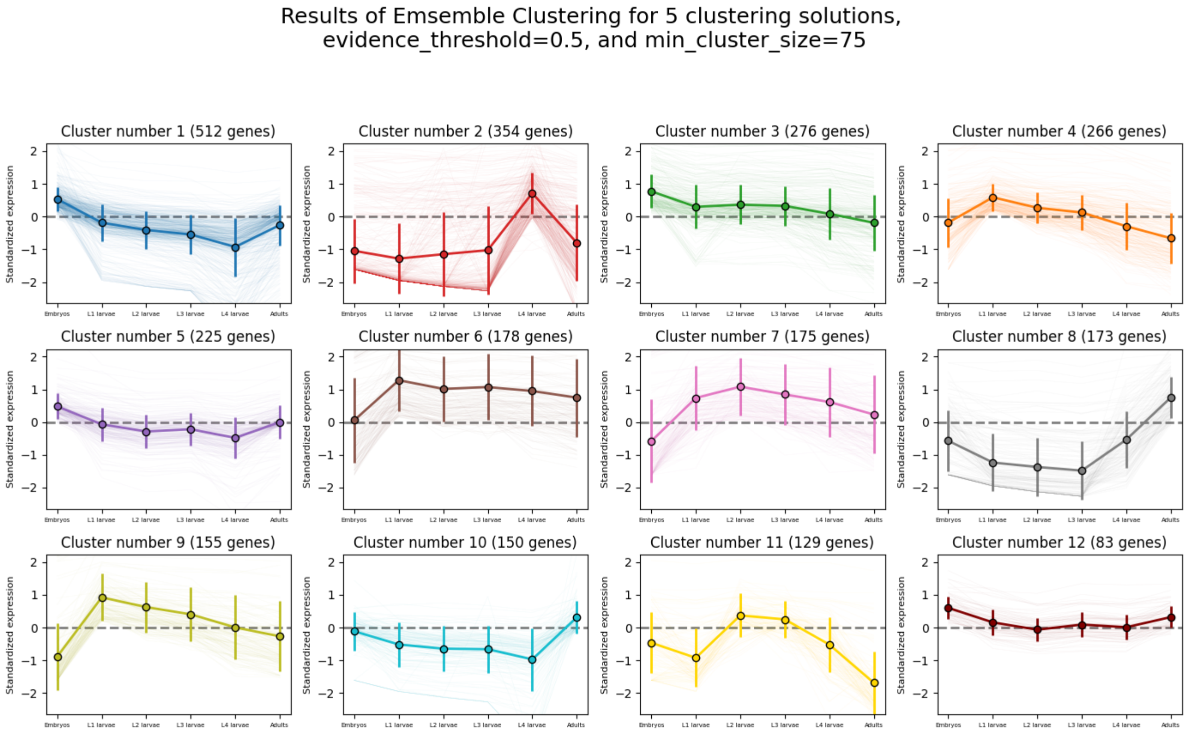

Results of Emsemble Clustering for 5 clustering solutions,
evidence_threshold=0.5, and min_cluster_size=75
Cluster number 1 (512 genes)
Cluster number 2 (354 genes)
Cluster number 3 (276 genes)
Cluster number 4 (266 genes)
Cluster number 5 (225 genes)
Cluster number 6 (178 genes)
Cluster number 7 (175 genes)
Cluster number 8 (173 genes)
Cluster number 9 (155 genes)
Cluster number 10 (150 genes)
Cluster number 11 (129 genes)
Cluster number 12 (83 genes)
Standardized expression
Embryos
L1 larvae
L2 larvae
L3 larvae
L4 larvae
Adults


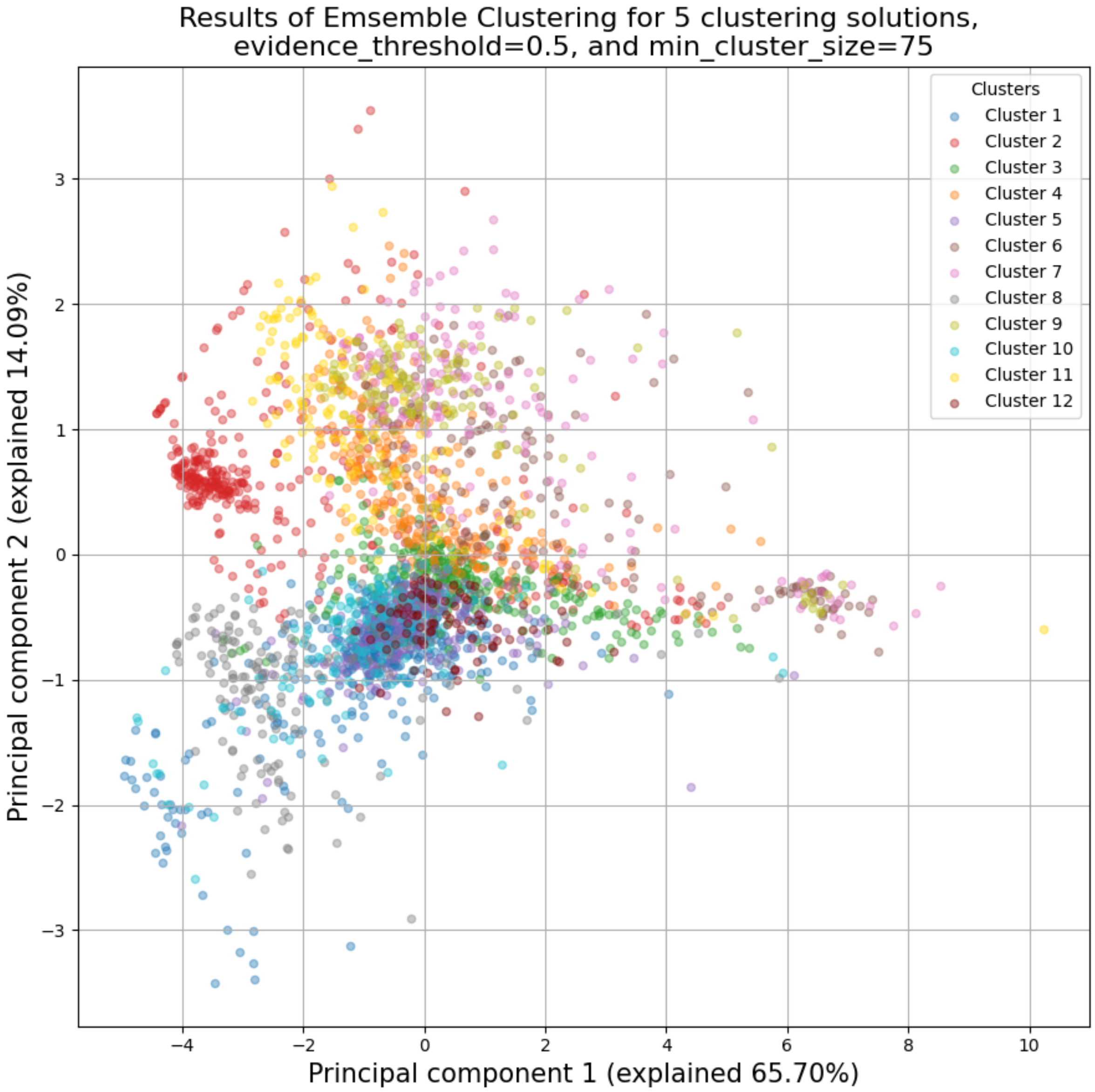

Results of Emsemble Clustering for 5 clustering solutions,
evidence_threshold=0.5, and min_cluster_size=75
Clusters
Cluster 1
Cluster 2
Cluster 3
Cluster 4
Cluster 5
Cluster 6
Cluster 7
Cluster 8
Cluster 9
Cluster 10
Cluster 11
Cluster 12
Principal component 2 (explained 14.09%)
Principal component 1 (explained 65.70%)

Once again, since two of the clustering setups we chose are non-deterministic, the clusters you end up with might end up looking a bit differently from these. Let's choose a few good-looking clusters to keep, and give them a name that indicates their expression pattern:

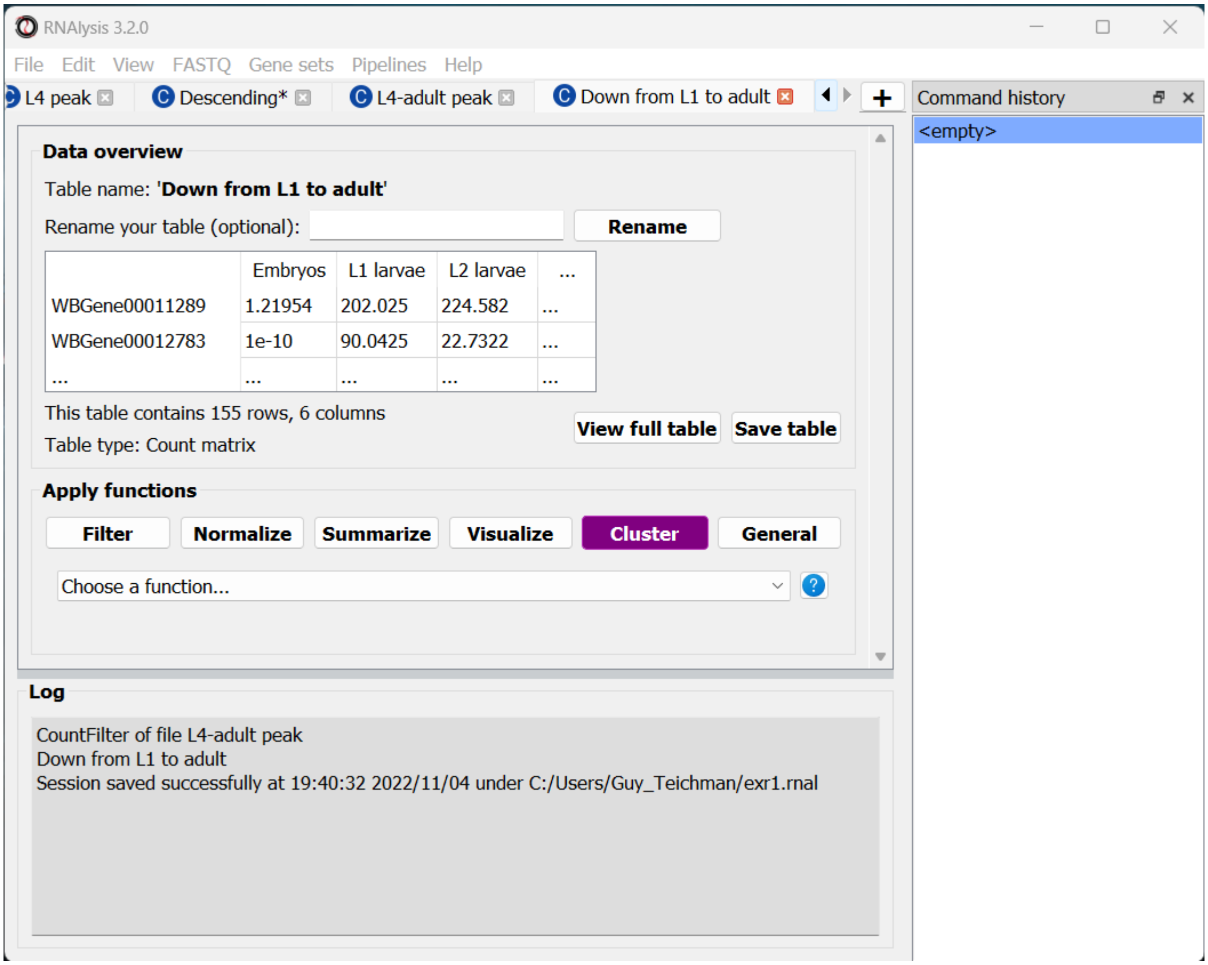


For this tutorial, I chose to keep clusters #1 ("down from embryo to L4"), #2 ("L4 peak"), and #9 ("Down from L1 to adult").

#### Enrichment analysis

Now that we have extracted a few clusters of interest, we can try to characterize their biological significance using enrichment analysis. We will look at the most commonly-used enrichment analysis - Gene Ontology enrichment.

##### Running enrichment analysis

To open the Enrichent Analysis window, open the 'Gene sets' menu and click "Enrichment Analysis":

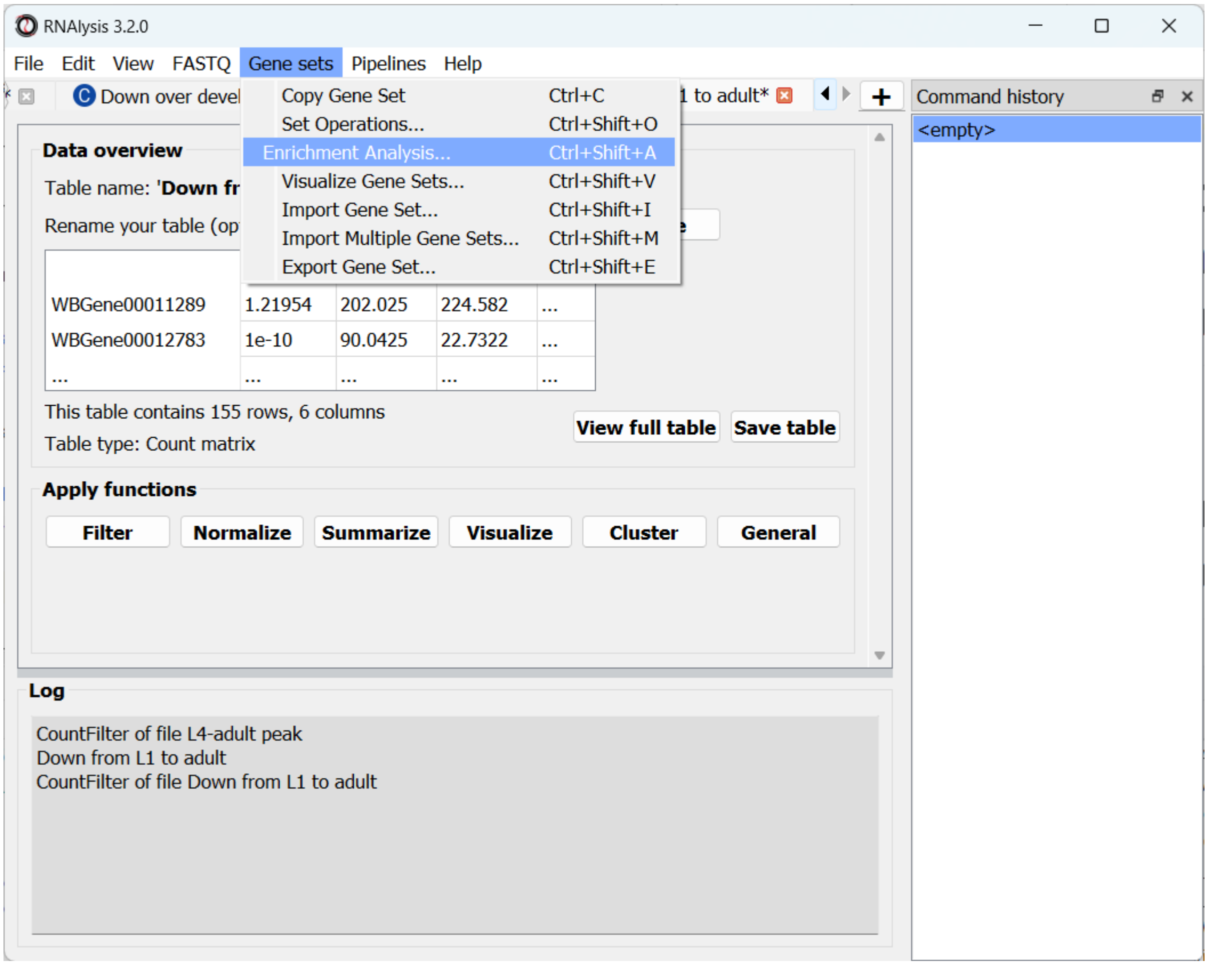


For basic enrichment analysis, we first need to choose our *enrichment set* (the gene set we are interested in - for example, "Down from L1 to adult", cluster #9 we found earlier), and our background set (the reference genes we will be comparing our enrichment results to - for example, the genes in our original filtered table). We can choose our sets from the two drop-down menus in the Enrichment window:

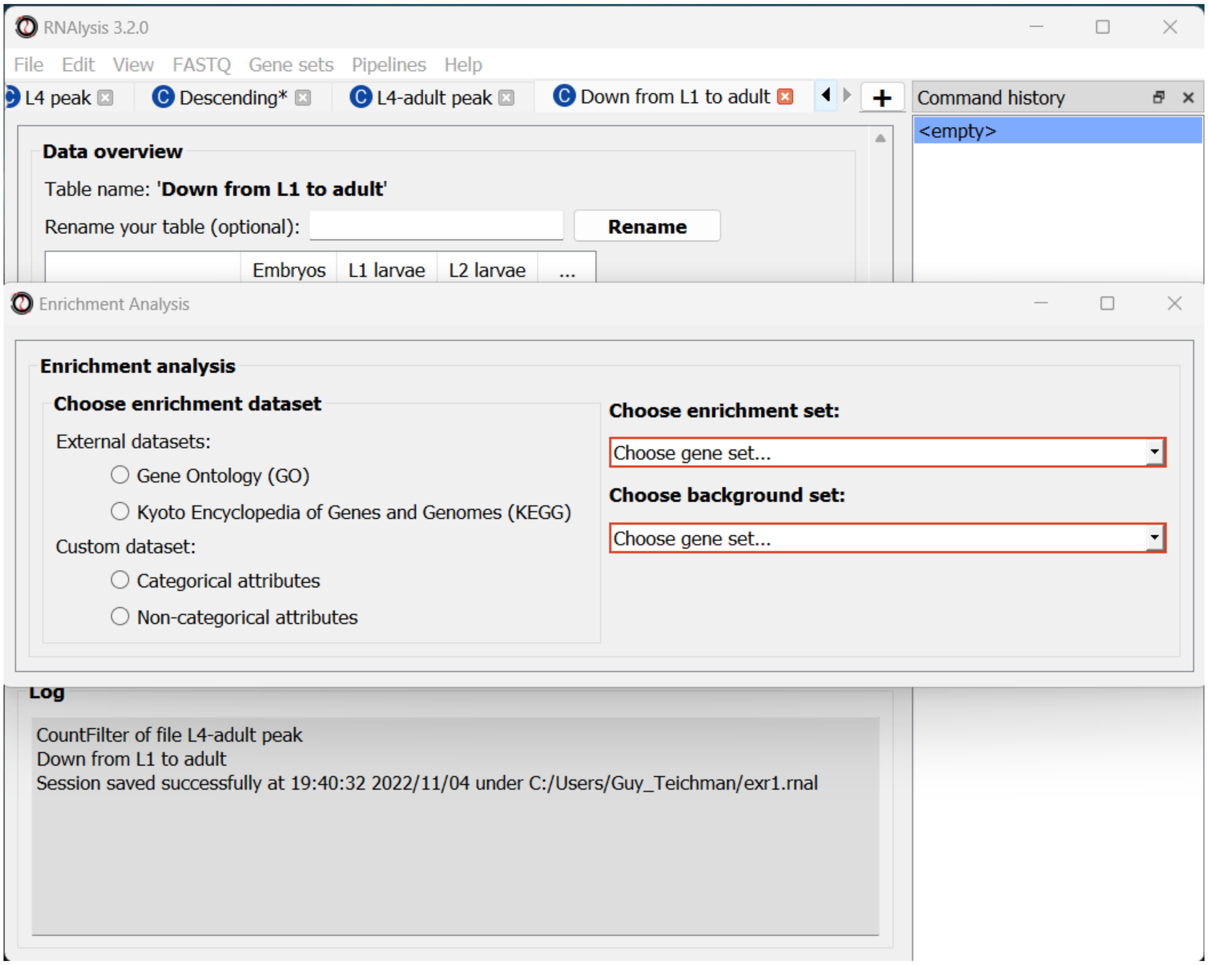


Next, we can choose the dataset we want to draw annotations from. In our case, we will choose Gene Ontology (GO). After picking the dataset, the window expanded to show us all of the parameters we can modify for our analysis:

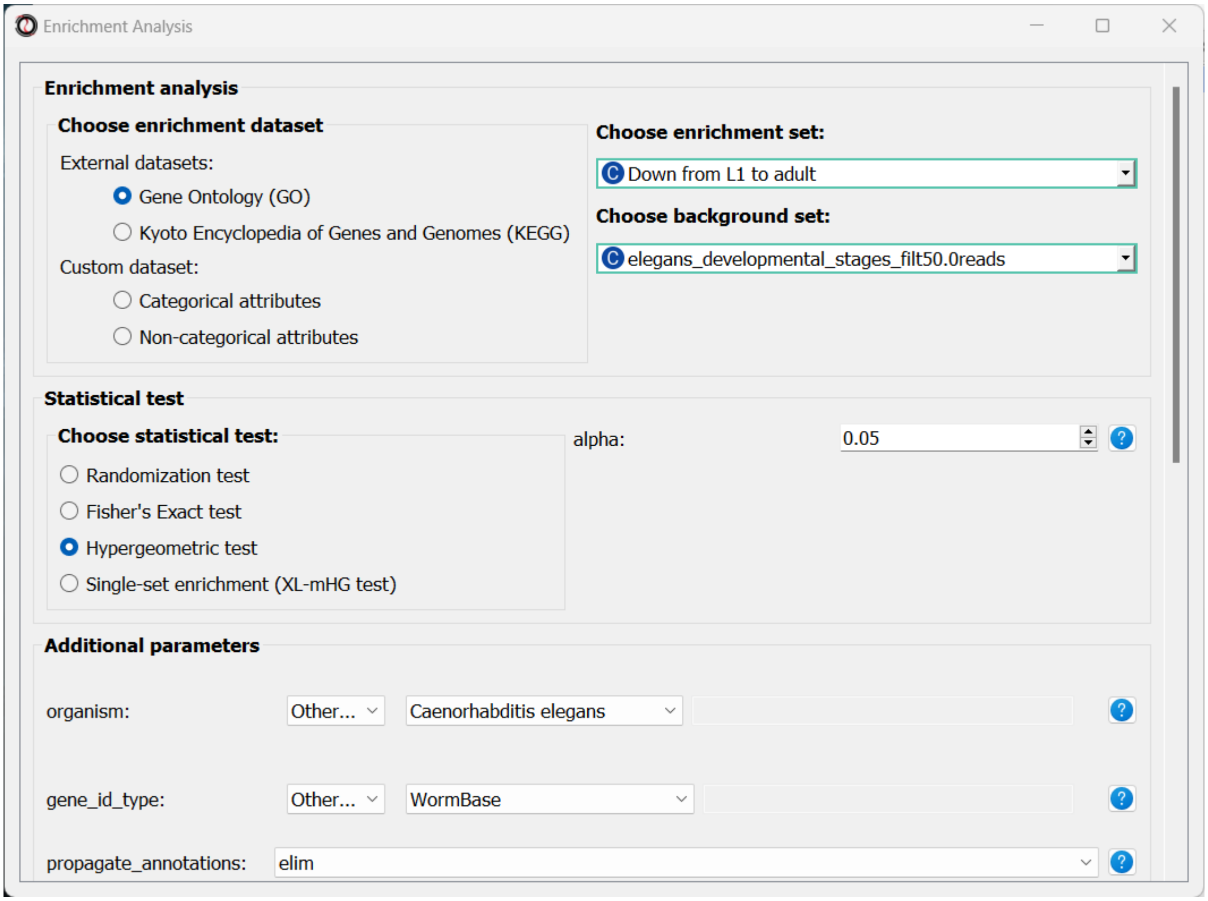


Let's set the statistical test to 'hypergeometric', the organism to 'Caenorhabditis elegans' (matching our gene expression data), and the Gene ID Type to "WormBase" (matching the gene ID type of our original data table).

We will leave the rest of the settings on the default values, but keep in mind that you can customize the analysis to a significant degree: using different statistical tests (including a statistical test that doesn't require a background gene set), using only specific types of GO annotations, propagating the annotations differently, etc'. You can read more about these options in the complete user guide.

Scroll to the bottom of thw window and click on the "run" button to run the analysis:

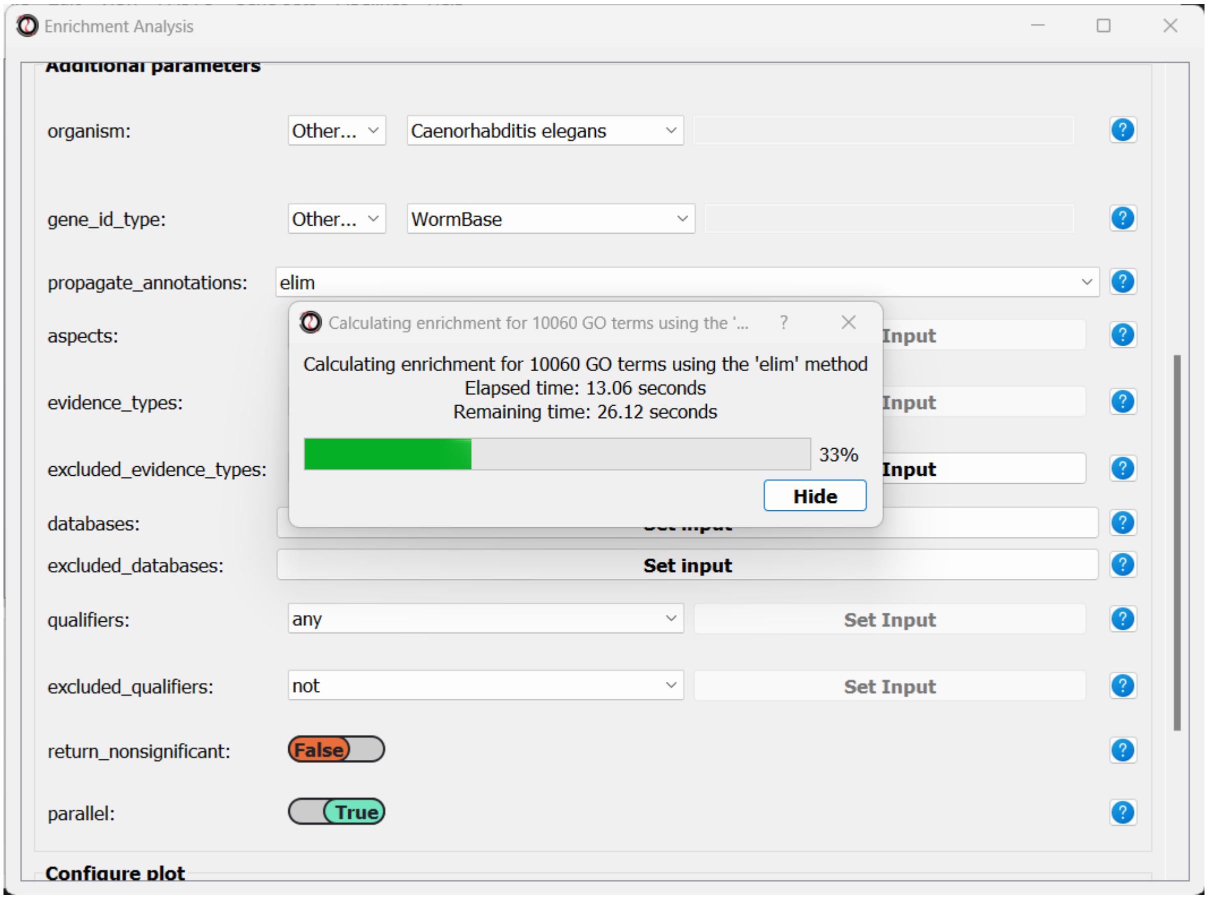


The enrichment window is going to minimize to allow you to read the log box on the main *RNAlysis* window, but you can enlarge it back if you want to look at the analysis parameters, or start a new analysis with the same parameters. The log will update you on the number of annotations found, whether some genes were left entirely unannotated, how many gene IDs were successfully mapped to match the annotations, etc.

Once the analysis is done, we will be able to observe our results in multiple formats.

The first is a tabular format, showing all of the statistically significant GO terms we found (or the all of the tested GO terms, if we set the *return_nonsignificant* parameter to True). The GO terms will be sorted by specificity, the most specific GO terms appearing at the top of the table. The table also includes the statistics for each GO term (number of genes in the cluster, number of genes matching the GO term, expected number of genes to match the GO term, log2 fold change, p-value, and adjusted p-value).

View of 'Enrichment results for set Down from L1 to adult'

Table 'Enrichment results for set Down from L1 to adult': 22 rows, 8 columns

Save table

| | name | samples | obs | exp | log2_fold_enrichment | pval | padj | significant |
|---|---|---|---|---|---|---|---|---|
| GO:0007218 | neuropeptide signaling pathway | 114 | 29 | 1.... | 3.8685561964297155 | 1.... | 1.... | True |
| GO:0005634 | nucleus | 114 | 10 | 31.06179775... | -1.6351413304981093 | 6.... | 0.... | True |
| GO:0043231 | intracellular membrane-bounded... | 114 | 23 | 58.66516853... | -1.3508703203509125 | 6.... | 1.... | True |
| GO:0048522 | positive regulation of cellular ... | 114 | 6 | 12.07247191... | -1.0086867008584899 | 4.... | 0.... | True |
| GO:0005737 | cytoplasm | 114 | 37 | 62.53988764... | -0.7572513565688234 | 1.... | 1.... | True |
| GO:0043227 | membrane-bounded organelle | 114 | 23 | 59.04943820... | -1.3602894726833066 | 3.... | 7.... | True |
| GO:0043229 | intracellular organelle | 114 | 40 | 69.07247191... | -0.7881108545840732 | 1.... | 1.... | True |
| GO:0050794 | regulation of cellular process | 114 | 42 | 30.80561797... | 0.44719585069773654 | 3.... | 0.... | True |
| GO:0016043 | cellular component organization | 114 | 5 | 22.92808988... | -2.1971161706878757 | 1.... | 0.... | True |

The second output format is an Ontology Graph, depicting the statistically-significant GO terms in each GO aspect (biological process, cellular component, and molecular function), as well as their ancestors in the ontology graph. The hierarchy of GO terms in the graph matches the official Gene Ontology hierarchy. The color of the terms on the graph indicates their log2 fold change, and the depth in the tree indicates the specificity of the term, with more specific GO terms being at the bottom.

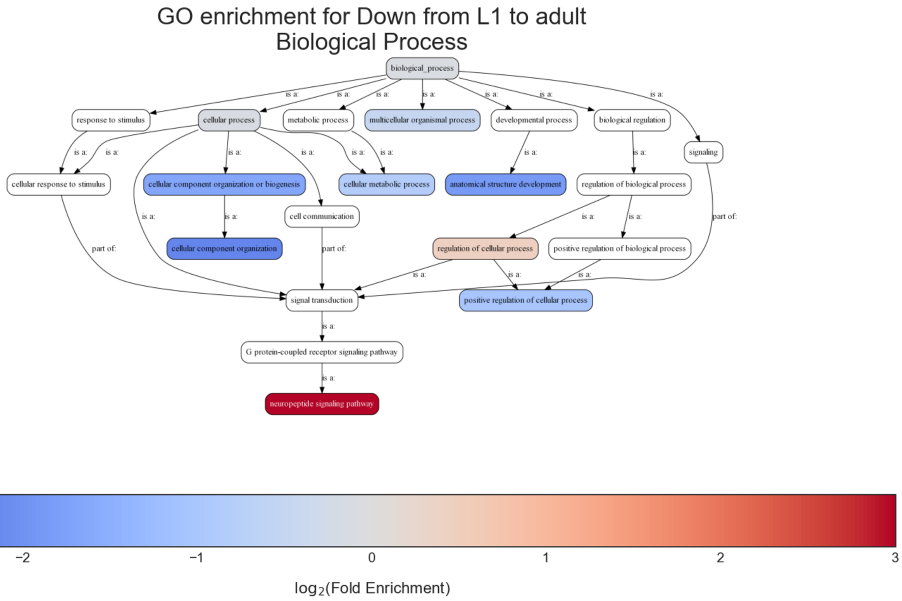


The final output format is a bar plot depicting the log2 fold change values, as well as significance, of the 10 most specific GO terms that were found to be statistically significant in our analysis.

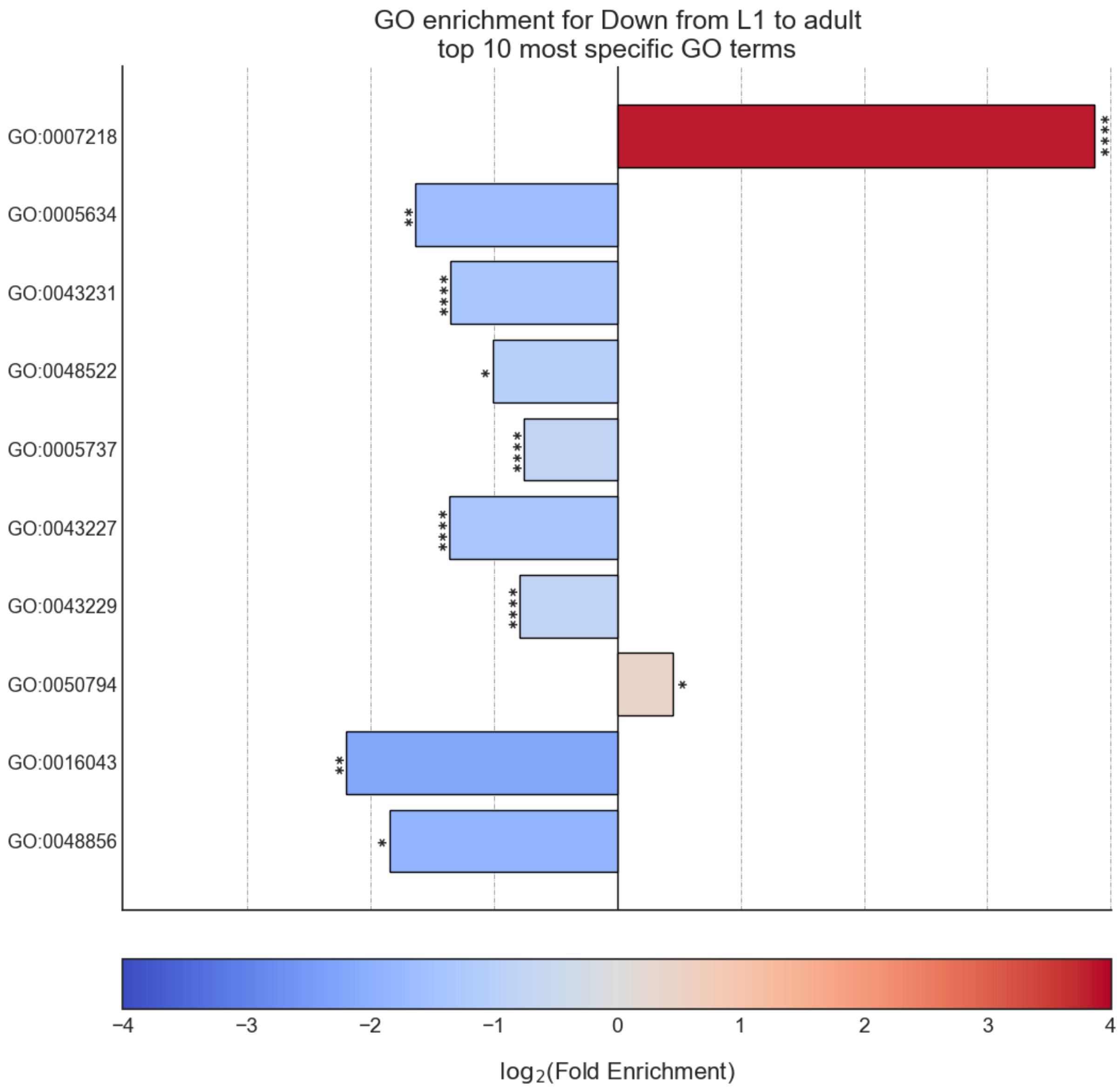


### Analysis #2 - differential expression data

#### The dataset

In the second analysis, we will analyze three publicly available datasets from GEO. Each individual dataset is an RNA sequencing experiment of *Caenorhabidtis elegans* nematodes, measuring the effect of a particular stress condition on gene expression. Our goal in this analysis will be to examine the similarities between the different stress conditions, and to answer a specific question - how are Epigenetic genes and pathways affected by exposure to stress?

In this analysis, unlike the first analysis, we will not start from a pre-made counts table. Instead, we are going to pre-process and quantify each dataset directly from the raw FASTQ files that are available on GEO.

The datasets we will use are:

1. **mRNA sequencing of worms exposed to osmotic stress: GSE107704. This is a single-end unstranded RNA sequencing experiment.**

    - Control sample 1 - SRR6348197
    - Control sample 2 - SRR6348198
    - Control sample 3 - SRR6348199
    - Osmotic sample 1 - SRR6348200
    - Osmotic sample 2 - SRR6348201
    - Osmotic sample 3 - SRR6348202

2. **mRNA sequencing of worms exposed to starvation: GSE124178. This is a paired-end stranded RNA sequencing experiment.**

    - Control sample 1 - SRR8359132
    - Control sample 2 - SRR8359133
    - Control sample 3 - SRR8359142
    - Starvation sample 1 - SRR8359134
    - Starvation sample 2 - SRR8359135
    - Starvation sample 3 - SRR8359136

3. **mRNA sequencing of worms exposed to heat shock: GSE132838. This is a paired-end stranded RNA sequencing experiment.**

    - Control sample 1 - SRR9310678
    - Control sample 2 - SRR9310679
    - Control sample 3 - SRR9310680
    - Heat sample 1 - SRR9310681
    - Heat sample 2 - SRR9310682
    - Heat sample 3 - SRR9310683

#### Quantify FASTQ files and Differential Expression Analysis

RNAlysis offers user the opportunity to start analysis right from raw FASTQ data, using XYZ tools.

Since our input data is raw FASTQ files, we will first have to pre-process them, quantify them, and then detect differentially expressed genes. When using RNAlysis, you can start analyzing your data at any point in the analysis pipeline: raw FASTQ files, trimmed FASTQ files, quantified read counts, or differential expression tables. This means you can combine *RNAlysis* with any other tools or pipelines you prefer.

*RNAlysis* provides a graphical interface for *CutAdapt*, *kallisto*, and *DESeq2*, and we will use those tools in this tutorial analysis.

#### Quality control with *FastQC*

The first step in analyzing RNA sequencing data is uaually quality control. This can help us get a general overview of our FASTQ files, and detect any potential issues with them. One of the most popular tools for this purpose is FastQC. FastQC is an open-source tool, and it has both a friendly graphical interface and a programmatic API. You can get it here.

#### Trim adapters and remove low-quality reads with *CutAdapt*

After doing a quality-control of our FASTQ files with FastQC, we can see that some of our samples had a small portion of reads that still contain adapter sequences. Therefore, we will start our analysis by trimming the leftover adapters. We will also perform quality-trimming, removing bases with low quality scores from our reads.

The first dataset we will trim is the osmotic stress dataset, which happens to be single-end sequencing. To begin, let's open the "FASTQ" menu in *RNAlysis*, and under "Adapter trimming" choose "Single-end adapter trimming":

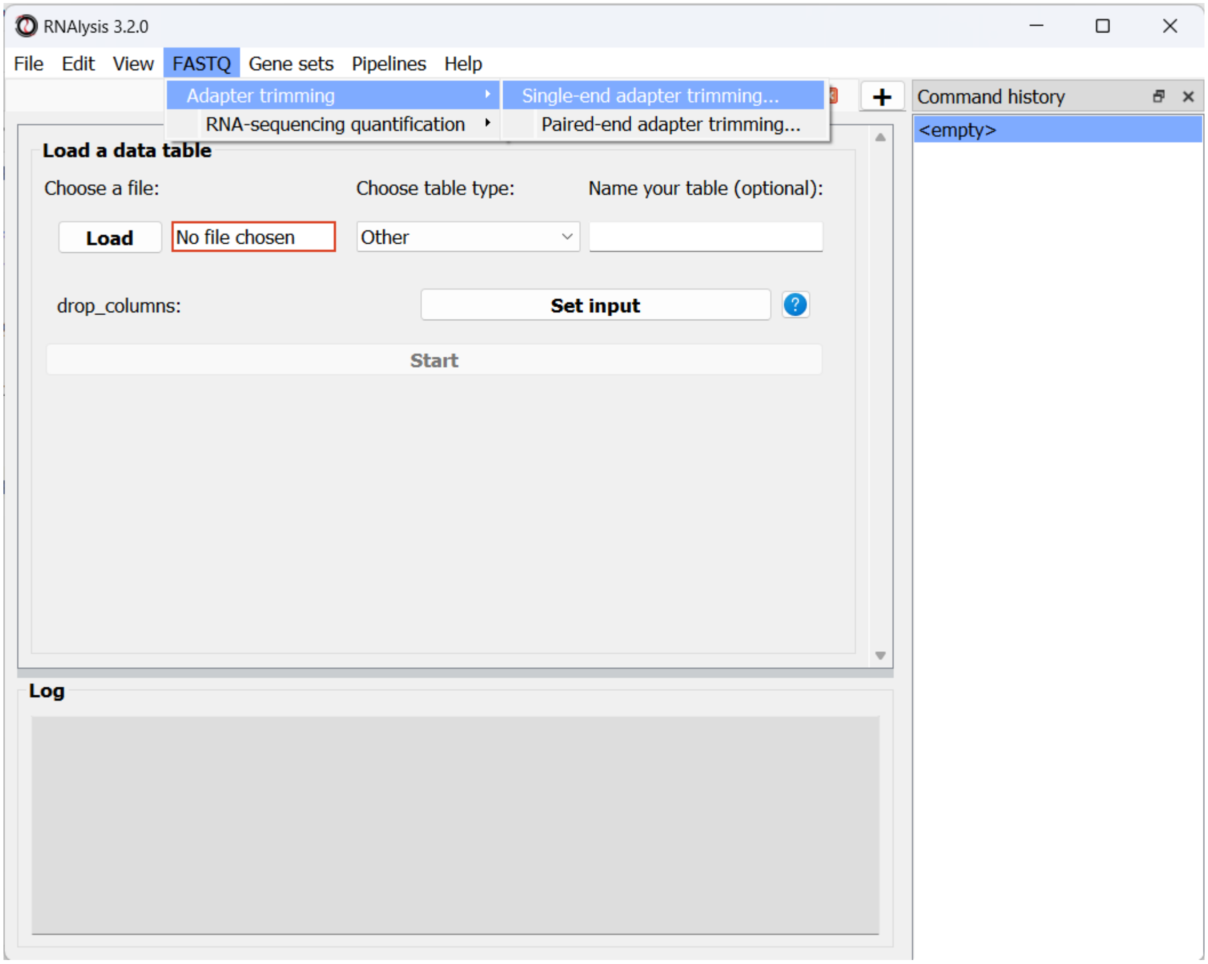


In the new window that opened, we can choose the folder that contains our raw FASTQ files, as well as the folder our trimmed files should be written to:

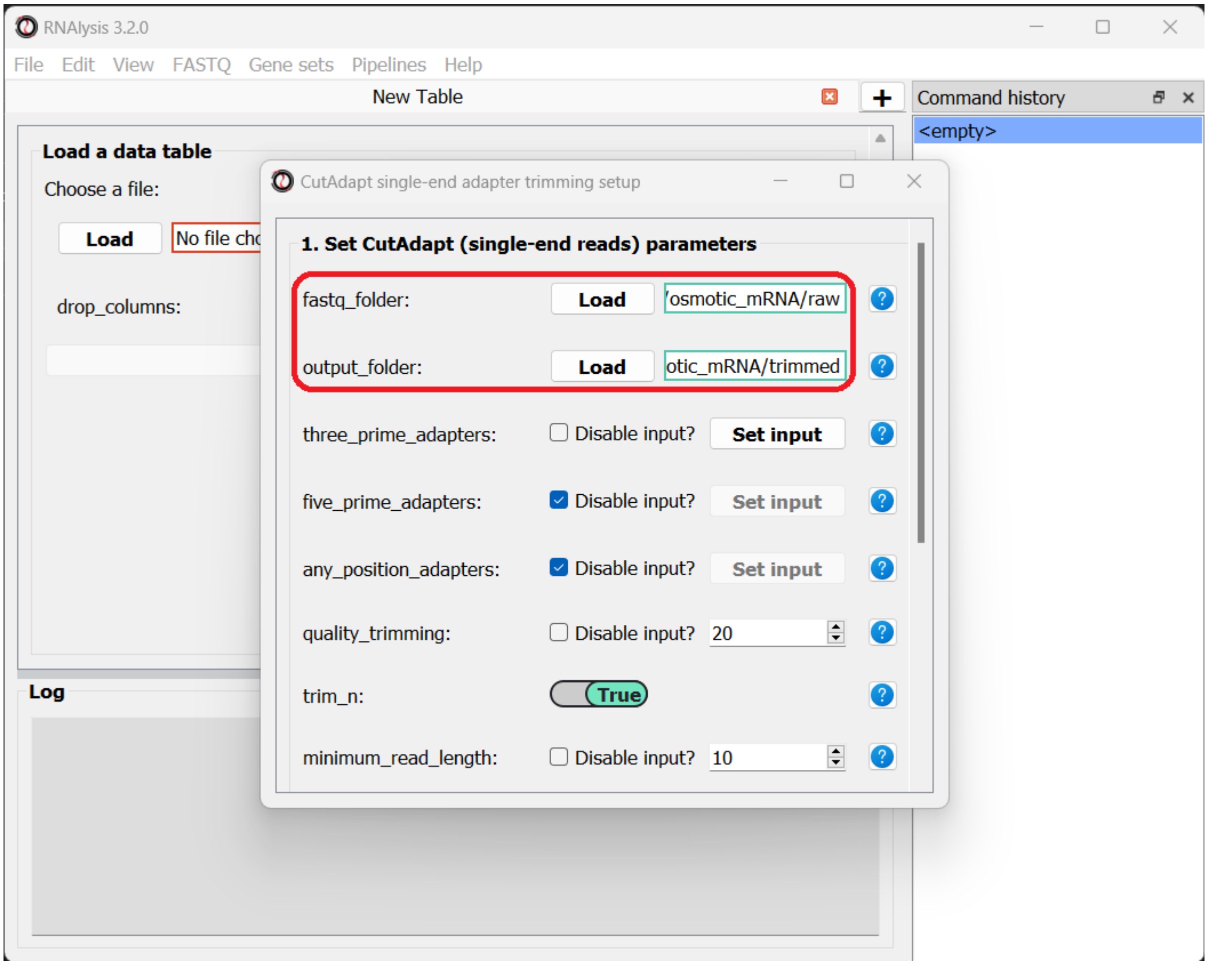


The input folder can contain any number of raw FASTQ files, and *RNAlysis* will trim them one by one and save the trimmed output files in the output folder.

Next, let's set the adapter sequence we want to trim. The adapter used on these samples is the three-prime Illumina TruSeq adapter, with the sequence "AGATCGGAAGAGCACACGTCTGAACTCCAGTCA". Let's click on the "Set input" button of the *three_prime_adapters* parameter:

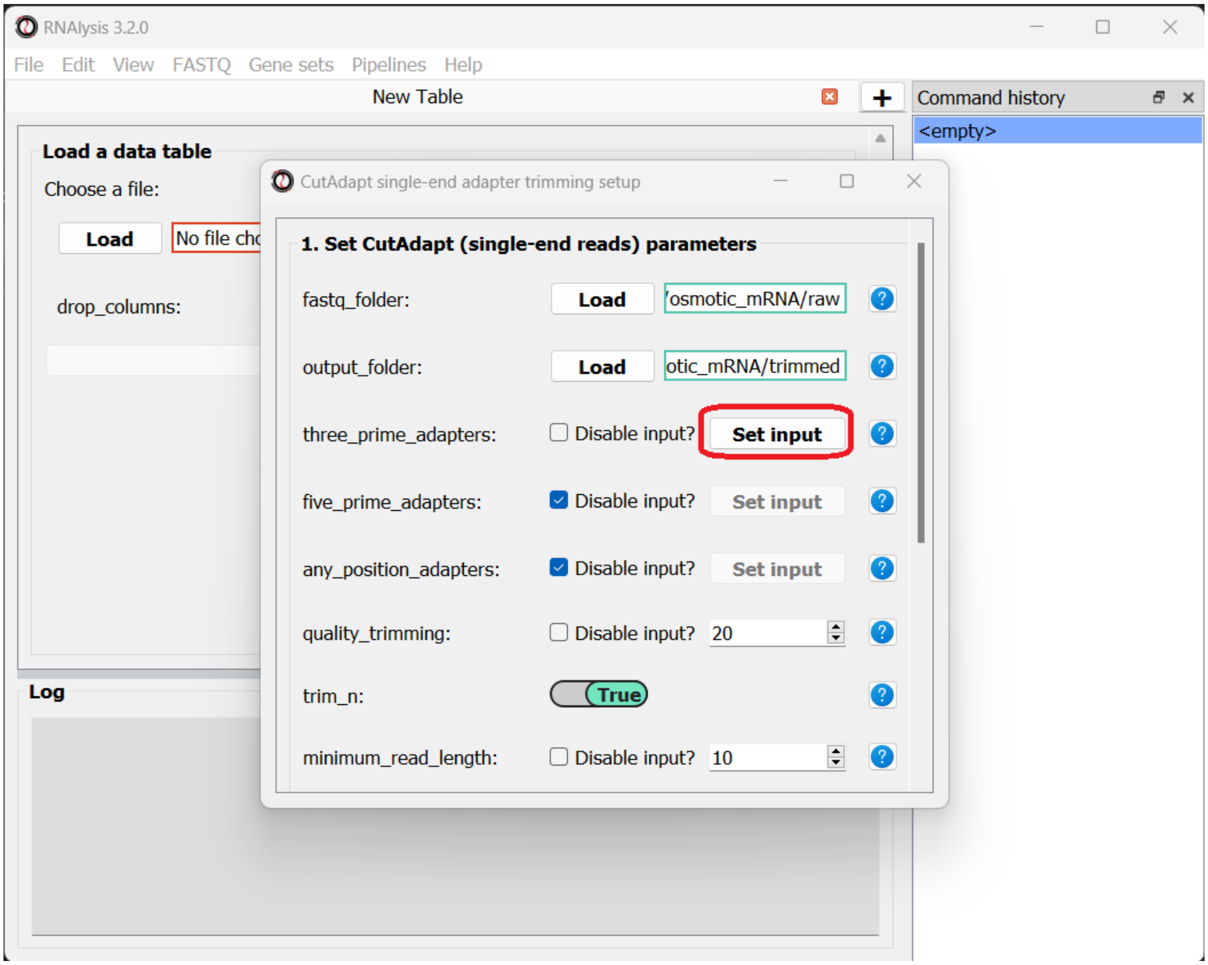


This will open a dialog box, where we can enter the sequence of our adapter:

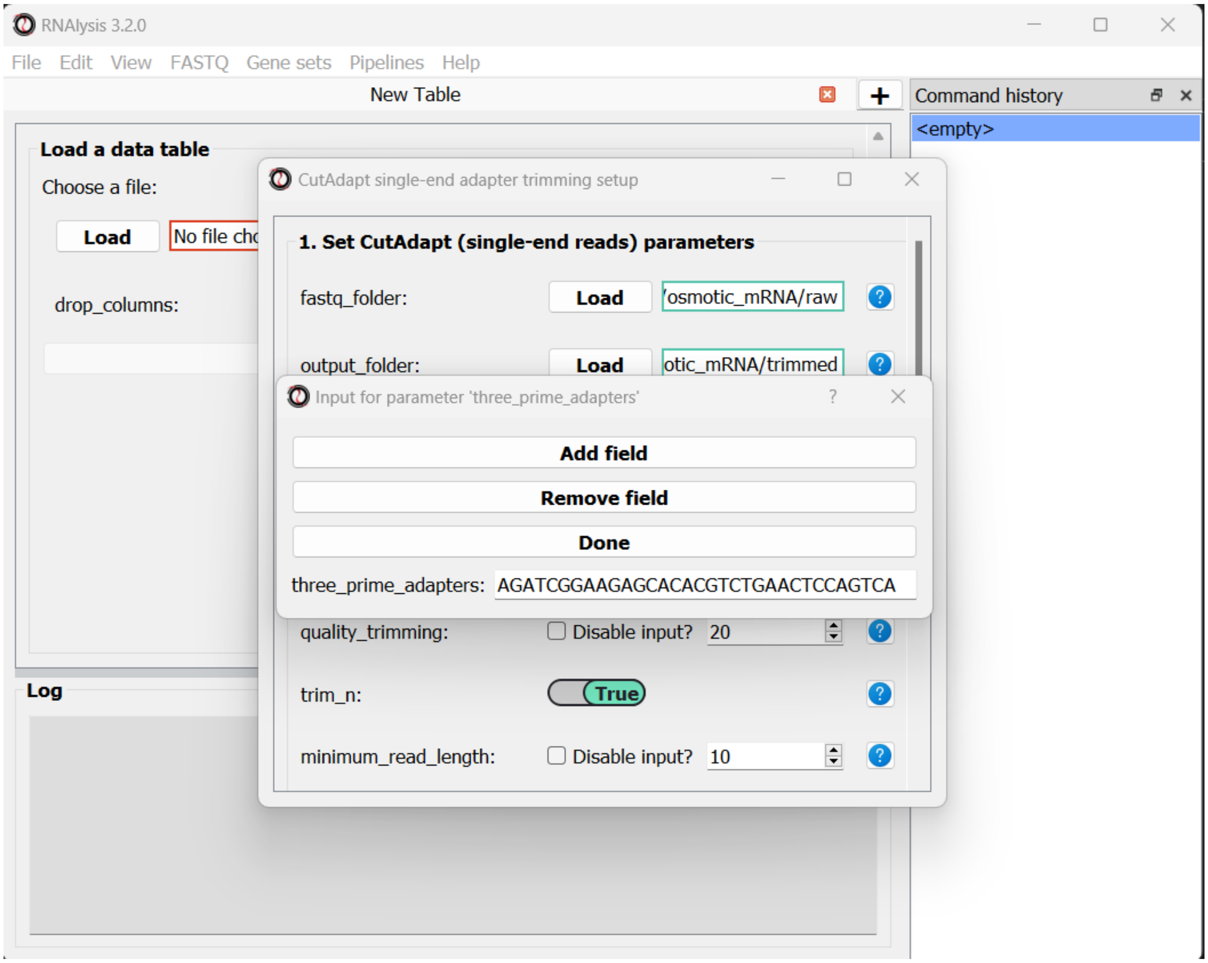


We will leave the quality-trimming parameters on their default values - trim bases below quality score 20 from the 3’ end of reads, trim flanking N bases from the reads, and filter out any read that ends up shorter than 10bp after trimming:

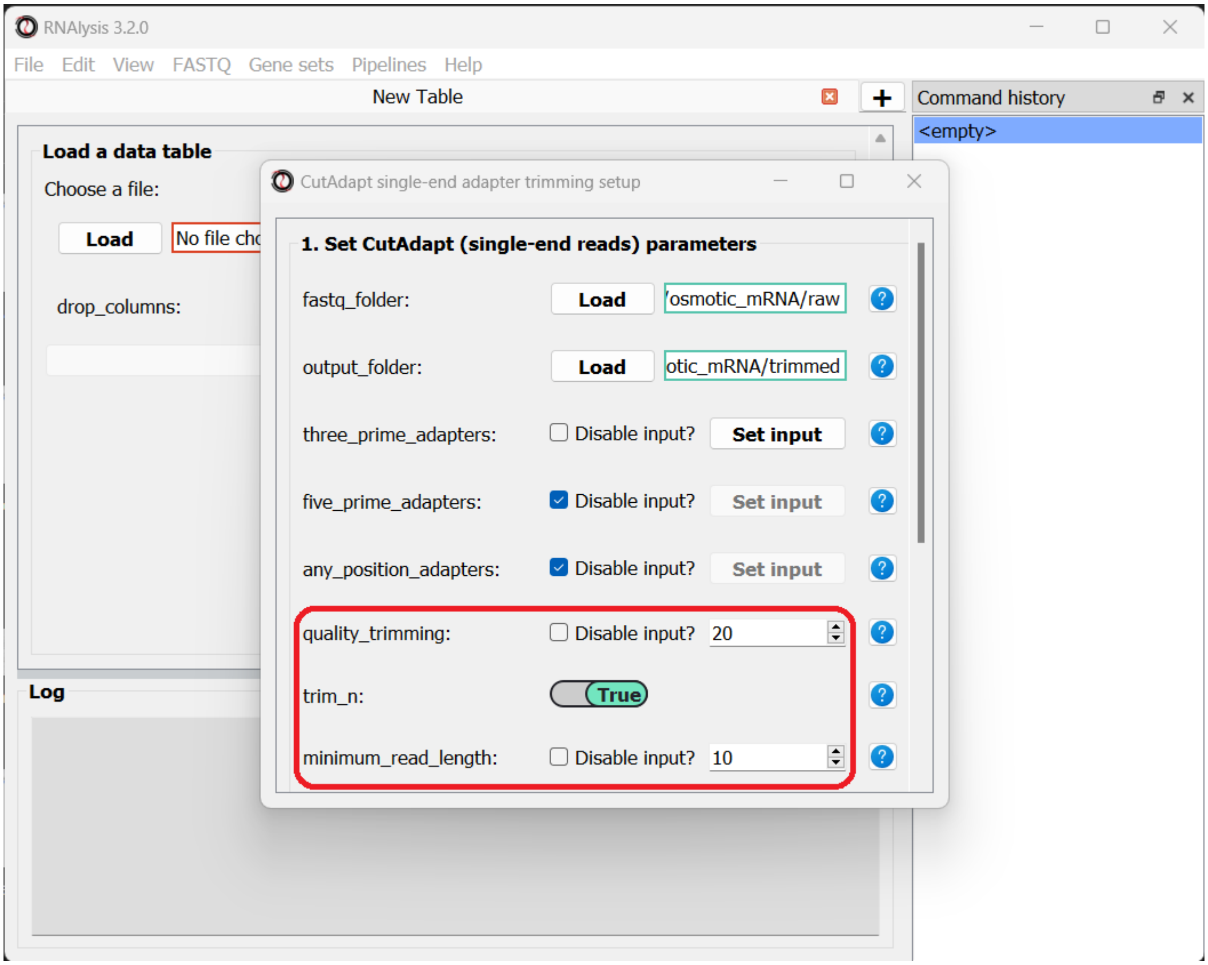


Let's now scroll to the bottom of the window and set the *discard_untrimmed_reads* parameter to False, so that *CutAdapt* will not discard reads that were previously trimmed properly. Usually we do want to discard any reads that didn't contain adapter sequences, but in our case, most of the reads were already trimmed, so we don't want to throw them away.

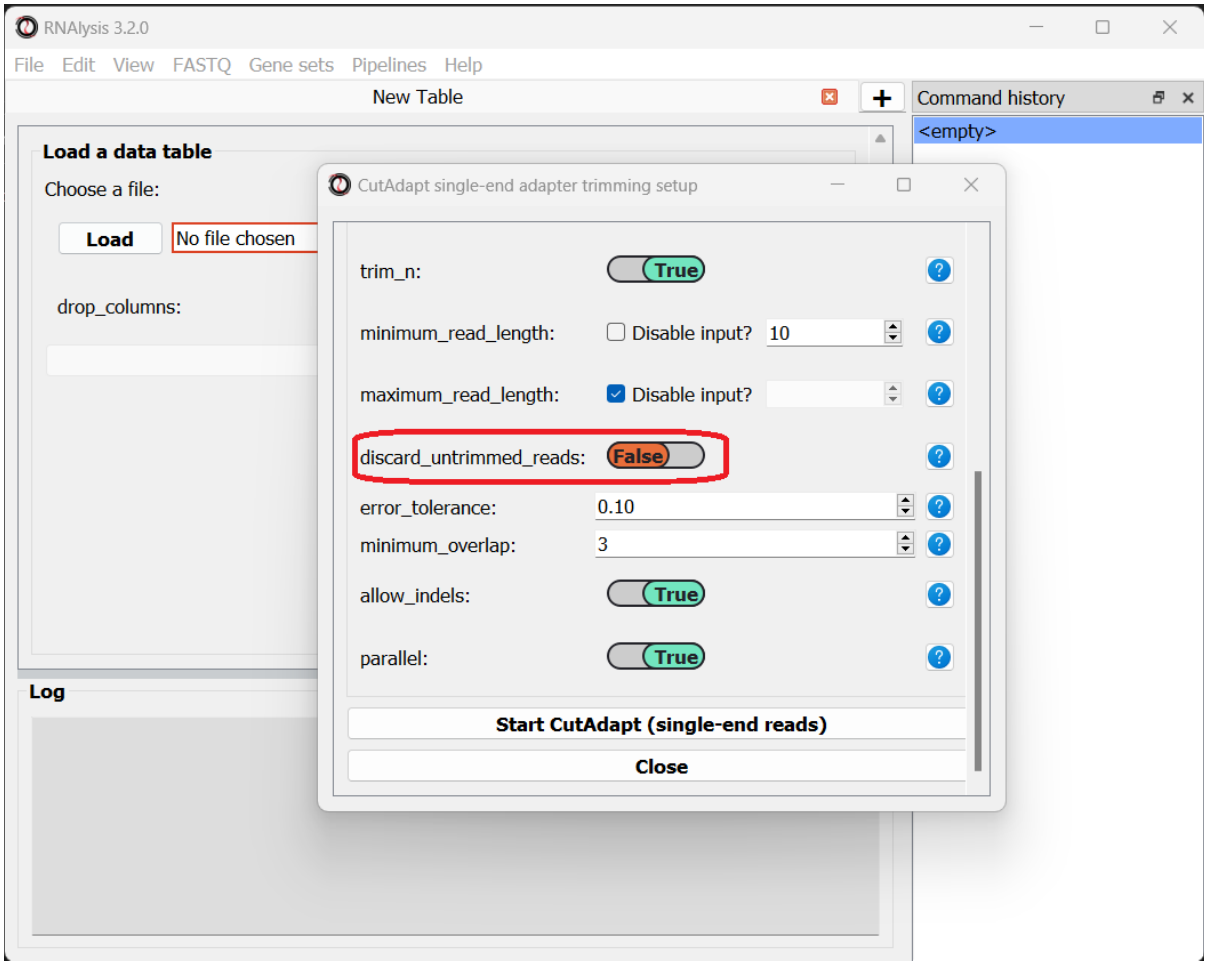


Once we are happy with the trimming parameters, we can click on the "Start CutAdapt" button at the bottom of the window. A loading screen will now appear, and the trimmed FASTQ files, as well as the trimming logs, will be saved to our output folder.

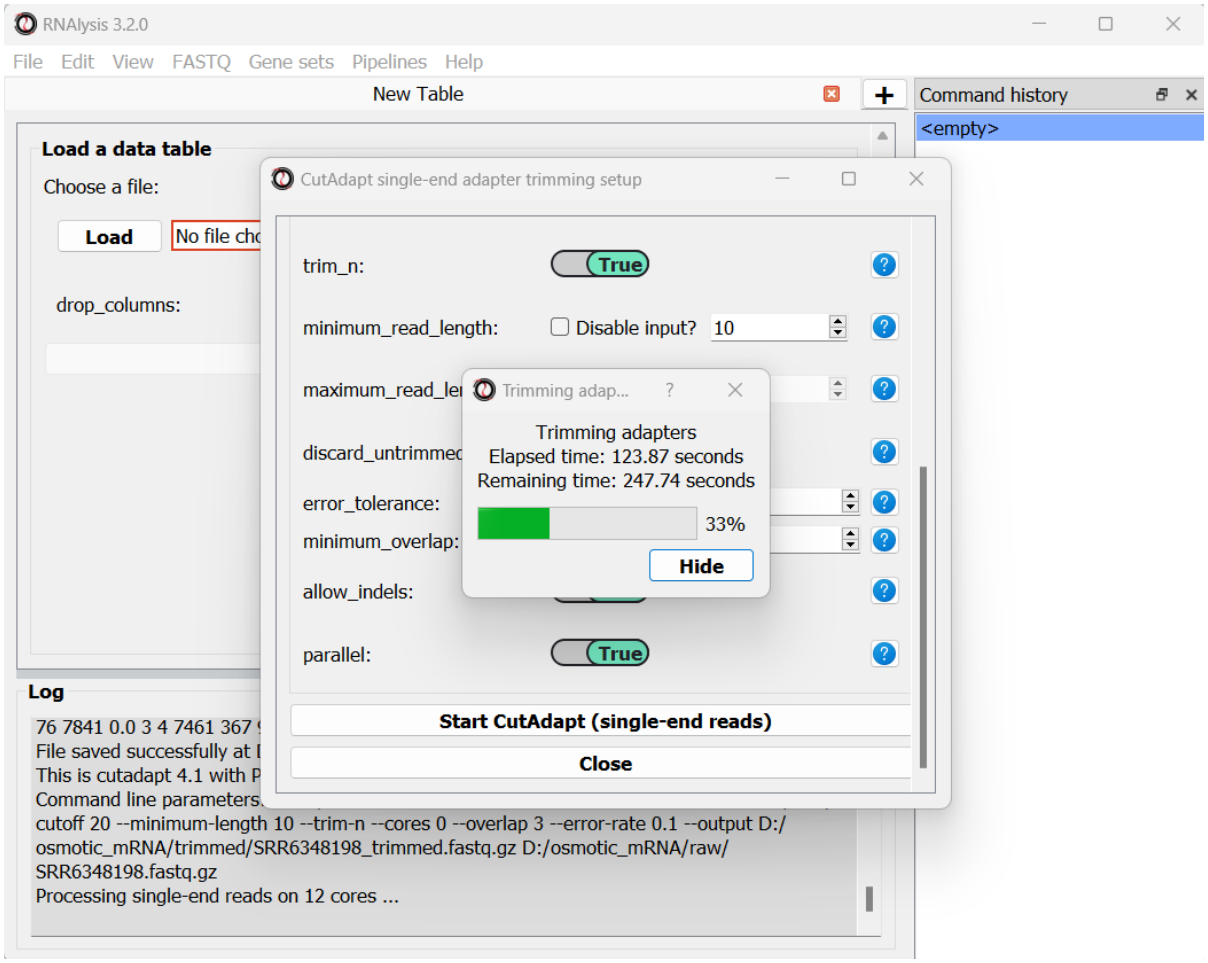


Let’s now apply the same trimming procedure to our paired end samples. Open the Paired End adapter trimming from the FASTQ menu:

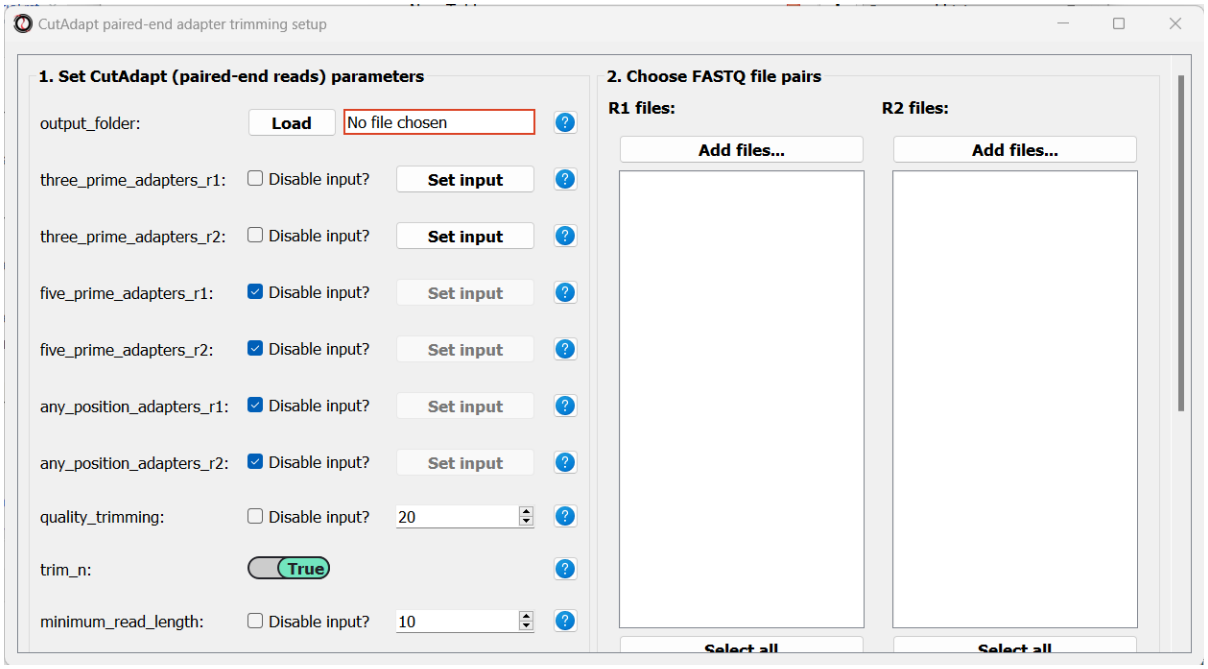


First, we will set the output folder for our trimmed FASTQ files. Then, we can set the adapter sequences: for read#1 we will use the TruSeq Read 1 adapter “AGATCGGAAGAGCACACGTCTGAACTCCAGTCA”, and for read#2 we will use the TruSeq Read

2 adapter "AGATCGGAAGAGCGTCGTGTAGGGAAAGAGTG". We will leave the quality-trimming parameters the same as last time, and once again set the *discard_untrimmed_reads* parameter to False:

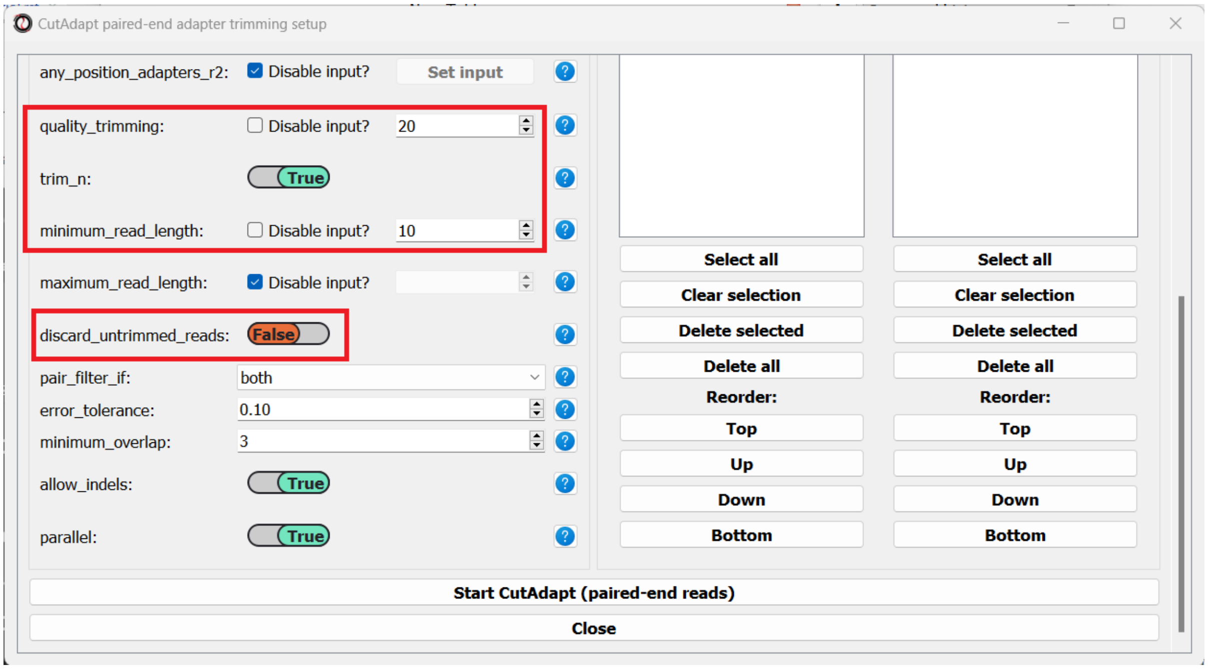


Finally, we can choose the files we want to trim. Since we are doing paired-end trimming, we have to make sure the files are properly paired. Therefore, we will pick the read#1 and read#2 files separately by clicking on the "Add files" buttons:

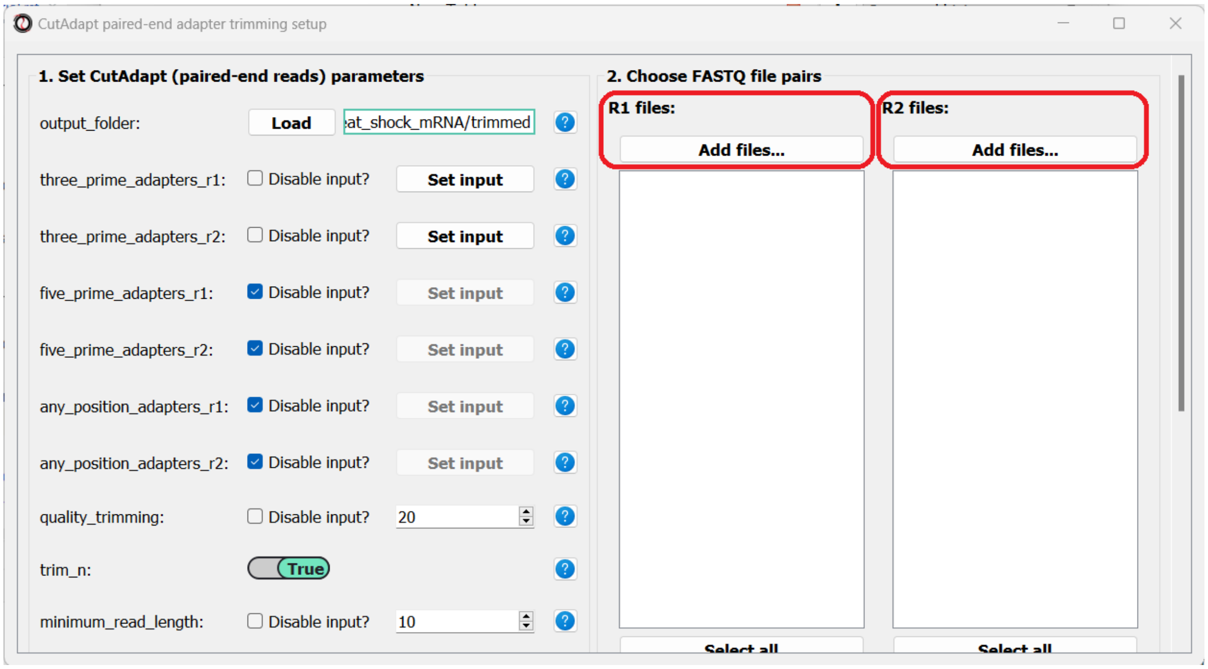


After adding the files, we make sure the orders of the two file lists match, so that the files are properly paired:

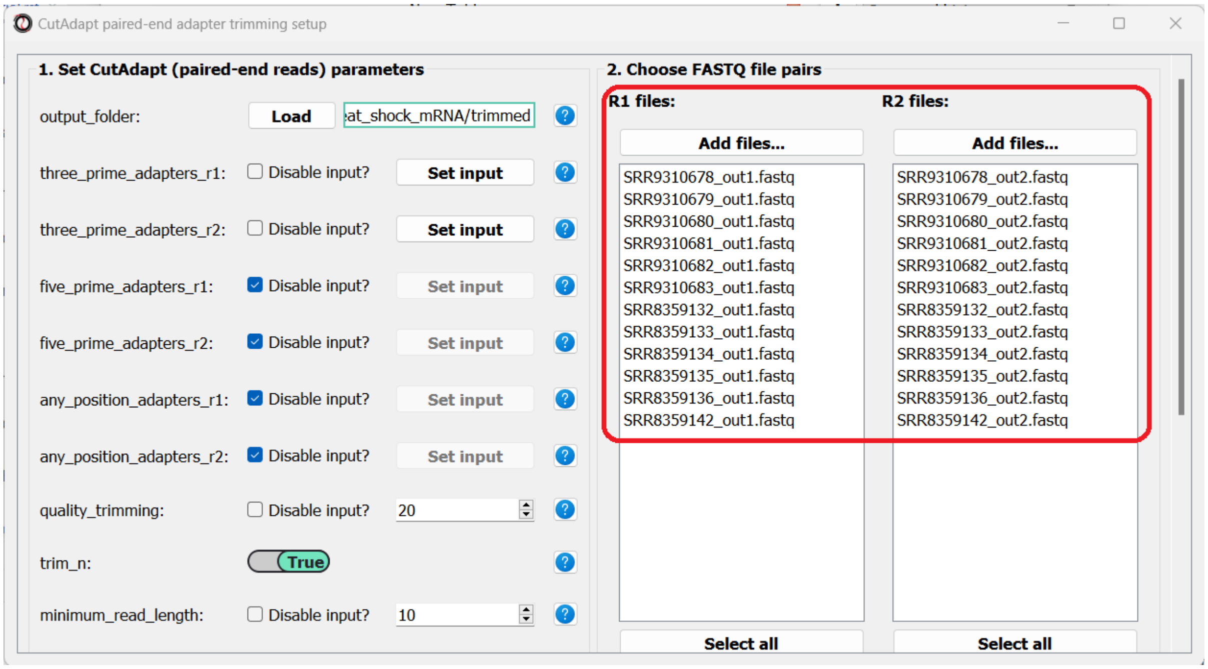


You can use the buttons at the bottom of each list to reorder the list, or remove some of the files you added:

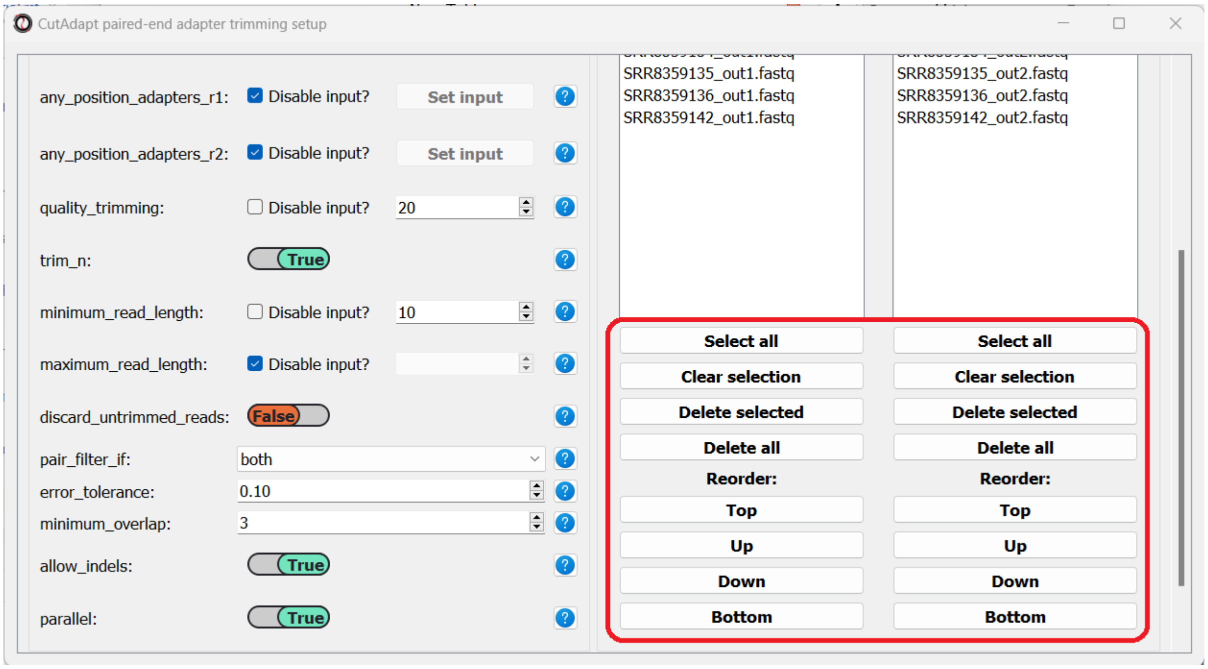


Once we are done setting up the trimming parameters, we can scroll down and click on the "Start CutAdapt" button.

#### Quantify gene expression with *kallisto*

Now that out data has been pre-processed, we can proceed with quantifying the expression of each gene. For this purpose we will use kallisto - a program for rapid quantification of transcript abundances. *kallisto* uses a method called **pseudoalignment** to directly estimate the number of reads originating from each transcript in our target transcriptome, without actually aligning the reads to an exact physical position on the genome/transcriptome. This means that alignment with *kallisto* is extremely quick, while providing results that are as good as traditional alignment methods. Before proceeding with this step, make sure you have installed kallisto on your computer.

To run this analysis, in addition to our processed FASTQ files, we will need a transcriptome indexed by *kallisto*, and a matching GTF file describing the names of the transcripts and genes in the transcriptome. *kallisto* provides pre-indexed transcriptomes and their matching GTF files for most common model organisms, which can be downloaded here. Our data was sequenced from *Caenorhabditis elegans* nematodes, so we will download the *C. elegans* index and GTF files from the above link. If your experiment requires a different transcriptome that is not available above, you can index any FASTA file through *RNAlysis*, by entering the "FASTQ" menu -> "RNA sequencing quantification" -> "Create kallisto index...".

As we did earlier with adapter trimming, let's begin quantification with our single-end osmotic stress samples. Open the "FASTQ" menu, and under "RNA sequencing quantification" select "Single-end RNA sequencing quantification...":

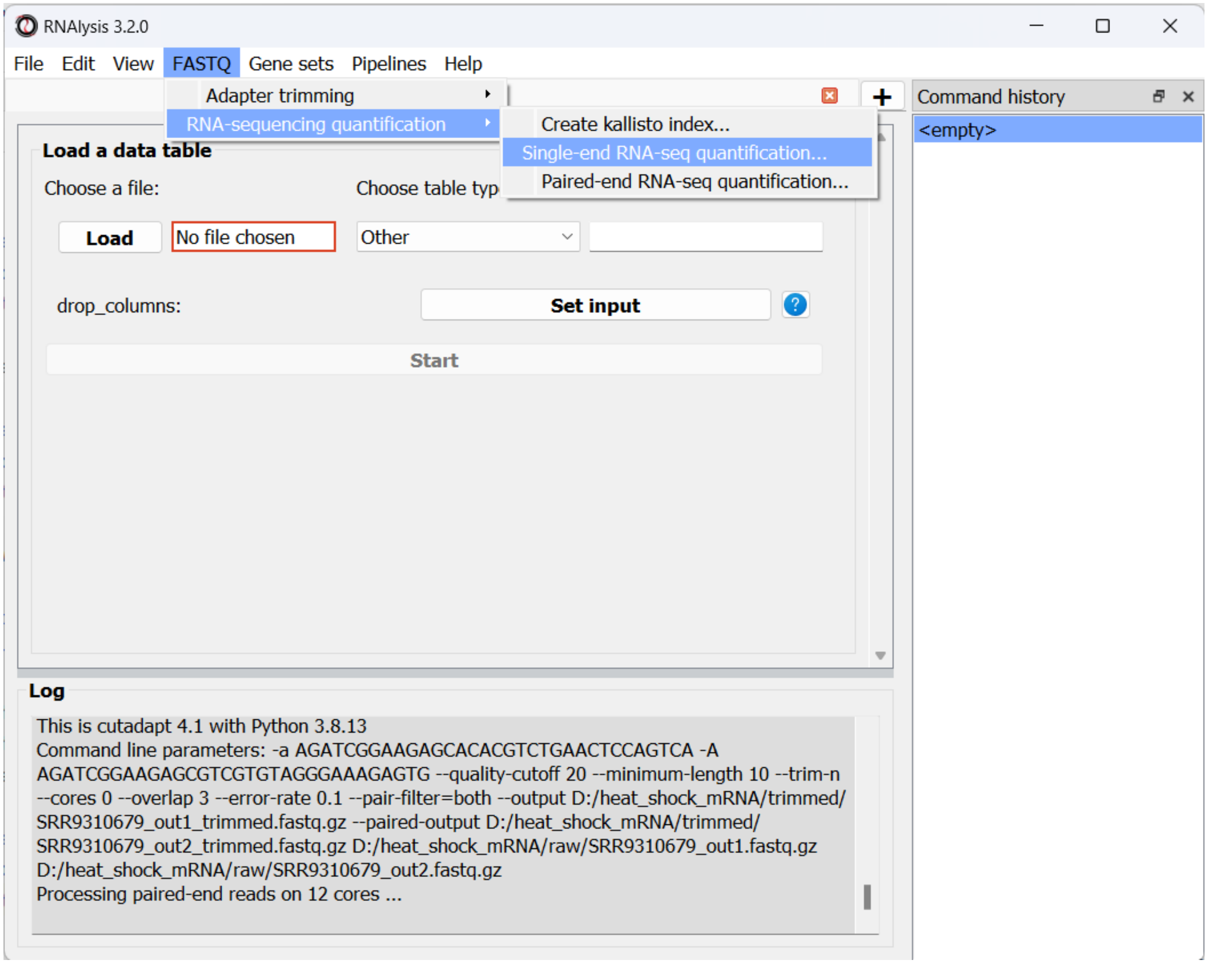


Like before, in the new window that opened we can set the input folder containing our trimmed FASTQ files, and the output folder. We should also set the path to our transcriptome index file, GTF file, and the folder in which you installed *kallisto* (unless you have also added it to your system's PATH):

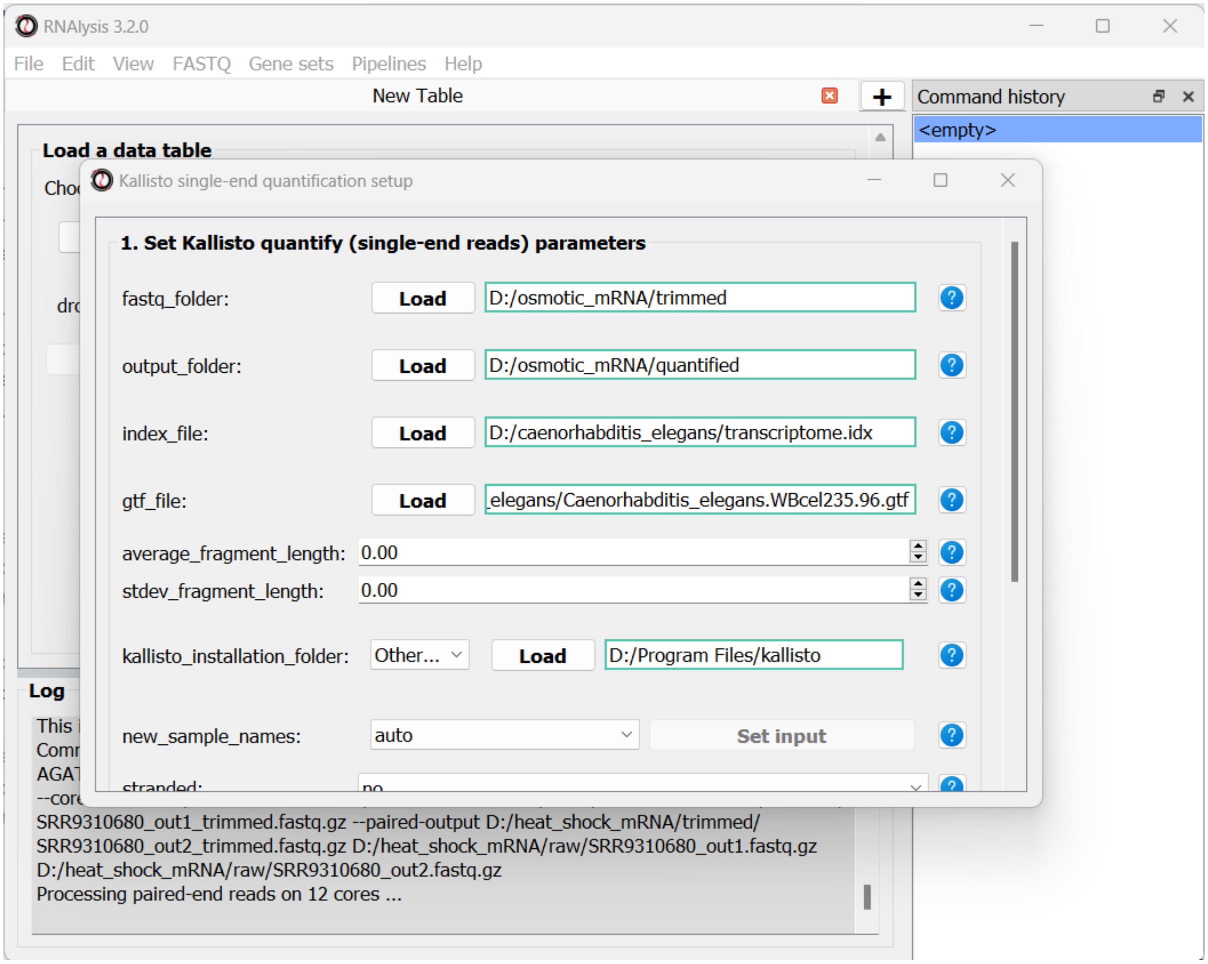


Since our data were sequenced with single-end reads, *kallisto* cannot estimate the size of the fragments in the sequencing run, so we will have to supply an estimate of the average fragment length and the standard deviaton. Let's set them to 200 and 20 accordingly:

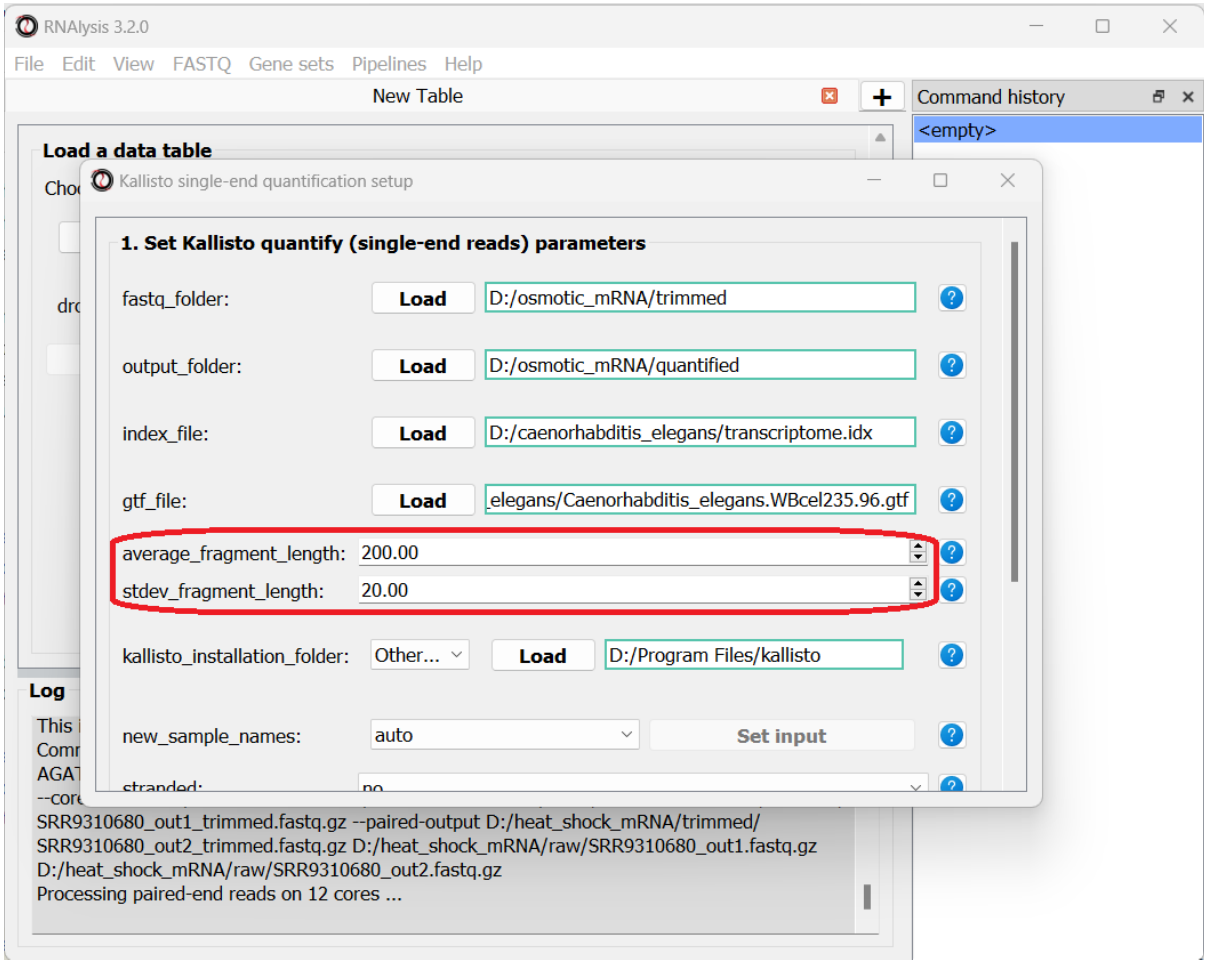


We can optionally use the *new_sample_names* parameter to give our samples new, more readable names. For example:

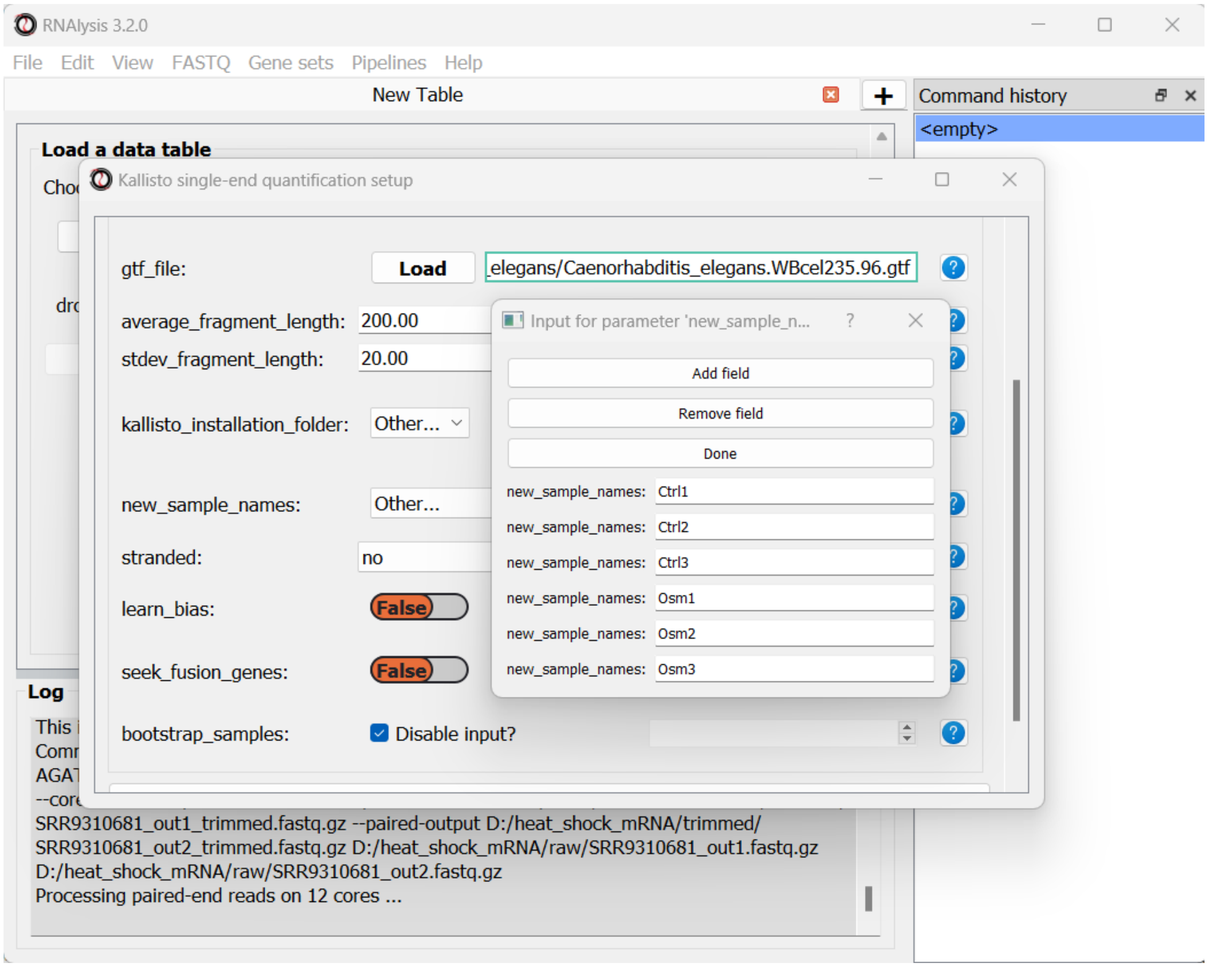


Once we are done setting up the quantification parameters, we can scroll down and click on the "Start kallisto quantify" button, then wait for the analysis to finish. In the output folder, we can find the results of kallisto quantification for each of the individual FASTQ files, each in its own sub-folder. Alongside these files, we can find three CSV files (comma-separated values, can be opened with Microsoft Excel or with RNAlysis): a per-transcript count estimate table, a per-transcript TPM (transcripts per million) estimate table, and a per-gene scaled output table.

The per-gene scaled output table is generated using the *scaledTPM* method (scaling the TPM estimates up to the library size) as described by Soneson et al 2015 and used in the tximport R package. This table format is considered un-normalized for library size, and can therefore be used directly by count-based statistical inference tools, such as DESeq2, for differential expression analysis later on. *RNAlysis* will load this table automatically once the analysis is finished, and we can see it in a new tab of the main window:

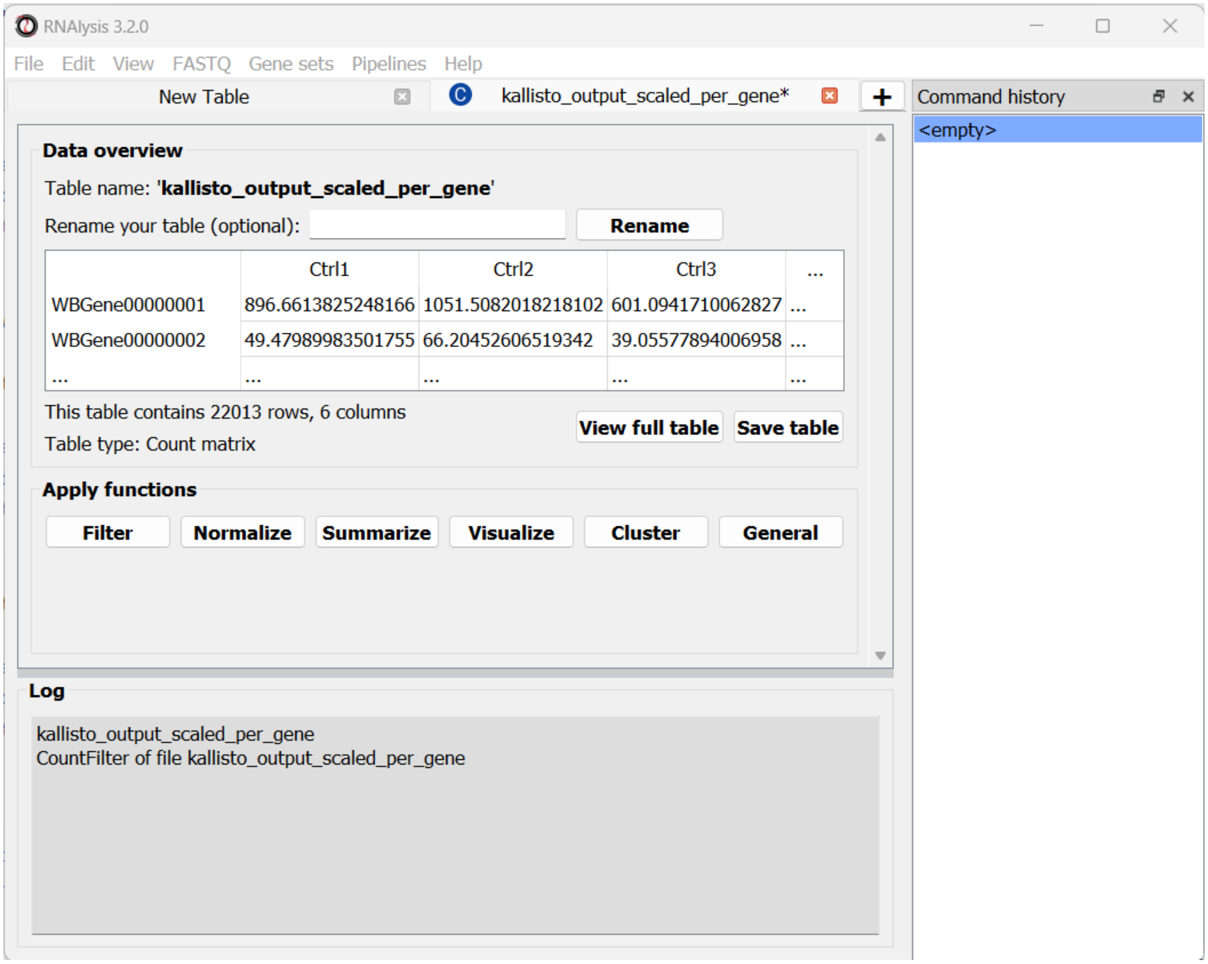


At the bottom of the window we can also see a log box - this is where *RNAlysis* will display notes and warnings regarding our analysis.

If you wanted to analyze differential expression for transcripts instead of genes, you could use the per-transcript count estimate table that is located in the output folder.

Let's now apply the same quantification procedure to our paired-end starvation samples. Open the Paired end quantification window from the FASTQ menu:

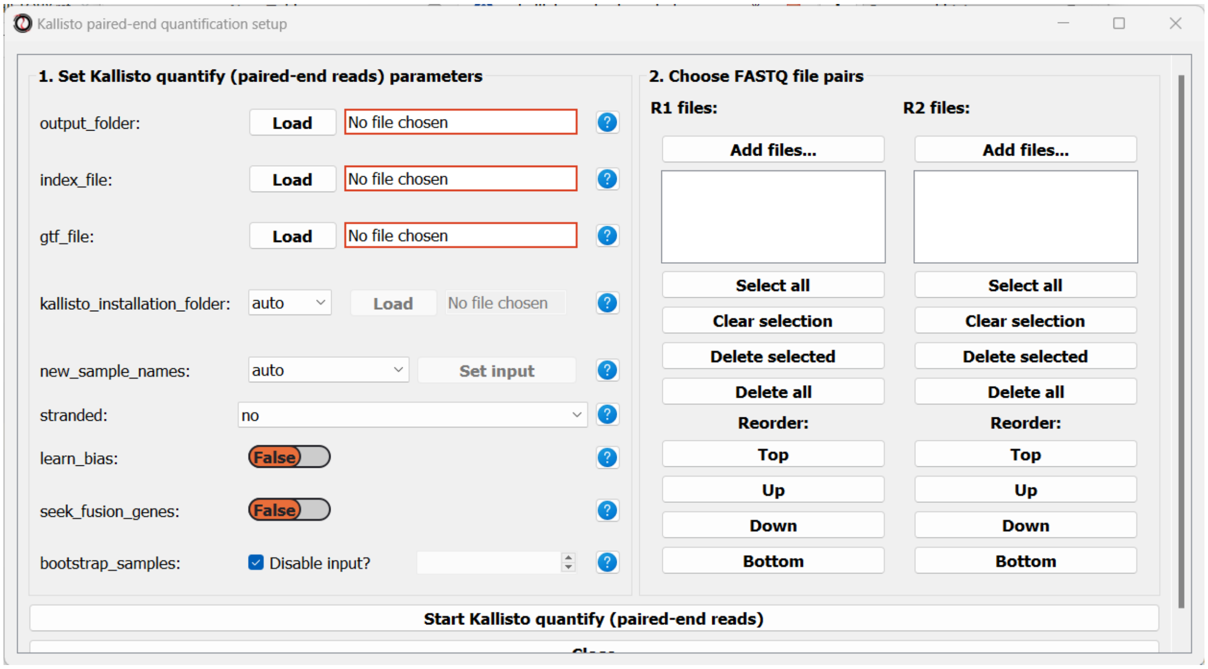


Like before, we will set the path to our output folder, transcriptome index file, GTF file, and the folder in which you installed *kallisto* (unless you have also added it to your system's PATH). In addition, since these sequencing samples are Stranded, we will set the *stranded* parameter to "forward":

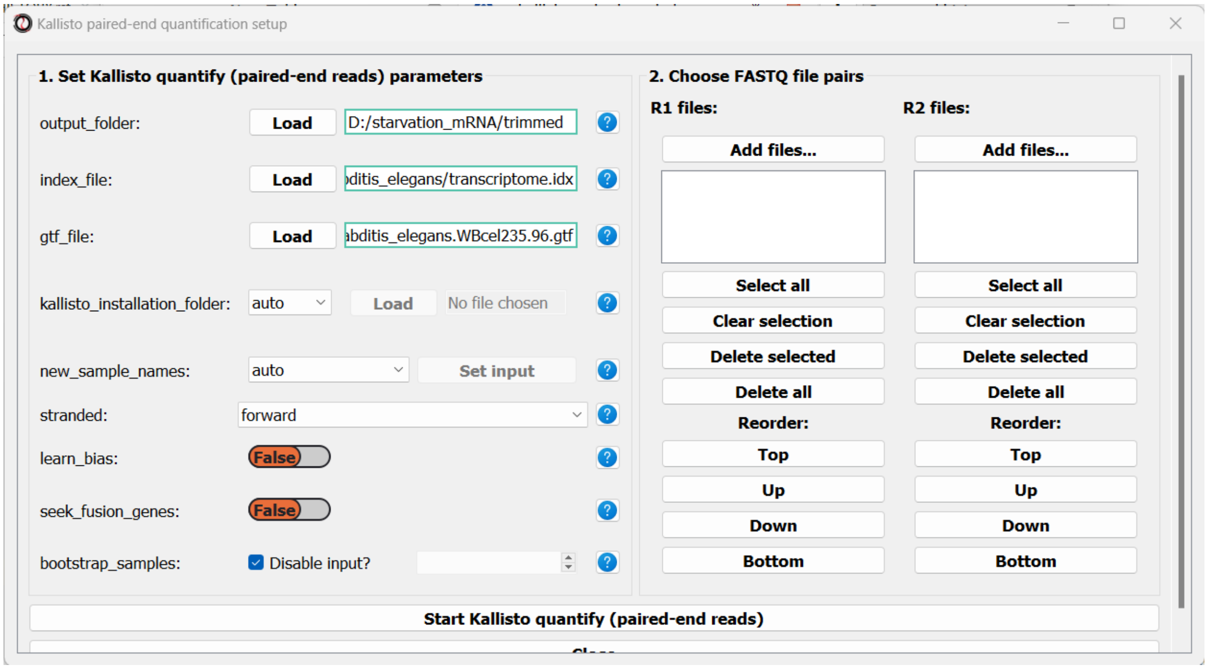


Next, we can choose the files we want to quantify. Like we did when trimming paired-end adapters, we will pick the read#1 and read#2 files separately by clicking on the "Add files" buttons underneath each list, and then sort the files so the pairs match up:

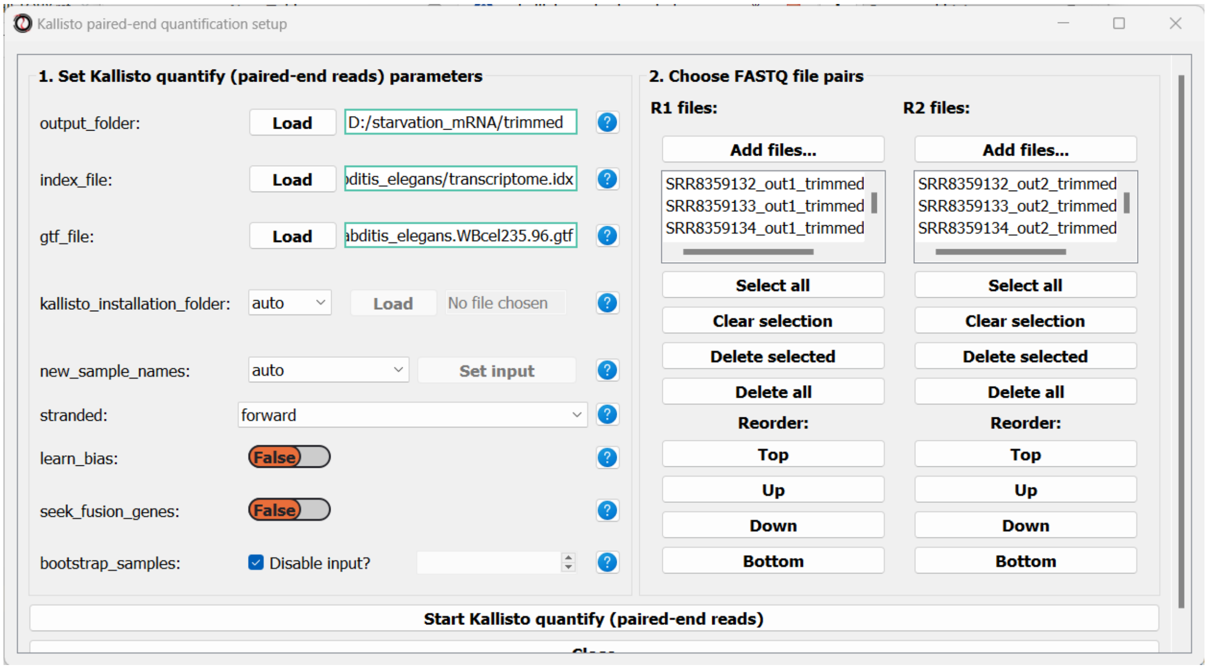

Finally, we can optionally use the *new_sample_names* parameter to give our sample pairs new, more readable names. For example:

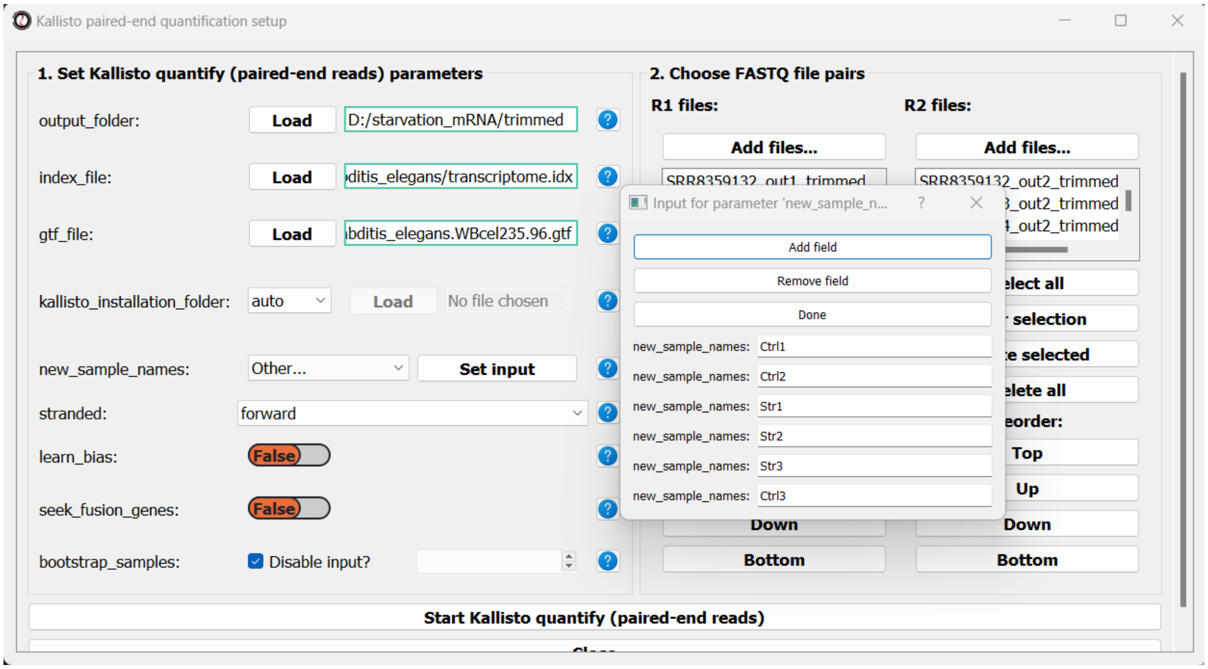

We can now scroll to the bottom of the window and click on the “Start kallisto quantify” button, and wait for the analysis to finish.

Like with the single-end reads analysis, when the analysis is done, we will find the three summarized tables, as well as a subfolder with kallisto output for each pair of FASTQ files.

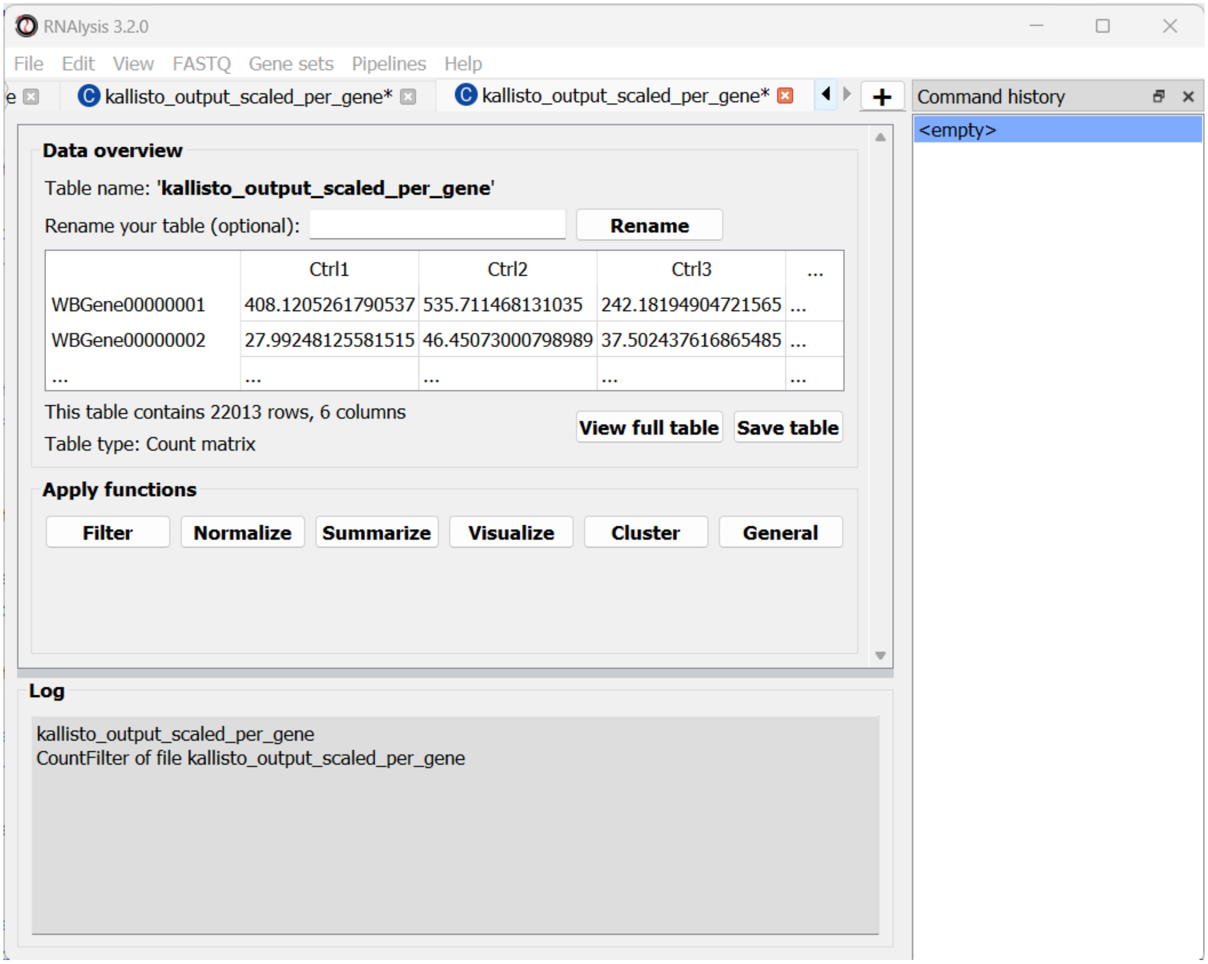


Last but not least, we can quantify our paired-end heat-shock data. The procedure follows the same principle of the starvation samples, with one difference - the heat shock reads are supplied in the reverse orientation, so we should set the *stranded* parameter to 'reverse'. This is what our results should look like:

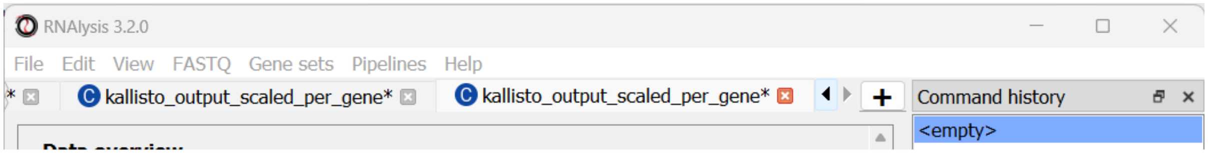


Let's rename the tables to reflect the names of the experiments, and proceed to differential expression analysis.

#### Run differential expression analysis with *DESeq2*

We can now start Differential Expression analysis. For this purpose we will use DESeq2 - an R package for differential expression analysis. Before proceeding with this step, make sure you have installed R on your computer. You don't have to install DESeq2 on your own - *RNAlysis* can install it for you, as long as you have installed the R language on your computer already.

To open the Differential Expression window, choose an *RNAlysis* tab with one of the scaled count tables, click on the "General" tab, and from the drop-down menu below select "Run DESeq2 differential expression":

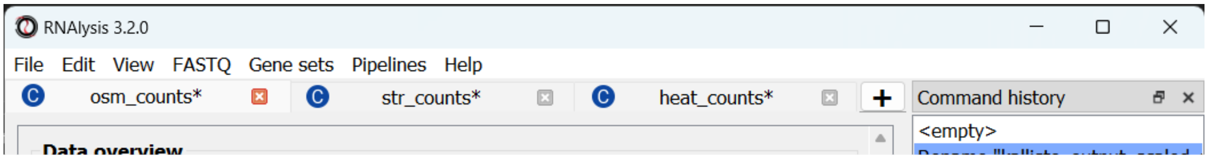


The Differential Expression window should now open. On the left side of the window, set the path of your R installation (or keep it on ‘auto’ if you have previously added R to your computer’s PATH).

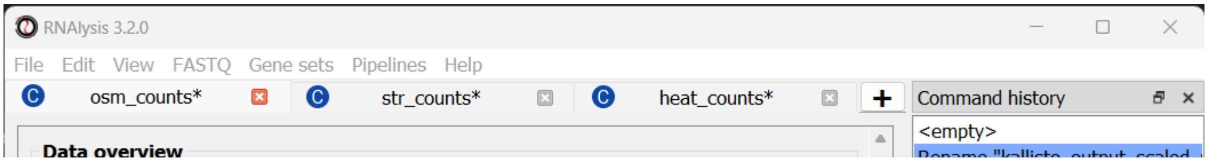


Next, we need to define a **design matrix** for each of our count tables. The first column of the design matrix should contain the names of the samples in the count table. Each other column should contain a variable to be added to the experimental design formula of the dataset. For example: experimental condition, genotype, or biological replicate/batch. For example, the design matrix for our osmotic stress dataset would look like this:

| Name | condition | batch |
|---|---|---|
| Ctrl1 | Ctrl | A |
| Ctrl2 | Ctrl | B |
| Ctrl3 | Ctrl | C |
| Osm1 | Osm | A |
| Osm2 | Osm | B |
| Osm3 | Osm | C |

You can create your design matrix in a program like Microsoft Excel or Google Sheets, and then save it as a CSV or TSV file.

Once you have prepared your design matrix, choose that file from the DESeq2 window and click on the “Load design matrix” button:

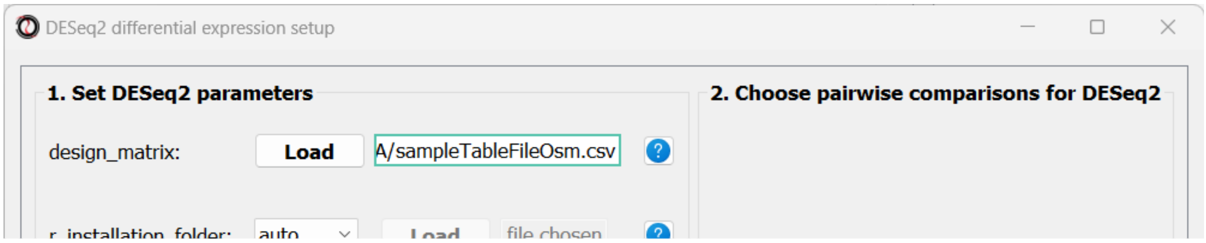


The right side of the window will now update, allowing you to choose which pairwise comparisons you want to run, based on your design matrix. You can make as many pairwise comparisons as you want, each comparing two levels of one of the variables in the design matrix. In our case, we are only interested in one comparison - osmotic stress VS control. Note that the order of conditions in the comparison matters - the first condition will be the numerator in the comparison, and the second condition will be the denominator.

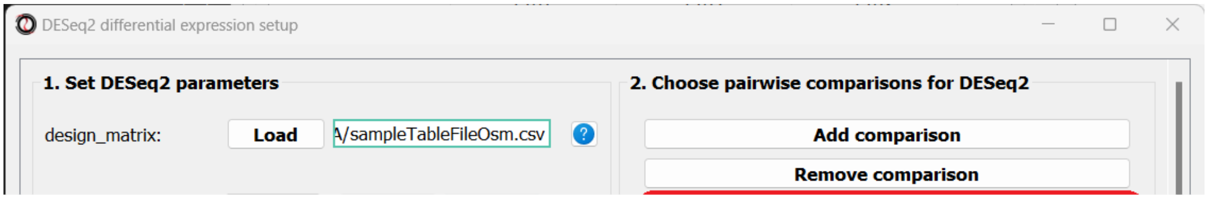


After picking the comparisons you want to run, click on the “Start DESeq2”.

When the analysis ends, a dialog box will pop up, prompting you to choose which differential expression tables do you want to load into *RNAlysis*:

After choosing to load the table, it will open in a new tab in *RNAlysis*:

We can now repeat the same procedure with the other two count tables, and proceed with analyzing the differential expression tables.

#### Data filtering and visualization with customized Pipelines

Since we ran the differential expression analysis through *RNAlysis*, the differential expression tables were loaded into the program automatically. Therefore, we can start analyzing the data straight away!

For each of the differential expression tables we generated, we would like to generate a volcano plot, to filter out genes which are not differentially expressed, and then split the differentially expressed genes into an 'upregulated' group and 'downregulated' group.

We could apply each of those operations to our data tables one by one, but that would take a long time, and there's a decent chance we'll make a mistake along the way. Therefore, we will instead create a customized Pipeline containing all of those functions and their respective parameters, and then apply this Pipeline to all three tables at once.

##### Create a customized Pipeline

To create a Pipeline, let's open the Pipelines menu and click on "New Pipeline":

In the new window that opened, we can name our Pipeline, and choose the type of table we want to apply the Pipeline to. Let's select "differential expression" from the drop-down menu:

After choosing a name and table type, we can click on the "Start" button. The window will now update to show a preview of our new (empty) Pipeline:

At this point, we can start adding functions to the Pipeline. Adding functions to a Pipeline works very similarly to applying functions to table, as we did in Analysis #1. First, let's click on the "Visualize" tab, and choose the "Volcano plot" function:

A volcano plot can give us an overview of the results of a differential expression analysis - it will show us the distribution of adjusted p-values and log2 fold change values for our genes, and highlight the significantly up- and down-regulated genes.

Let's click on the "Add to Pipeline" button:

We can see that the Pipeline overview has updated to show the function we added to it, and this is displayed in the log textbox as well.

Let's now add a filtering function to our Pipeline. Click on the "Filtering" tab and choose the "Filter by statistical significance" function from the drop-down menu:

You may notice that the *inplace* parameter that is usually present for filtering functions is missing. This is because when we apply the Pipeline later on, we can choose whether it will be applied inplace or not.

Finally, we will select the 'Split by log2 fold-change direction' function from the drop-down menu and add it to the Pipeline as well. We can now click on the "Save Pipeline" button to save the Pipeline we created for later use. We can also click on the "Export Pipeline" button to export the Pipeline to a file, so that we can use it in future sessions, or share it with others.

We can now close the Pipeline window and resume our analysis.

#### Apply the Pipeline to our datasets

Now that the hard part is done, we can apply the Pipeline to our differential expression tables. Open the "Pipelines" menu, and then under the "Apply Pipeline" menu, choose the Pipeline we created:

Now, you will be prompted on whether you want to apply this Pipeline inplace or not. Let's choose "no", so that we keep a copy of our original tables.

Finally, you will be prompted to choose the tables to apply your Pipeline to. Let's choose all three of our differential expression tables.

We can now examine the output of our Pipeline - three volcano plots, and six new tables - the significantly up/down-regulated genes from each differential expression table.

At this point, we could optionally rename our tables to make it easier to differentiate them later on.

#### Visualizing and extracting gene set interesctions

We want to know how much do the three stress conditions have in common. One easy way to do this is to visualize the intersections between the differentially expressed genes in the three stress conditions.

#### Create a Venn diagram

Open the “Gene sets” menu, and click on “Visualize Gene Sets…”:

A new window will open. On the left side of the window, we can choose which data tables/gene sets we want to visualize. Let’s pick the three tables that contain significantly downregulated genes:

Next, we can choose the type of graph we want to generate. *RNAlysis* supports Venn Diagrams for 2-3 gene sets, and UpSet plots for any number of gene sets. Let’s choose a Venn Diagram. The window will now update and display various parameters to modify our graph, and a preview of the graph on the right:

We can change these plotting parameters to modify our graph - for example, to change the colors of the Venn circles, the title, or set whether or not our plot will be proportional to the size of the sets and their intersections. Once we are happy, we can click on the “Generate graph” button to create a big version of our graph that we can export and share.

From the looks of it, there is a rather large overlap between the three sets. Let’s generate a similar graph for the tables containing the significantly upregulated genes:

#### Extract the gene subsets we are interested in

Now that we have seen the intersection between the three sets, we want to get the actual list of genes that are significantly downreglated under all stress conditions. To do that, let’s open the “Gene sets” menu once again, and this time click on “Set Operations…”:

A new window will open:

On the left side of the window, we can choose which data tables/gene sets we want to intersect. Let’s pick the three tables that contain significantly downregulated genes:

We will now see a simplified Venn Diagram depicting our three gene sets. We can now proceed to extract the subset we are interested in. We can do this by choosing the “Intersection” set operation from the multiple choice list.

A drop-down menu will now appear, prompting us to pick the primary gene set in this operation. Like filtering operations, some set operations can be applied ‘inplace’, filtering down an existing table instead of returning a new copy of the filtered gene set. For this purpose, *RNAlysis* wants to know what is our “primary set” for this set operation - meaning, if we were to apply it inplace, which table should it be applied to? Our choice will only matter if we apply this set operation inplace - but for our example we won’t be applying the operation inplace, so it doesn’t matter what we choose.

Once we pick a primary set, the subset representing the intersection between all three gene sets will now highlight. *RNAlysis* supports a large number of set operations. You could try using a different set operation on the data - for example, using ‘majority-vote intersection’ to select all genes that were significantly downregulated in *at least* two stress conditions.

Finally, let’s click on the “Apply” button to extract our gene set of interest. It opens in a new tab:

Let’s rename it so we remember what it contains:

Let’s now open the Set Operations window once more to intersect the other set of tables, containing the significantly upregulated genes. This time, let’s use a different method to extract our subset of interest - simply click on the middle subset in the interactive Venn Diagram!

We can use this interactive graph to select exactly the subsets we are interested in, and click the “Apply” button to extract them. After renaming the set, it should look like this:

#### Enrichment analysis

We now have two lists of genes we want to analyze - genes that are significantly downregulated in all three stress conditions, and genes that are significantly upregulated in all three stress conditions. Let’s now proceed to enrichment analysis!

##### Define a background set

In order to run enrichment analysis, we will need an appropriate background set. One good example would be the set of all genes that are not lowly-expressed in at least one sequencing sample. We can use the process we learned earlier to generate an appropriate background set:

1. Create a Pipeline for count tables, that will normalize our count tables and then filter out lowly-expressed genes (for example, normalize with Relative Log Expression, and filter out genes that don't have at least 10 normalized reads in at least one sample)
2. Apply the Pipeline to the three count matrices we generated earlier
3. Use the Set Operation window to calculate the Union of the three filtered count matrices

To keep this tutorial short, we will not go through these individual steps. If you are not sure how to filter count tables for lowly-expressed genes, read Analysis #1 above!

This is what your output should look like:

Now that we have a test set and a background set, we can finally approach enrichment analysis.

#### Define our custom enrichment attributes

Let's assume that we want to specifically find out whether genes downregulated in all stress conditions are enriched for epigenetic genes. Unfortunately, the most common sources for functional gene annotation, Gene Ontology (GO) and KEGG, do not contain a single, well-annotated category of epigenetic genes in *Caenorhabditis elegans* nematodes. Therefore, to be able to test this hypothesis, we will have to source our own annotations.

Luckily, when using *RNAlysis*, you can define your own customized attributes and test enrichment for those attributes. These could be anything - from general categories like "epigenetic genes" and "newly-evolved genes" to highly specific categories like "genes whose codon usage deviates from the norm". The annotations can be either categorial (yes/no - whether a gene belongs to the category or not), or non-categorical (any numerical value - distance to nearest paralog in base-pairs, expression level in a certain tissue, etc).

A legal **Attribute Reference Table** for *RNAlysis* should follow the following format: Every row in the table represents a gene/genomic feature. The first column of the table should contain the gene names/IDs. These should be the same type of gene IDs you use in your data tables. If the gene IDs in your data tables do not match those of your Attribute Reference Table, *RNAlysis* offers a function that can convert gene IDs for your data tables. You can it under the "general" tab.

Every other column in the Attribute Reference Table represents a user-defined attribute/category (in our case, “epigenetic genes”). In each cell, set the value to NaN if the gene in this row is negative for the attribute (in our example - not an epigenetic gene), and any other value if the gene in this row is positive for the attirbute. As an example, see this mock Attribute Reference Table:

| feature_indices | Epigenetic genes | Has paralogs | gut expression |
|---|---|---|---|
| WBGene0000001 | 1 | NaN | 13.7 |
| WBGene0000002 | NaN | 1 | 241 |
| WBGene0000003 | NaN | 1 | 3.6 |
| WBGene0000004 | 1 | 1 | NaN |
| WBGene0000005 | 1 | NaN | 21.5 |

For our analysis, we curated a list of epigenetic genes based on gene lists from the following publications: Houri et al 2014, Dalzell et al 2011, Yigit et al 2006, and based on existing GO annotations. You can find the pre-made curated Attribute Reference Table here.

#### Running enrichment analysis

To open the Enrichent Analysis window, open the ‘Gene sets’ menu and click “Enrichment Analysis”:

We can choose our test and background sets (as explained in Analysis #1) from the two drop-down menus in the Enrichment window:

Next, we can choose the dataset we want to draw annotations from. In our case, we will choose Custom Dataset Categorical Attributes.

After picking the dataset, the window expanded to show us all of the parameters we can modify for our analysis:

Let’s set the statistical test to ‘randomization’, and then set the name of our attributes and the path of our Attribute Reference Table:

Scroll to the bottom of thw window and click on the “run” button to run the analysis:

The enrichment window is going to minimize to allow you to read the log box on the main *RNAlysis* window, but you can enlarge it back if you want to look at the analysis parameters, or start a new analysis with the same parameters.

if you examine the log, you will notice that *RNAlysis* issued a warning about some of the genes in our background set not being annotated. *RNAlysis* will automatically detect and remove from your background set any genes that have no annotations associated with them, since the presence of those genes can make your enrichment results appear inflated.

If the number of genes removed from our background set was large, we may have cause for concern about the validity of your analysis. However, since the percentage of genes removed is very small (123 out of 18924 genes), we can proceed as usual.

Once the analysis is done, we will be able to observe our results in two formats.

The first is a tabular format, showing all of the statistically significant attributes we found (or the all of the tests attributes, if we set the *return_nonsignificant* parameter to True). The table also includes the statistics for each attribute (number of genes in the test set, number of genes matching the attribute, expected number of genes to match the attribute, log2 fold enrichment score, p-value, and adjusted p-value).

The final output format is a bar plot depicting the log2 fold enrichment scores, as well as significance, of the attributes we tested enrichment for.

### Final words

This concludes our A-to-Z tutorial of *RNAlysis*. This tutorial could not possibly encompass every single feature in *RNAlysis*, but at this point you should be familiar enough with *RNAlysis* to start analyzing your own data. If you want a deeper dive into all of the features available in *RNAlysis*, or want to learn how to write code with *RNAlysis*, check out the user guide. If you still have questions about using *RNAlysis*, feel free to drop them in the discussions area.

*RNAlysis* keeps updating all the time, so check out the documentation from time to time to find out what's new.

We hope you found this tutorial helpful, and wish you luck with your future bioinformatic endeavors!
