## Supplementary figures and images for "*RNAlysis*: analyze your RNA sequencing data without writing a single line of code"

### Supplementary Figure 1

Results of K-Medoids Clustering for n_clusters=11, metric='yr1', power_transform=True

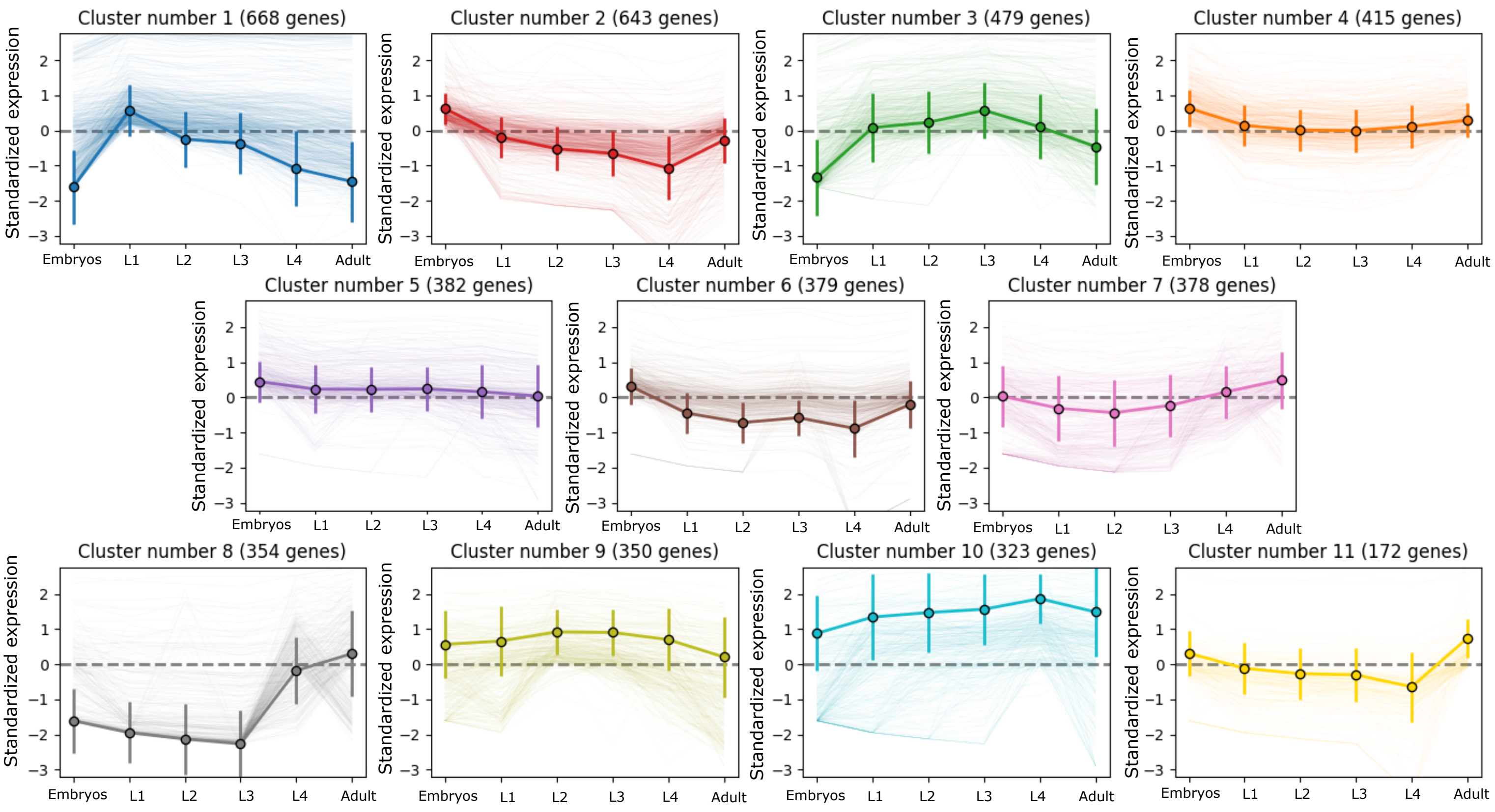
