## Supplementary Figure 2 for "*RNAlysis*: analyze your RNA sequencing data without writing a single line of code"

Results of HDBSCAN Clustering for min_cluster_size=75, min_samples = 1, metric='yr1', epsilon=0.0, method='eom' and power_transform=True

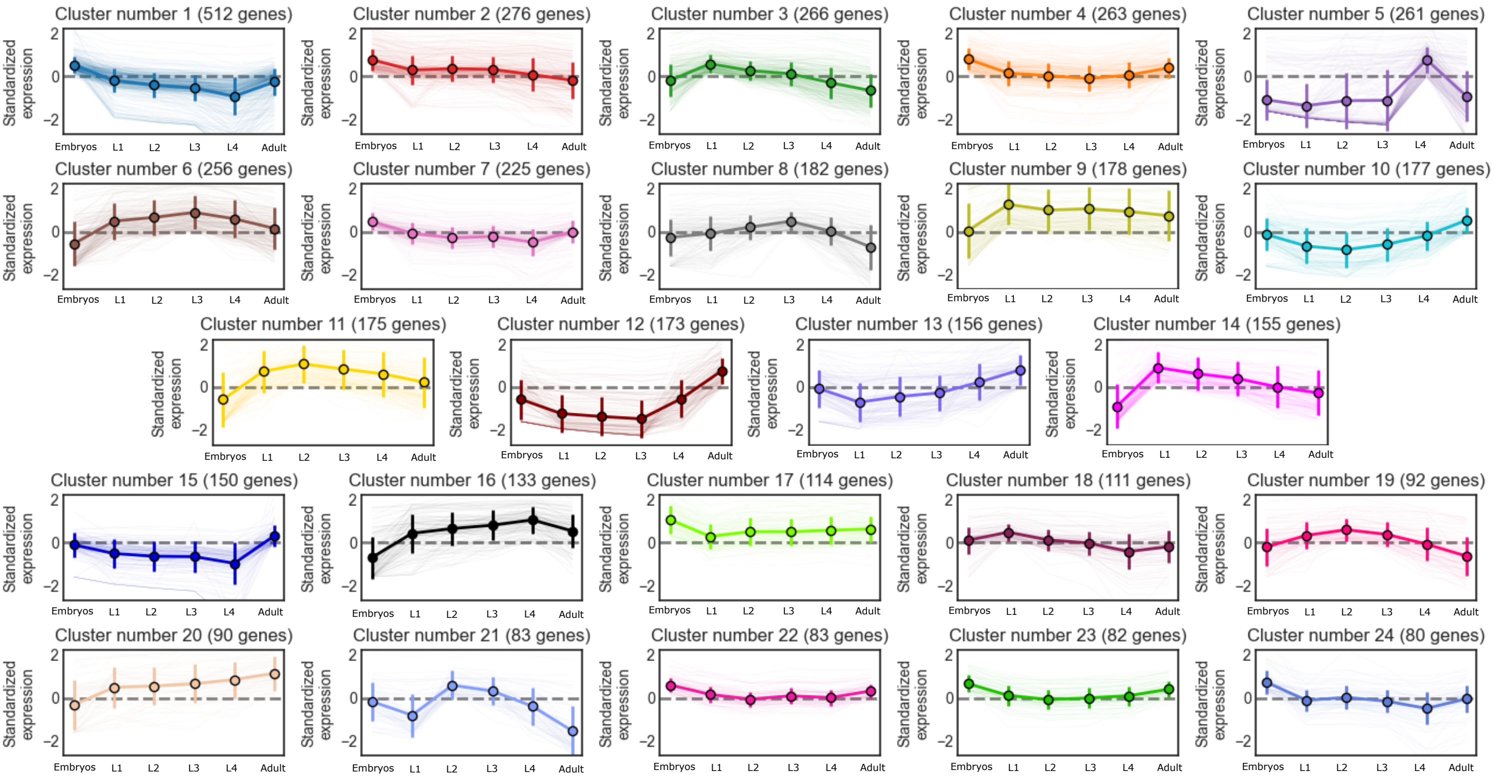
